# Single-Cell Profiling Reveals Innate Lymphoid Cells and OX40 Activation in Breast Cancer-Related Lymphedema

**DOI:** 10.64898/2026.09.09.749825

**Authors:** Gabriela Martinez-Chacón, Martin Rosenborg, Heli Jokela, Susanna Pajula, Heidi Gerke, Damien Kaukonen, Katri Orte, Emilia Peuhu, Marko Salmi, Pia Rantakari, Pauliina Hartiala

## Abstract

Breast cancer-related lymphedema (BCRL) is a severe complication affecting up to 40% of breast cancer survivors. Stage II or chronic disease stages are characterized by upper-limb edema, fibrotic adipose tissue accumulation, pain, and recurrent infections. Despite its clinical burden, BCRL lacks effective pharmacological therapies and the molecular mechanisms driving disease progression remain poorly understood. To better define the regulatory mechanisms underlying chronic BCRL, we performed transcriptomic and quantitative analyses of stromal vascular fraction cells (SVF or non-adipose cell fraction) in BCRL-derived subcutaneous adipose tissue. CD45^+^ immune cells from stage II BCRL were compared with those from healthy lean and obese adipose tissue controls. Although our analyses identified multiple leukocyte populations, we focused primarily on innate lymphoid cells (ILCs), particularly ILC2 and ILC3 subsets, which were more abundant and active in BCRL tissues. Bioinformatics analysis of cell-cell communication positioned ILCs as central immune-regulatory hubs in BCRL through inflammatory and tissue-remodeling pathways, including connections to IL2, OX40, KIT, and LT signaling. Strong bidirectional communication between ILCs and regulatory T cells via OX40 signaling was uniquely detected in BCRL. Consistent with these findings, protein-based analyses confirmed increased inflammatory mediators and enrichment of the OX40 and OX40L co-stimulatory pair in BCRL tissues. Taken together, our findings identify OX40-dependent ILC-signaling as a novel regulatory program supporting chronic inflammation and pathological tissue remodeling in BCRL.

## INTRODUCTION

Breast cancer is the most common cancer among women worldwide (1). Treatment of metastatic breast cancer requires surgical treatment and radiation therapy of the axillary lymph nodes, disrupting lymphatic flow in the closest arm. This may lead to breast cancer-related lymphedema (BCRL), one of the most common long-term complications of breast cancer treatment (2). The cumulative incidence of BCRL is up to 40% following axillary lymph node dissection, and symptoms may develop months or even years after the initial operation.

Lymphedema is a chronic disorder with a substantial global health burden. BCRL is one of the most common forms of secondary lymphedema, but it may also develop after treatment for other malignancies, including gynecological, urogenital and skin cancers, as well as following trauma and recurrent infections (3,4). Regardless of its cause, lymphedema typically progresses from an early stage characterized predominantly by reversible interstitial fluid accumulation (stage I) to a chronic stage (stage II) marked by persistent swelling, fibrosis and pathological adipose tissue accumulation. In patients with chronic BCRL, the excess fibrotic adipose tissue can reach several liters, leading to pain, heaviness, impaired arm function and recurrent soft tissue infections that often require prophylactic antibiotic treatment (5,6).

BCRL is treated conservatively with compression garments, manual lymphatic drainage, and exercises to improve lymphatic drainage (3). During recent years, reconstructive surgical procedures, such as microvascular lymph node transfer and lymphaticovenous anastomosis (LVA), have become more frequent (7,8). However, these procedures are beneficial to less than 40% of patients (9). Currently, one of the most effective procedures for patients with chronic lymphedema remains liposuction, which reduces arm volume and relieves the symptoms of lymphedema, although it does not improve lymphatic flow (6).

Chronic inflammation, fibrosis, and adipose hypertrophy are well-established histological features of lymphedema (10). In particular, the expansion and activation of the CD4^+^ Th2 cell population in lymphedema has been shown to promote inflammation and fibrosis via IL4 and IL13 production (11). Increased infiltration of T regulatory cells (Tregs) and dendritic cells (DCs) has also been observed in affected tissues. Tregs have been demonstrated to modulate chronic inflammation through local immunosuppressive mechanisms (12). Circulating DCs have been shown to migrate to lymphedematous tissue, become activated through damage-associated molecular pattern (DAMP) signals, and migrate to regional lymph nodes to interact with T cells (13). Immunomodulatory drugs, such as tacrolimus, a calcineurin inhibitor (14) and IL4/13 neutralizing antibodies (15) targeting these events have been tested in several clinical trials for patients with lymphedema. However, none have effectively reduced the excess fluid or fibrotic adipose tissue in the lymphedema arm (14–18). Despite extensive research, there are still no definitive cures or effective medical treatments for lymphedema. This is at least in part due to the fact that animal models only partially reflect the clinical pathophysiology and treatment history of lymphedema patients who have often undergone aggressive combinations of radical surgery, radiation therapy, chemotherapy, and hormonal therapy.

To better understand the pathophysiology of BCRL in humans, we analyzed fresh tissue samples obtained from patients with chronic BCRL undergoing liposuction treatment. Using single-cell RNA sequencing (scRNA-seq), we characterized the gene expression profiles of CD45⁺ immune cells of the subcutaneous adipose tissue and compared them with those from healthy individuals who were either lean or obese. Our analysis revealed an increased abundance of innate lymphoid cells (ILCs) in BCRL tissue, which was further confirmed by flow cytometry. ILC2 and ILC3 cells are tissue-resident immune cell populations involved in barrier integrity, tissue repair, and immune homeostasis, and have been shown to contribute to dysregulated Th2 and Th17 immune responses in several other chronic inflammatory diseases (19,20). Bioinformatics analysis of cell-cell communication revealed a highly active immune network in which ILCs act as major emitters and receivers of IL2, KIT, LT, and OX40 signaling among other immune cells, including T, NK, and Treg cells. Elevated protein levels of MIF, TNFα, and IL1RA were detected in BCRL tissue lysates, demonstrating chronic tissue inflammation and remodeling. We also demonstrate an increased expression of the OX40L/OX40 co-stimulatory pathway in BCRL at the protein level using comprehensive histological analyses. Our study provides a broad characterization of the immune microenvironment in human BCRL and identifies novel immune pathways that are likely to contribute to persistent inflammation and pathological tissue remodeling.

## RESULTS

### Characterizing the cellular diversity and expression signatures in stage II BCRL using scRNA-seq

Average age and BMI of the stage II BCRL patients (n=4) were 57,5 ± 2,5 years and 29,2 ± 1,5 kg/m^2^, respectively (Table 1). As a control non-BCRL group, we used the scRNA-seq dataset from a study that compared lean (LSAT) and obese (OSAT) subcutaneous adipose tissue (GSE155960) (21). This dataset included individual samples from 3 lean and 3 obese women (Supplementary table 1) with an average age of 38 ± 3,2 years (LSAT) and 36,6 ± 1.2 years (OSAT), and a BMI of 20,7 ± 2,2 kg/m^2^ (LSAT) and 31,67 ± 0,9 kg/m^2^ (OSAT). A schematic workflow of the study is provided in Figure 1.

**Figure 1.**
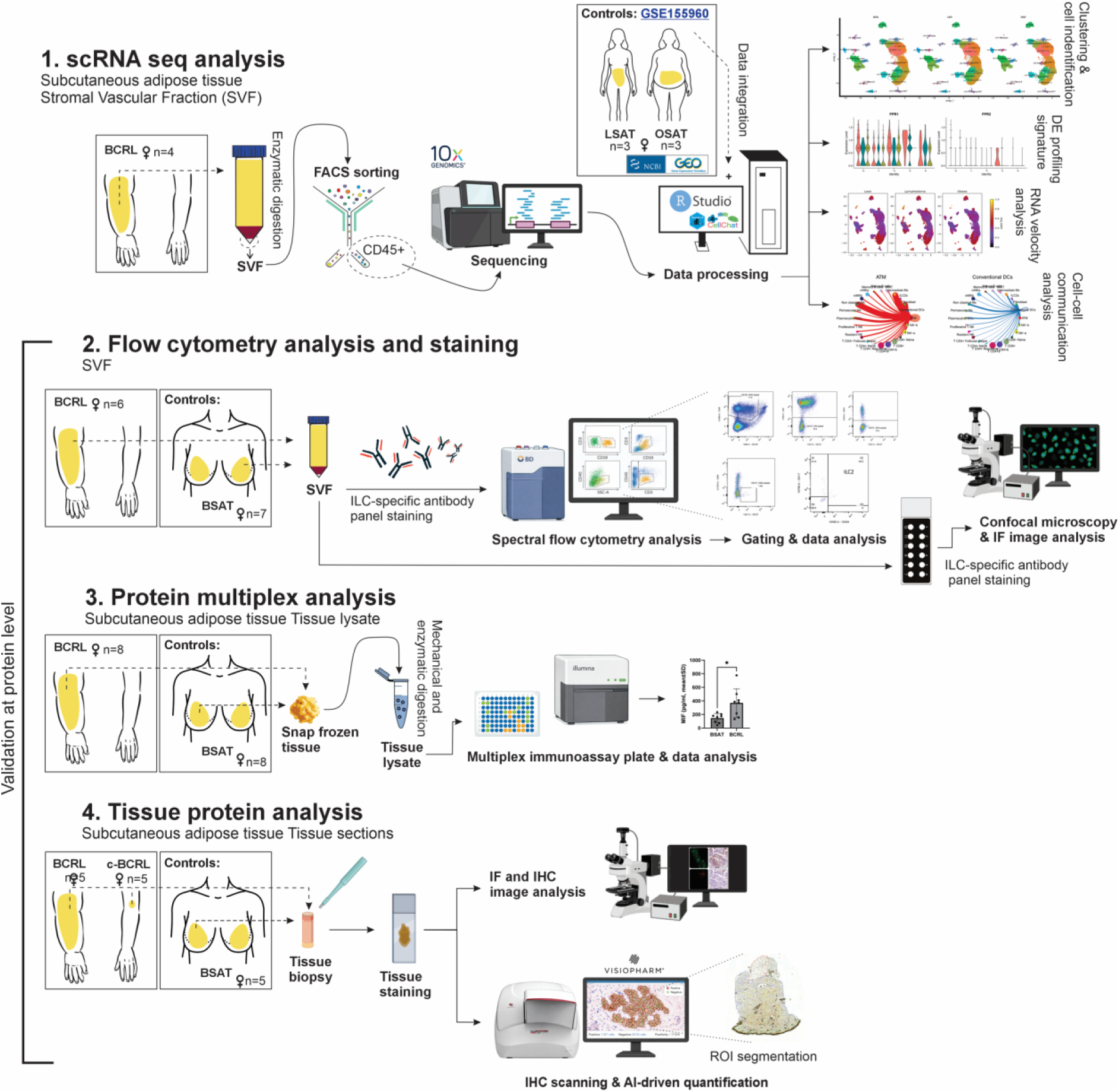
Schematics of the Experimental Worflow. BCRL lipoaspirate was collected from patients during surgical treaments. Isolated stromal vascular fraction (SVF) was used for scRNA-seq, spectral flow cytometry, and immunofluorescence staining. Tissue samples were used for IHC staining and protein multiplex assay.

The resulting integrated dataset included BCRL_1–4: 20.521 cells, LSAT_1–3: 24.877 cells and OSAT_1–3: 19.641 cells, for a total of 65.039 high-quality cells (Supplementary figure 1A and 1B). After data integration, cell populations were revealed by unsupervised clustering and projected on a UMAP map (Figure 2A). Using a combination of known markers and the specific genes expressed in each cluster (Supplementary table 2), we identified 24 clusters (c0–c23, Figure 2A). The top 3 DE or signature genes from each of the clusters are presented in Figure 2B. Clusters were further classified into specific cell populations using hierarchical clustering and established lineage-specific markers (Figures 2C and 2D, respectively). Detailed transcriptional signatures used for the annotation of all 24 clusters are presented in Supplementary Information 1 and Supplementary table 2.

**Figure 2.**
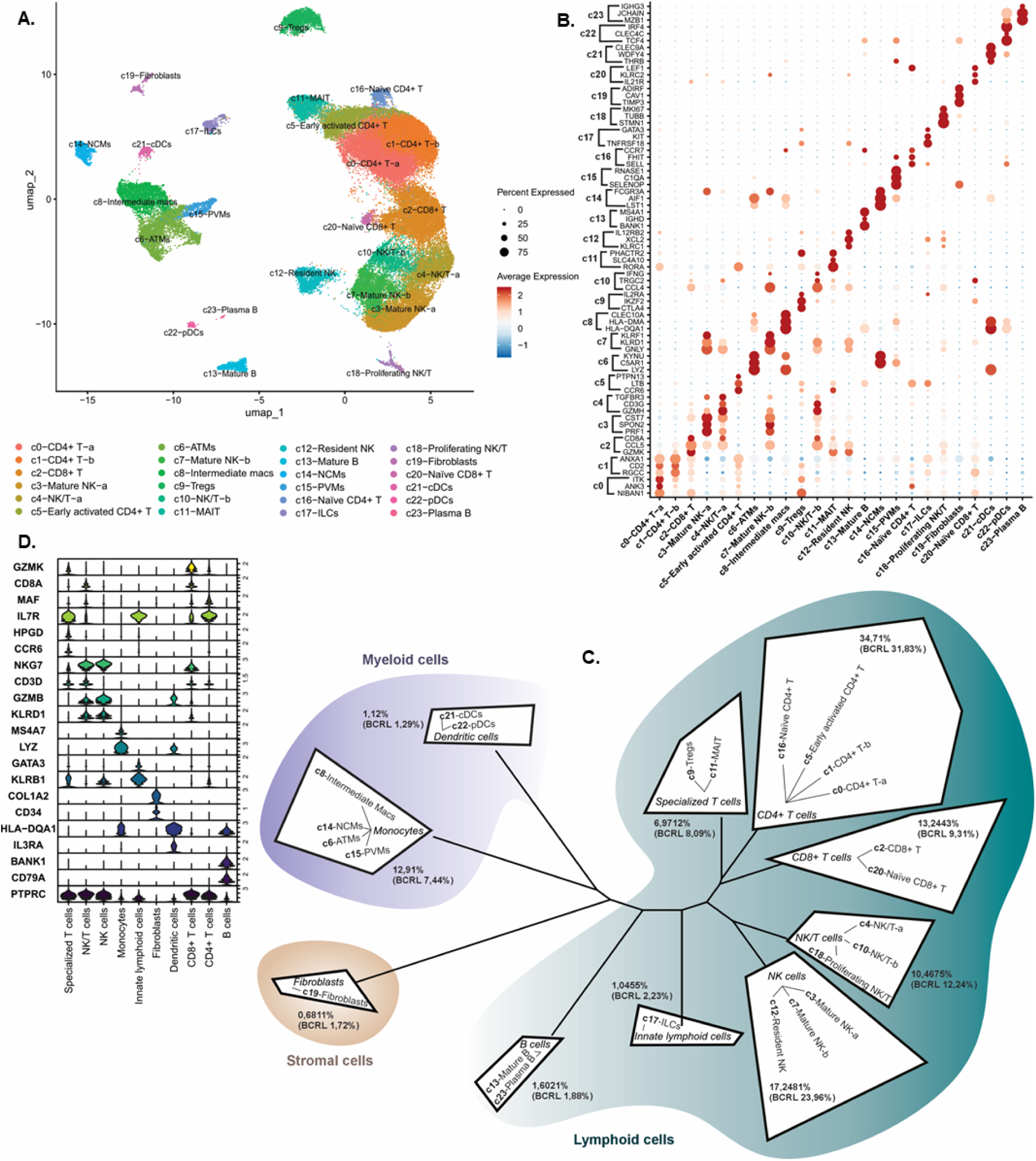
Single-cell transcriptomic landscape of immune populations in BCRL and control lean andsubcutaneous obese adipose tissue. **A.** UMAP visualization showing 24 immune cell clusters identified from the integrated scRNA-seq analysis of BCRL, LSAT and OSAT adipose tissue samples**. B.** Dot plot displaying selected top differentially expressed genes across identified populations. Color intensity represents the average expression level in expressing cells, and dot size indicates the percentage of cells within each cluster expressing the gene. **C**. Hierarchical clustering of major immune cell types based on the mean expression of the most variable genes in the integrated dataset. **D.** Expression patterns of established lineage and major cell-type marker genes used to annotate each cluster.

The frequencies (percentages) of each identified cell population within the total CD45^+^ population were quantified in the integrated dataset as well as in the BCRL samples alone (Figure 2C). In total, 10 different cell group populations were identified: CD4^+^ T cells, NK cells, CD8^+^ T cells, Monocytes, NK/T cells, Specialized T cells, B cells, Dendritic cells, Innate lymphoid cells and fibroblasts, within the three main cell lineages present in the SVF: lymphoid, myeloid and stromal cells (Figures 2C and 2D).

Among these, **CD4^+^ T cells** (marked by *CD3D*, *IL7R* and *MAF* genes) constituted the largest group (34,71%) of the total CD45^+^ (*PTPRC*) cell population. Other larger immune cell types identified were (Figures 2A–D), **natural killer cells** (**NK** 17,24%; marked by *KLRD1* and *PRF1* genes and lack of *CD3* expression (22)) including cells in c3 (Mature NK-a, mNK-a), c7 (Mature NK-b, mNK-b) and c12 (Resident NK); **CD8^+^ T cells** (13,24%; marked by *CD8A* and *GZMK* (23)) including cells in c2 (CD8+ T) and c20 (Naïve CD8 T); **Monocytes** / macrophages (12,90%; marked by *LYZ* and *MS4A7* (24)) including cells in c6 (Adipose Tissue Macrophages, ATMs), C8 (Intermediate Macrophages, macs), c14 (Nonclassical Monocytes, NCMs) and c15 (Perivascular Monocytes, PVMs); **NK/T cells** (10,46%; marked by *NKG7*, *CD3D*, *KLRD1 and PRF1* genes (25)) including cells in c4 (NK/T-a), c10 (NK/T-b) and c18 (Proliferative NK/T) and **Specialized T cells** (6,97%; marked by *CCR6* and *HPGD* (*26*)) including cells in c9 (CD4^+^ T regulatory cells, Tregs) and c11 (Mucosal Associated Invariant T, MAITs).

In addition to the above described clusters, we also identified less abundant cell populations, including **B cells** (1,60%; marked by *CD79A* and *BANK1* (27)) in c13 (Mature B) and c23 (Plasma B); **Dendritic cells** (DCs, 1,12%; marked by *IL3RA* and *HLA-DQA1* (28)) in c21 (Conventional, cDCs) and c22 (Plasmacytoid, pDCs); **Innate Lymphoid Cells** (ILCs 1,04%; marked by *KLRB1* and GATA3 and the lack of *CD3* and *KLRD1* (*22*)) in c17 ; and **Fibroblasts** (0,68%; marked by *ADIRF* and *COL1A2* (29)) in c19 (Figures 2A–D).

### BCRL is characterized by shifts in immune cell profiles and enrichment of activation and co-stimulatory gene signatures

To identify shared and unique immune cell features between BCRL and control LSAT and OSAT SVFs, we compared the average proportion of cells within each cluster across conditions. Importantly, cells from all three conditions were represented across all 24 clusters, supporting successful dataset integration based on shared lineage signatures rather than sample-specific features (Figure 3A and Supplementary figure 1A-B).

**Figure 3.**
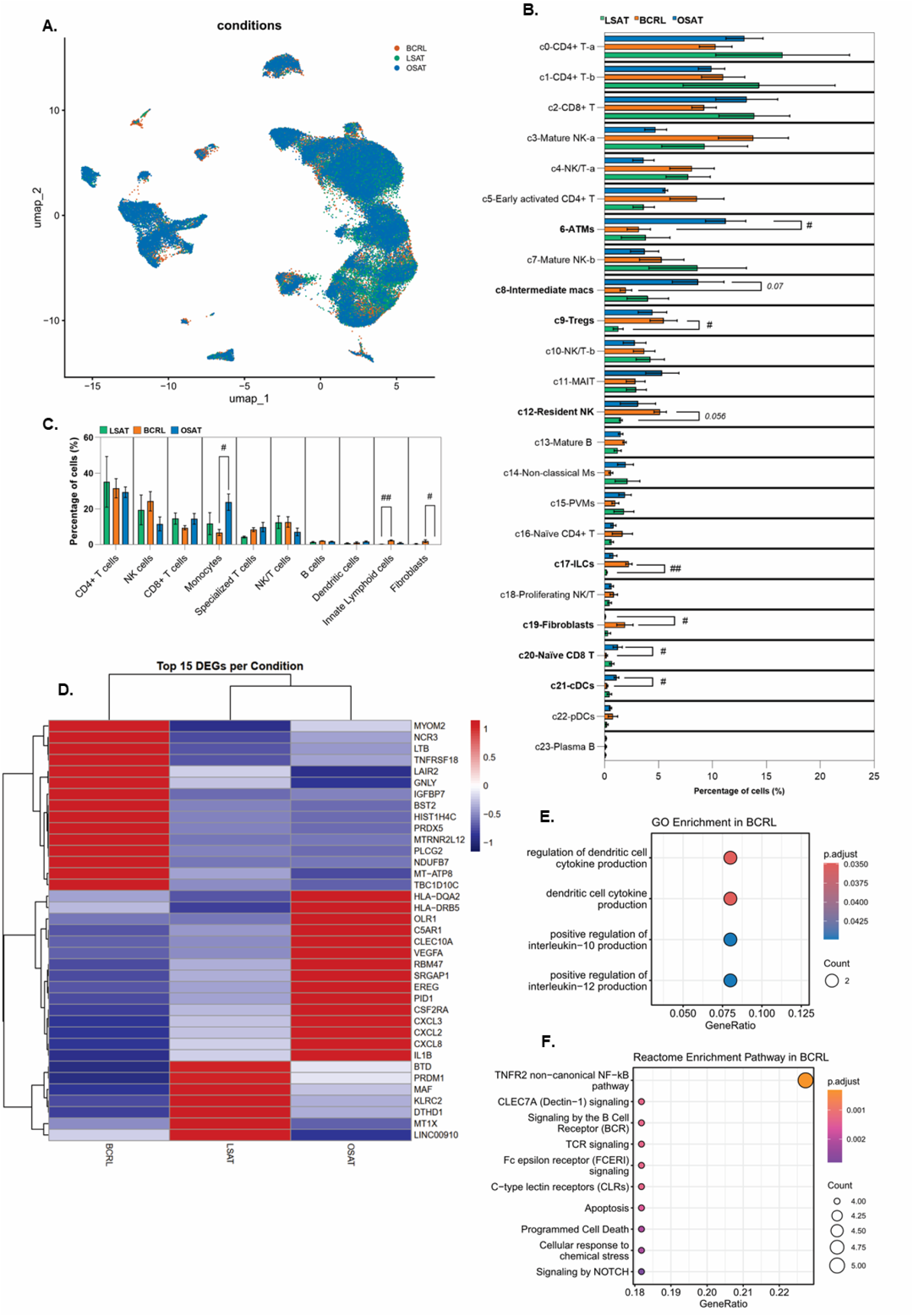
Integrated scRNA-seq analysis reveals immune cell diversity and enrichment of ILCs in BCRL-affected adipose tissue. **A.** UMAP visualization showing the distribution of 20.521 cells from BCRL and 19.555 and 19.641 cells from control LSAT and OSAT samples across identified immune cell clusters. **B.** Bar plot showing the relative frequency of cells derived from BCRL, LSAT and OSAT samples within each cellular cluster. **C.** Frequency of cells across conditions grouped by lineage or major immune cell type. For B and C, statistical analysis performed using the Kruskal–Wallis test, followed by Dunn’s post hoc test (*p≤* 0,05). **D.** Heatmap displaying the top 15 differentially expressed genes (DEGs) across BCRL, LSAT and OSAT conditions. **E.** Gene Ontology (GO) enrichment analysis of top 30 DEGs from BCRL highlighting enriched biological (immune related) processes. **F.** Reactome pathway enrichment analysis of top 30 upregulated genes in BCRL compared to control conditions. Dot size represents the number of genes mapped to each pathway, and color indicates the adjusted p-value (Benjamini-Hochberg correction). Only significantly enriched pathways are shown.

No major differences were observed in the overall frequencies of CD4^+^ T cells, NK/T cells, dendritic cells, B cells, or CD8^+^ T cells across conditions (Figure 3B). However, cluster-level analysis identified a trend toward increased resident NK cells in BCRL tissue *(p= 0,056),* and a reduced frequency of naïve CD8^+^ T cells *(p=* 0,05) and conventional dendritic cells compared *(p=* 0,05) with control conditions (Figure 3C). No significant differences were detected in mature or plasma B-cell populations.

When comparing BCRL and control SVF samples, macrophages, Tregs, ILCs, and fibroblasts showed the most pronounced changes in cell frequencies (Figures 3B and 3C). Within the monocyte group, BCRL showed a significant reduction (Figures 3B and 3C), particularly in macrophages compared to OSAT (p = 0,05; BCRL 6,72% ± 3,47; OSAT 23,74% ± 7,98). Resident ATMs (BCRL 3,18% ± 2,16; OSAT 11,23% ± 3,23) and Intermediate macrophages (BCRL 1,972% ± 1,08; LSAT 4,02% ± 3,347; OSAT 8,674% ± 4,13) showed the larger reduction. In contrast, Tregs (c9) from the specialized T cell group showed a significant expansion (*p =* 0,046) in BCRL (Figure 3C), specifically compared to the LSAT control samples (BCRL 5,478% ± 2,511; LSAT 1,264% ± 0,789; OSAT 4,432% ± 2,285).

In addition, our study revealed a previously unreported expansion of ILCs in BCRL samples (*p* = 0,009). These cells appeared to represent a distinct feature of the disease, as ILCs were rarely detected in the SVF of control tissues (BCRL 2,25% ± 0,58; LSAT 0,22% ± 0,06; OSAT 0,81% ± 0,56) (Figures 3B and 3C). We also verified that the increased frequency of ILCs was consistently exhibited in all the BCRL samples (Supplementary figure 1A). Similarly, we found that 80% of all fibroblasts in c19 derived almost exclusively from the BCRL samples (BCRL 1,88% ± 1,51; LSAT 0,32% ± 0.42; OSAT 0,08% ± 0,01) (Figures 3A and 3B). However, due to their limited quantity in the BCRL SFVs and very low expression of *PTPRC*, we did not analyze them further in this study.

To visualize gene expression patterns unique in BCRL, we compared cells from lymphedema with those from the other two conditions and identified significantly upregulated genes. The top 15 DE genes for each condition were selected based on the highest average log₂ fold change (Figure 3D). BCRL samples displayed a distinct gene expression profile that clearly differentiates this pathological condition from both OSAT and LSAT. There was an upregulation of cytotoxic and pro-inflammatory genes, including *NCR3* (30) (NKp30), *GNLY, PLCG2* (31), and *BST2* (32), suggesting robust activation of NK and cytotoxic T cells in BCRL tissue. In turn, elevated expression of the immunoregulatory gene *TNFRSF18* may indicate engagement of Th2-related cells in the BCRL tissue microenvironment (33). Tissue remodeling markers such as *IGFBP7* (34) and *LTB* (35,36) were found enriched in BCRL tissues, particularly in fibroblasts, leukocytes and B cells, potentially reflecting local tissue reorganization and lymphangiogenic responses. One of the most prominent transcriptional changes observed in BCRL is the upregulation of *LTB* across multiple immune populations, including B cells, DCs, ILCs and T cell subsets, indicating a systemic response rather than a cell-type-specific gene signature.

To pinpoint biological processes or pathways that are potentially involved in BCRL, we performed Gene Ontology (GO) and Reactome analysis. GO analysis revealed enrichment of processes related to DC cell cytokine production (Figure 3E), including regulation of IL10 and IL12, suggesting activation of antigen-presenting cell functions as previously described (37). Reactome analysis identified enrichment of TNFR2 signaling together with multiple immune-related pathways, including B cell receptor (BCR), T cell receptor (TCR), and C-type lectin receptor signaling (Figure 3F), suggesting broad activation of both innate and adaptive immune responses (38,39). Collectively, the enrichment of these pathways indicates that cytokine-mediated interactions and co-stimulatory signaling may converge to drive chronic inflammation in BCRL.

### Gene signatures identify diverse ILC populations and activated immune states in BCRL

Because cluster c17 (ILCs) was predominantly composed of cells from BCRL tissues, we next performed differential expression analysis to define its transcriptional signature. BCRL-associated ILCs lacked expression of CD3, CD8, CD14, CD16, CD20, FCER2 and KLRD1, distinguishing them from T cells, B cells, basophils and NK cells (Figure 4A, left panel). They expressed canonical ILC markers, including KLRB1 and IL7R, whereas the ILC1-associated transcription factors TBX21 (T-bet) and EOMES were minimally expressed (Figure 4A, middle panel). Instead, these cells exhibited a mixed ILC2/ILC3-like phenotype characterized by expression of the ILC2-associated markers GATA3, RORA and AREG (40), together with KIT, a marker shared by ILC2s and ILC3s, and IL1R1, AHR and AFF3, which are characteristic of ILC3s (21,41,42) (Figure 4A). Feature plots further highlighted the robust and uniform expression of AREG, whereas PTGDR2 (CRTH2), a marker of mature ILC2s (40), was only weakly expressed (Figure 4B). High expression of *AREG* (amphiregulin), an EGF receptor ligand, have been associated with processes involving tissue repair, fibrosis, and immune regulation (4,43). In addition, BCRL-associated ILCs prominently expressed TNFRSF18 (GITR) and TNFRSF4 (OX40), members of the TNF receptor superfamily associated with immune activation and Th2-related inflammatory responses (44), together with SPINK2, suggesting a dynamic ILC differentiation and activation state (45). Finally, these cells uniquely expressed the ILC3-associated factors LTB, XCL1 and XCL2, molecules implicated in immune-cell recruitment, chronic inflammation, lymphoid tissue organization and fibrosis (36,46) (Figure 4A and 4B).

**Figure 4.**
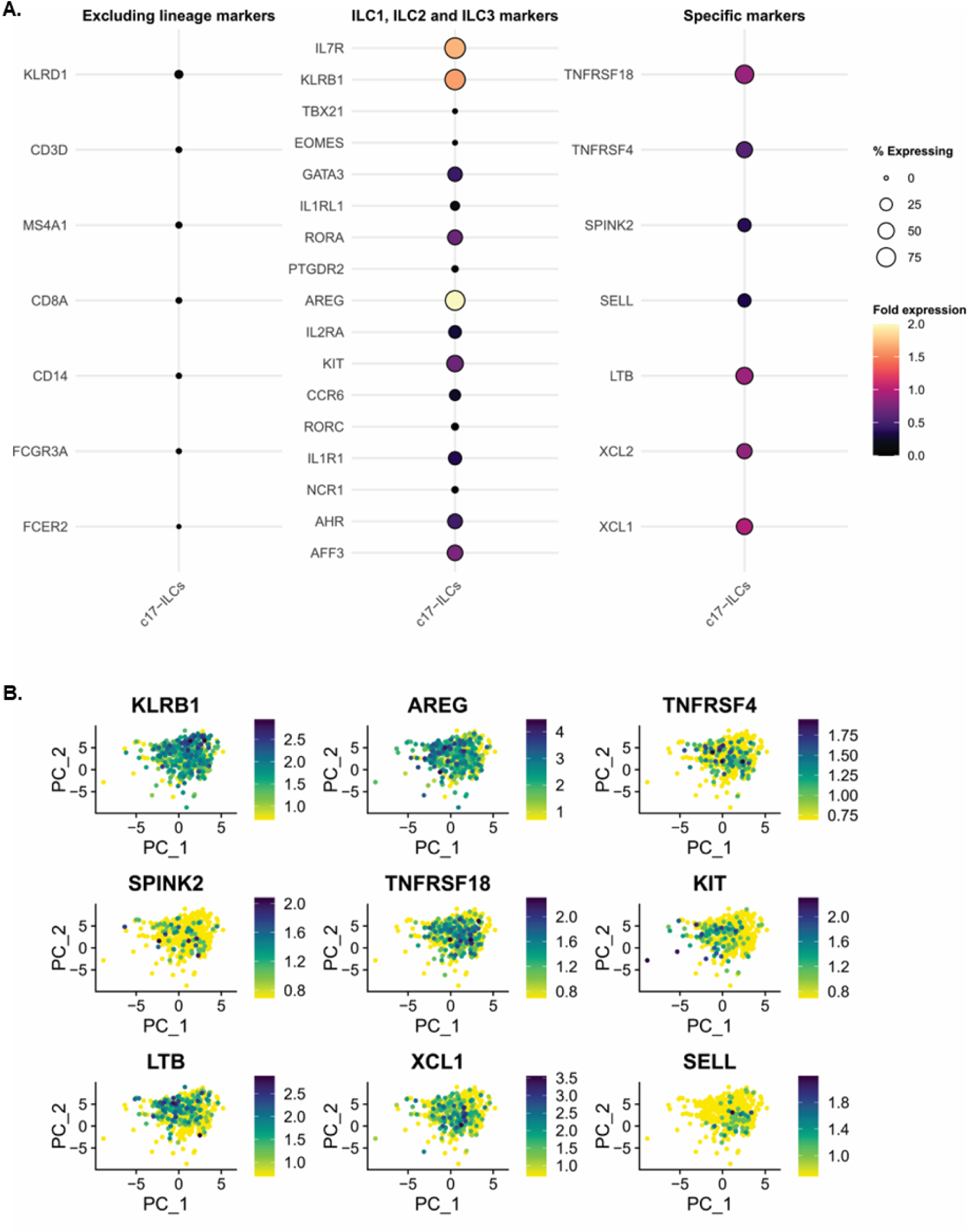
BCRL-associated ILCs exhibit a mixed ILC2- and ILC3-like transcriptional profile. **A.** Dot plot showing the expression of canonical ILC marker genes together with key markers enriched in the ILCs identified in the BCRL dataset. Dot size indicates the percentage of expressing cells and color intensity reflects average expression levels. **B.** Feature plots showing the distribution of ILC2- and ILC3-associated marker expression across the subsetted BCRL-associated ILCs. Color gradient represents normalized gene expression levels, with lighter colors indicating low expression and darker colors indicating higher expression.

### BCRL associates with activated transcriptional states in Tregs and dendritic cells and remodels immune-cell dynamics

As the amount of Tregs was also increased in BCRL and given their central role in modulating immune responses, we next investigated the transcriptional profile of this population compared to control conditions. DE gene analysis revealed an activated and co-stimulatory phenotype, characterized by increased expression of *TNFRSF4* and *TNFRSF18* (Supplementary figure 2A). In addition, Tregs expressed LTB and other T cell activation and pro-inflammatory genes, including *LIME1, PRDX1, TUBB4B* and *IRF1*.

As DE gene enrichment analysis indicated altered DC-related processes in BCRL, we examined the transcriptional profile of DC clusters c21 and c22. BCRL-associated DCs displayed an activated plasmacytoid-like phenotype characterized by high expression of interferons and effector genes, including *CLEC4C*, *GZMB*, *PLAC8*, *CTSC* and *LILRA4* (Supplementary figure 2B) (32). In addition, these cells displayed increased expression of lipid-associated genes such as *PTGDS* and *PLAAT3*, suggesting a potential metabolic adaptation to the BCRL fatty tissue microenvironment (Supplementary figure 2B) (47).

To investigate whether BCRL transcriptional signatures alter immune dynamics, we performed scVelo RNA velocity analysis across all conditions (Supplementary figure. 3A). Comparative latent-time distributions revealed substantial remodeling of immune states in BCRL. Specifically, DCs, B cells, ILCs and monocytes displayed increased latent-time values relative to LSAT and OSAT tissues (Supplementary figure. 3B, Supplementary table 3), suggesting progression toward chronic, inflammation-associated activated states (48,49). Consistent with these findings, BCRL tissues exhibited broadly reduced trajectory coherence and significantly decelerated differentiation velocity length across key cell clusters (Supplementary table 3). Monocytes, B cells, specialized T cells, CD4^+^ T cells and NK/T cells showed the most profound drops in velocity confidence and speed relative to the LSAT reference (*P. adj* < 0,001) for all comparisons; Supplementary table 3). This systemic loss of directionality and coherence indicates increased transcriptional heterogeneity and less coordinated cell-state transitions (50). Collectively, these findings demonstrate widespread perturbations in immune dynamic states within BCRL tissues, particularly impacting antigen-presenting and innate immune populations.

### Increased ILC and T cell intercellular communication in BCRL favors immune activation and tissue remodeling pathways

By comparing outgoing and incoming interaction strengths, we identified cell populations with significant changes in their signaling roles between BCRL (Figure 5A, right-plot) and each control condition, LSAT and OSAT (Figure 5A, left and central plots). Interestingly, major changes in communication roles occurred also in those cell populations that showed frequency shifts in BCRL. NK/T-a, naïve CD8^+^ T, ILC and Treg populations exhibited increased signaling activity in BCRL compared with LSAT and OSAT (Figure 5A, arrows pointing up). While, monocytes, macrophages and cDCs, displayed reduced signaling activity and effectiveness in interactions with other immune cells (Figure 5A, arrows pointing down).

**Figure 5.**
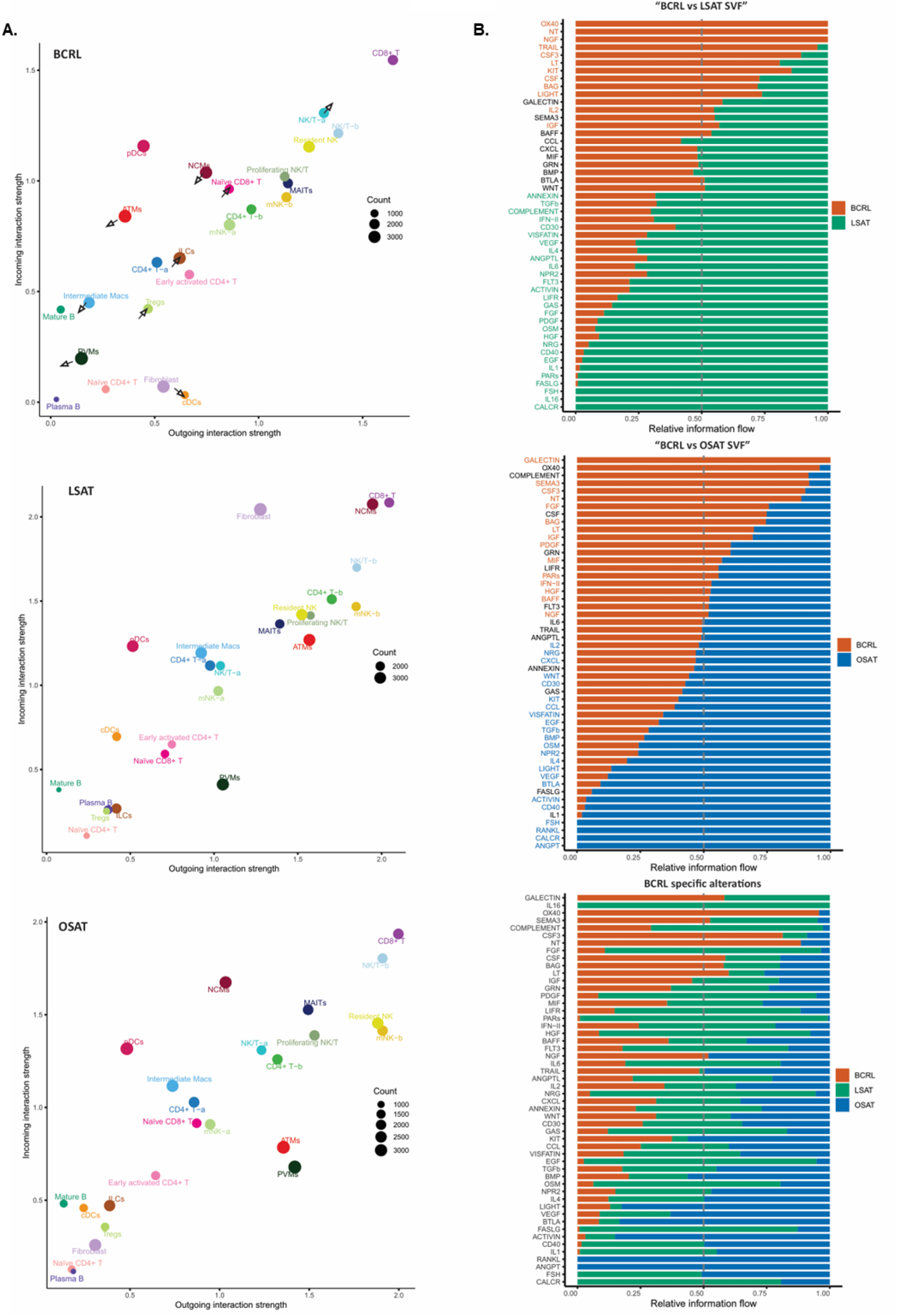
Cell-cell communication analysis identifies signaling patterns associated with Th2 inflammation and tissue remodeling in BCRL. **A.** Network plot illustrating intercellular communication dynamics among identified immune cell types across BCRL, LSAT and OSAT conditions. Arrows indicate the directionality of signaling, highlighting changes in sender and receiver activity between conditions **B.** Relative information flow analysis. Left and middle panels show pairwise comparisons of signaling flow between BCRL and LSAT or OSAT conditions, respectively. The right panel compares signaling flow across all three conditions. Orange labels indicate pathways significantly enriched in BCRL (Wilcoxon test)

Comparative signaling analysis identified several pathways uniquely enriched or increased in BCRL compared to control LSAT and OSAT conditions, including NT, NGF, CSF3, BAG, LT and IGF (Figure 5B). OX40 pathway also showed greater inferred communication strength in in BCRL compared with both control conditions, but it was only statistically significant when comparing that to LSAT (BCRL vs OSAT *p =* 0.125). Cell-cell communication analysis further revealed robust intercellular signaling activity of ILCs in BCRL, particularly involving interactions with pDCs, Tregs, naïve CD8^+^ T cells, ATMs, cytotoxic NK/T populations, fibroblasts and cDCs (Figure 6A). Among the major signaling contributors in BCRL, ILCs showed strong involvement in CXCL, MIF, EGF, NGF, ANNEXIN, BAG and WNT pathways, primarily targeting T cells, monocytes, pDCs, fibroblasts and mNK cells (Figure 6B). In addition, ILCs were actively involved in Th2- and Th17-associated signaling pathways, including LT, IL2, KIT, CXCL12 and OX40. Notably, ILCs and Tregs exhibited strong bidirectional communication through the OX40, KIT and IL2 signaling axes, supporting a functional interaction between these populations (Figure 6B).

**Figure 6.**
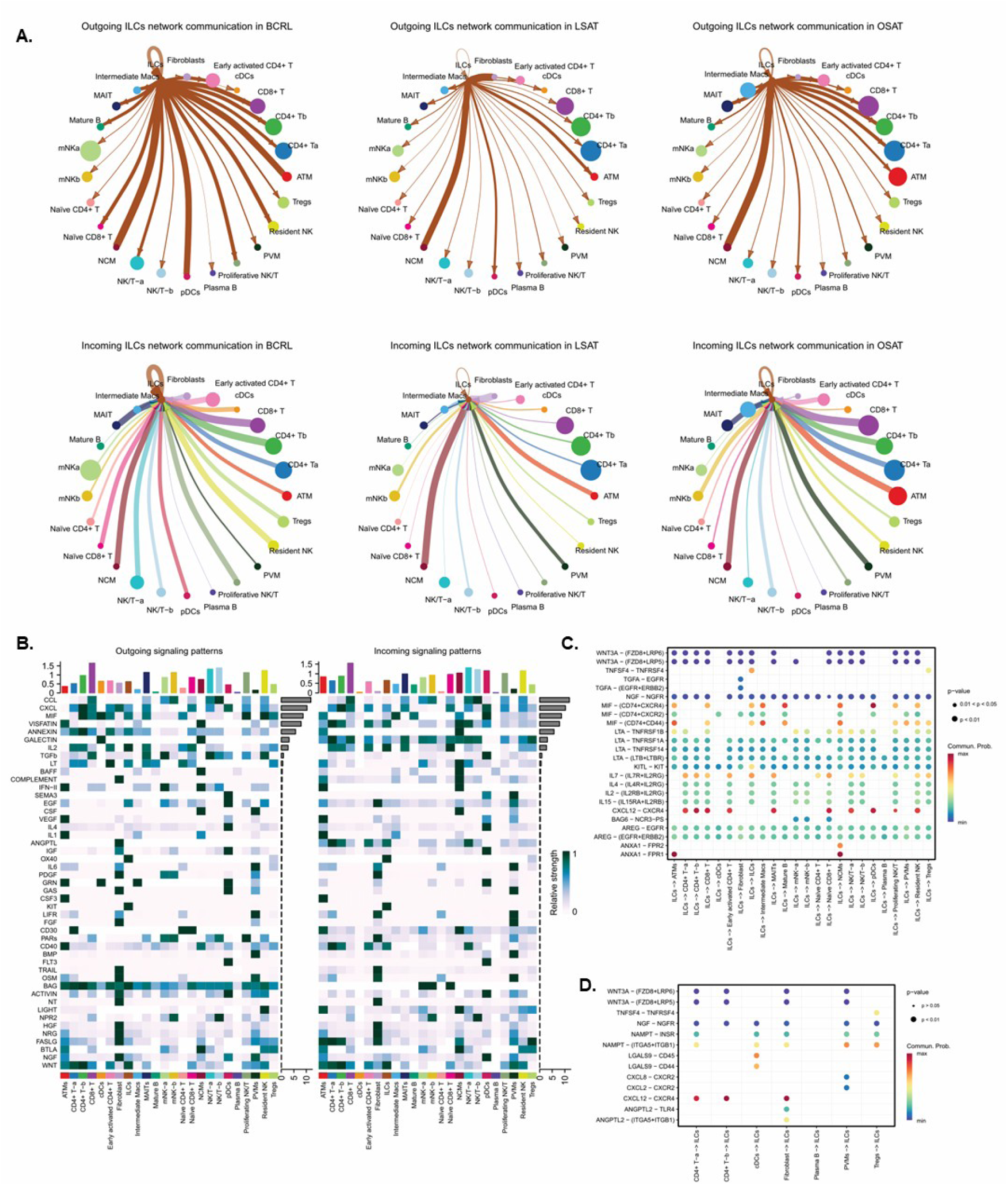
ILCs are key immune signaling communicators in BCRL. **A.** Incoming and outgoing communication strength of ILCs compared with other immune populations across conditions. Node size reflects the overall interaction strength of each cell group, and edge thickness represents the relative communication probability between interacting cell types. B. Heatmap showing global signaling patterns across BCRL cell populations. Color intensity represents relative signaling activity/communication probability. Top bars indicate the number of interactions contributed by each signaling pathway, and right-side bars represent overall signaling strength across cell groups. C. Ligand-receptor interaction bubble plot showing signaling pathways sent by major immune cell populations toward ILCs. D. Ligand-receptor interaction bubble plot showing signaling received by ILCs from other major immune cell groups. For C and D, dot color represents communication probability, and dot size indicates interaction significance/strength.

Next, we examined the individual ligand-receptor (L-R) pairs driving the most significant signaling sent (Figure 6C) or received (Figure 6D) by ILCs in BCRL. ILCs exhibited extensive crosstalk with monocytes and pDCs through ANNEXIN, LT, and MIF signaling pathways (Figure 6C). Importantly, we identified OX40 signaling as a specific, bidirectional communication axis between ILCs and Tregs in BCRL (Figure 6C). A similar pattern was observed for IL7 signaling, which appeared to preferentially connect ILCs with multiple T cell subsets, including Tregs, CD4^+^ T and CD8^+^ T cells. These findings raise the possibility that increased OX40, IL7 and LT signaling across ILCs and multiple immune populations may contribute to the maintenance of chronic immune activation and tissue remodeling in BCRL.

### ILC2 and ILC3 cells accumulate in BCRL-affected tissue

The presence of ILC subsets within the subcutaneous adipose tissue of patients affected by BCRL was further confirmed by flow cytometry (Figures 7A, B). For comparison, we used SVF samples from BCRL adipose tissue (BCRL 2–7) and from control breast subcutaneous adipose tissue (BSAT 1–7) adipose tissue. No significant differences were found in age (58,1 ± 6,4 vs 47 ± 13,1 years) and BMI (26,6 ± 1,7 vs 29,5 ± 3,8 kg/m2) between the groups, respectively.

**Figure 7.**
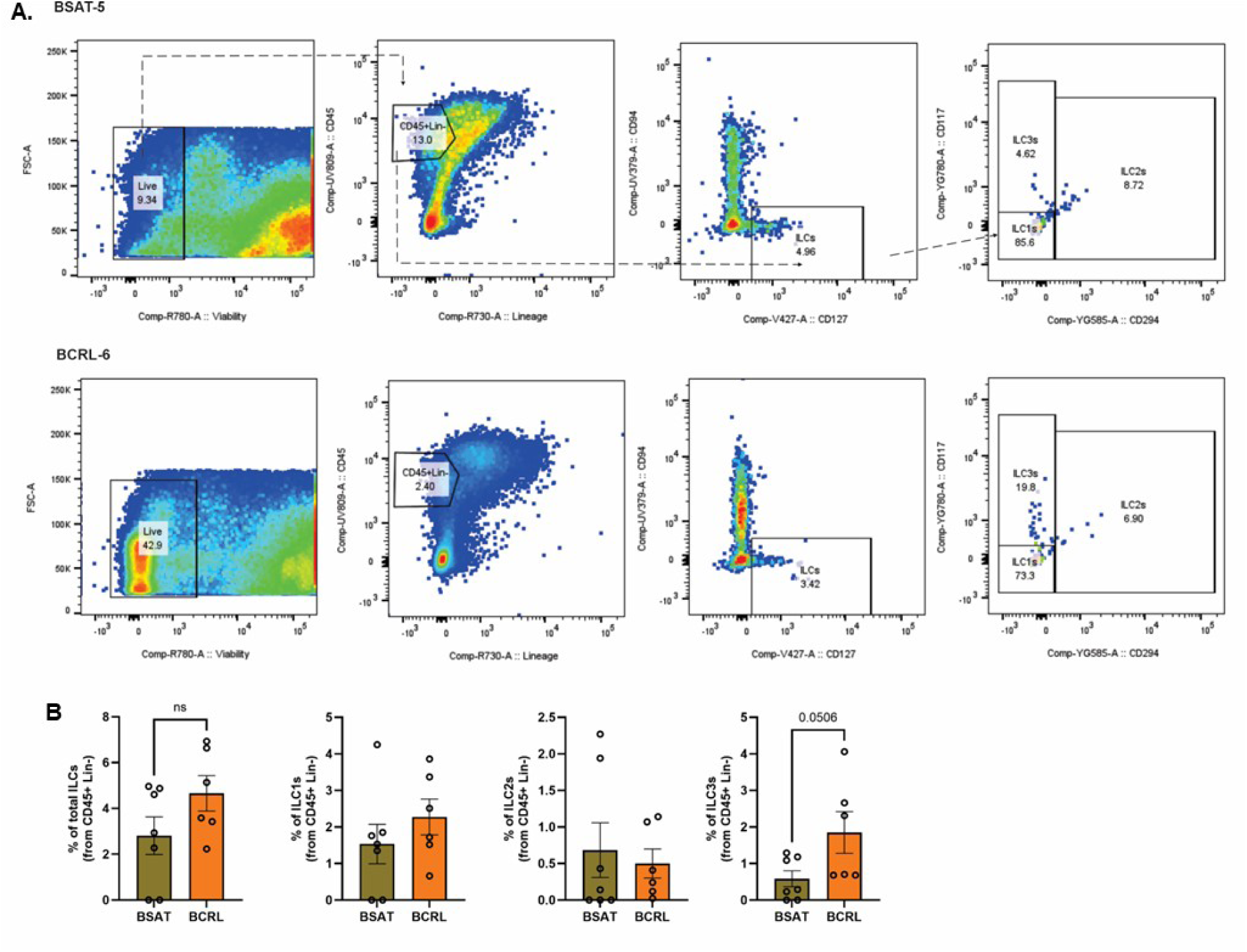
Flow cytometric validation reveals expansion of ILC2 and ILC3 subsets in BCRL adipose tissue. **A.** Representative flow cytometry gating strategy used to identify total innate lymphoid cells (ILCs) and their subsets (ILC1, ILC2 and ILC3) in BCRL and control breast subcutaneous adipose tissue (BSAT). **B.** Quantification of total ILCs and ILC subsets in BCRL (*n*= 6) and control BSAT (*n*= 6) samples. Data presented as the mean frequency of cells within total live CD45⁺ cells ± SEM. Statistical significance was determined using Mann-Whitney test; \**p*< 0,05.

In contrast to the scRNA-seq findings, the overall frequency of CD45⁺Lin⁻ (CD3, CD14, CD16, CD19, CD20, CD56) CD94⁻CD127⁺ ILCs measured by flow cytometry was comparable between BCRL and control adipose tissue (Figure 7B), representing approximately 1% of the total SVF (Figure 7A). Large inter-patient heterogeneity was observed in both ILC frequencies and the expression of canonical ILC markers. Phenotypic characterization of the individual ILC subsets in BCRL revealed a shift particularly in the ILC3 (CD117⁺⁺CRTH2⁻) frequency compared to the control BSAT (Figure 7B, *p =* 0,0506). Unfortunately, we could not detect differences in the frequency of ILC2 cells (CD117⁺CRTH2⁺) and ILC1 cells (CD117⁻CRTH2⁻) in BCRL compared to BSAT control tissue samples (Figure 7B).

### BCRL adipose tissue displays a pro-inflammatory and co-stimulatory protein signature

To further investigate the cytokine landscape associated with BCRL at protein level, we performed multiplex inflammatory protein profiling on BCRL adipose tissue lysates (BCRL 2, 3, 5–9, 12). BSAT lysates (BSAT 2–5, 14–17) served as controls. A customized panel comprising 18 proteins selected from our scRNA-seq results was analyzed using a LUMINEX platform. Compared with controls, BCRL samples showed enrichment of inflammatory and immune co-stimulatory mediators (Figure 8A). MIF (*p =* 0,020) and TNFα *(p =* 0,023) levels were significantly increased, while ICOSL (*p =* 0,081) displayed a trend toward higher expression in BCRL samples (Figure 8A). We also detected increased levels of IL1RA (*p =* 0,0140). Unfortunately, this method did not detect a significant change in OX40 levels between BCRL and control tissue lysates (Figure 8A; *p =* 0,533). However, these findings further support the presence of a complex and dynamically regulated immune microenvironment in BCRL. Notably, several cytokines included in the multiplex profiling panel were below the limit of detection in the tissue lysates and therefore could not be reliably quantified.

**Figure 8.**
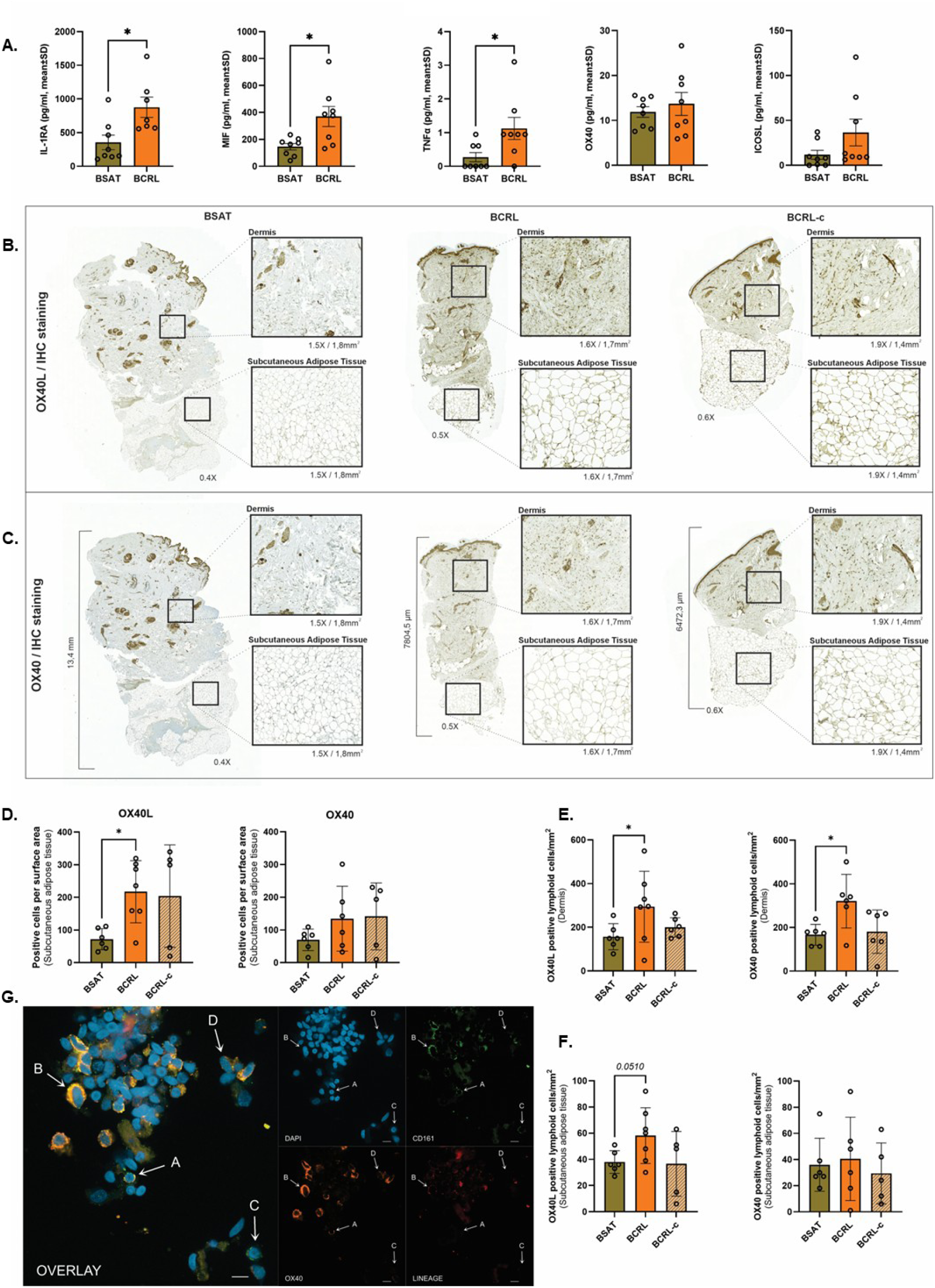
Chronic BCRL is characterized by a pro-inflammatory and co-stimulatory tissue immune microenvironment. **A.** Levels of MIF, IL1RA, TNFα, OX40 and ICOSL in BCRL subcutaneous adipose tissue (*n*= 8) compared with control BSAT tissue (*n*= 8). Data are shown as mean ± SEM. Statistical significance was determined using a Mann-Whitney test (*\*p*≤ 0,05). Representative images of **(B)** OX40L and **(C)** OX40 IHC stained skin and subcutaneous adipose tissue sections from healthy BSAT (*n*= 6) and BCRL (affected; *n*= 7) and non-affected contralateral tissue (BCRL-c; *n*= 5). Quantification of OX40L and OX40-positive lymphoid cells stained by IHC in **(D)** dermal and **(E)** subcutaneous fractions of BSAT, BCRL and BCRL-c tissue sections. Bars represent the number of stained positive lymphoid cells per mm² ± SD determined by IHC-based pathological assessment. Statistical analysis was performed using the Kruskal-Wallis test followed by Dunn’s multiple-comparison test *(p< 0,05).* **F.** Bar plots showing the amount of positive OX40L and OX40 cells in the overall subcutaneous adipose fraction of BSAT, BCRL and BCRL-c tissue sections. Analysis was performed using Visiopharm. Data are presented as number of positive cells per area ± SD. Statistical analysis was performed using Brown-Forsythe and Welch ANOVA tests (\**p*< 0,05). **G.** Immunofluorescently stained OX40-expressing ILC (cell A) among SVF-isolated cells isolated from BCRL subcutaneous adipose tissue. ILCs were defined as Lin⁻ (CD3, CD14, CD16, CD19, CD20, CD56; red), CD161⁺ (green), and OX40⁺ (orange). DAPI (blue) was used for nuclear staining. Scale bars: 10 µm. For comparison, cell B is CD161^+^OX40^+^Lin^+^, cell C is CD161^+^OX40^−^Lin^−^, and cell D is CD161^−^OX40^+^Lin^−^. Imaged using Nikon Ti2 (60x).

### OX40/OX40L pair expression is higher in BCRL dermis and subcutaneous adipose tissue

Given the prominent OX40 co-stimulatory signature observed in BCRL (Figures 8B, D-F), we next examined the distribution and relative abundance of OX40 co-stimulatory pair proteins in histological tissue sections from BCRL and BSAT. Representative images in Figures 8B and 8C show OX40L and OX40 immunohistochemical staining in BCRL tissues compared with control BSAT samples.

An automated analysis of the skin and adipose tissue compartments of the samples was performed by Visiopharm platform using a highly sensitive nuclear-detection algorithm. When normalized to total cellular content within the dermis, no significant differences in overall OX40L or OX40 positivity were detected between BCRL and control tissues (data not shown). However, significantly increased OX40L-positive cell frequencies were detected within subcutaneous adipose tissue from BCRL tissues compared with healthy BSAT controls (Figure 8D, left-panel; *p=* 0,010). Although several BCRL adipose tissue samples also displayed increased OX40-positive cell frequencies, these differences did not reach statistical significance overall (Figure 8D, right-panel).

Due to the lack of specific markers for identifying immune cells, OX40L and OX40-positive lymphoid cells were independently quantified by an experienced pathologist using the same samples. The dermis of BCRL biopsies contained significantly higher numbers of OX40L-positive (Figure 8E, left-panel*; p=* 0,033) and OX40-positive lymphoid cells (Figure 8E, right-panel; *p=* 0,029) compared with healthy BSAT dermis. To note, the amount OX40L and OX40 positive lymphoid cells appeared to be similar in the unaffected contralateral arm from BCRL patients than that of healthy BSAT tissues (Figures 8E and F). Within the subcutaneous adipose tissue compartment, identification and quantification of positive lymphoid cells was technically limited by the relatively small amount of adipose tissue present in a large subset of biopsies. Nevertheless, increased numbers of OX40L-positive lymphoid cells were observed in BCRL adipose tissues compared with healthy BSAT controls, reaching borderline statistical significance (Figure 8F, left-panel; *p=* 0,051). In contrast, OX40-positive lymphoid cell numbers within subcutaneous adipose tissue appeared comparable between groups (Figure 8F, right-panel).

To confirm the expression of the OX40 receptor in ILCs and thus their possible contribution to OX40 signaling in BCRL adipose tissue, SVF cells from a representative BCRL tissue were allowed to adhere to CD117 antibody coated slides and stained with antibodies against ILCs and OX40. Representative images revealed an OX40 receptor-associated signal in cells with a phenotype similar to that of ILCs (Lin^−^/CD161^+^) and also in Lin^+^ cells (Figure 8G).

## DISCUSSION

This study provides a comprehensive characterization of the immune microenvironment in chronic BCRL tissue, uncovering new insights into disease pathogenesis. By integrating single cell transcriptomics, cell-cell communication analysis, and tissue-level validation, our study highlights the expansion and activity of ILC2 and ILC3 immune cell populations, as well as amplified OX40 signaling, as novel defining features of the disease. Rather than single players, ILCs emerge as a central communication hub that actively bridges innate and adaptive immune responses. This cellular crosstalk links ILCs, T, B, and NK cells, and Tregs with an inflammatory, tissue-remodeling, and co-stimulatory signaling network, most notably through IL2, OX40, KIT, and LT signaling. Within this network, a robust, bidirectional axis between ILCs and Tregs through OX40 is uniquely enriched in BCRL tissues. These transcriptomic interactions are mirrored at the protein level, confirming increased levels of local inflammatory mediators and distinct enrichment of the OX40/OX40L co-stimulatory pair in BCRL-affected tissues. These findings reveal novel molecular features associated with BCRL and suggest that coordinated interactions between innate and adaptive immune populations promote chronic inflammatory and tissue-remodeling responses within affected tissues.

Experimental models have established that lymphatic dysfunction promotes chronic immune activation characterized by accumulation of CD4^+^ T cells together with Th2-associated cytokines, thus promoting fibrosis and adipose tissue expansion (11,12,51–53). However, studies in human lymphedema have reported more heterogeneous changes in immune-cell composition, likely reflecting differences in disease stage, tissue compartment and anatomical location. In our stage II human BCRL cohort, we did not observe major changes in the overall frequencies of CD4^+^ T cells, NK/T cells or DCs compared with healthy adipose tissue, whereas myeloid populations, particularly macrophages, were reduced. In contrast, other human studies using skin biopsies or dermolipectomy specimens, as well as murine models, have reported unchanged or increased macrophage abundance (54,55). Together, these findings emphasize that the immune landscape of lymphedema remains incompletely understood and highlight the need for further studies comparing different disease stages, tissue compartments, and experimental models.

Among the immune populations altered in BCRL, Tregs were consistently expanded and exhibited an activated transcriptional profile. This finding agrees with previous reports in experimental lymphedema (12,56) and may reflect the metabolic remodeling that accompanies adipose tissue expansion, as Treg maintenance is strongly influenced by lipid metabolism (57–59). Interestingly, studies of stage III or advanced lymphedema reported no changes in Treg abundance (24,54), while experimental models suggest progressive impairment of Treg suppressive function (12,60). Together, these observations suggest that Treg accumulation may represent an early adaptive response that changes as disease progresses.

Our study identified ILC2 and ILC3 populations as novel cellular and signaling contributors in BCRL. Although the identification of tissue-resident ILC subsets in humans remains challenging due to their low abundance and phenotypic plasticity, our transcriptomic and flow cytometric analyses consistently demonstrated increased ILC populations in BCRL tissues. ILCs in BCRL expressed a unique transcriptional signature associated with tissue remodeling, immune cell recruitment, and inflammatory activation, including AREG, LTB, XCL1, and TNF receptor superfamily members. Similarly, these responses have been described in several transcriptomics studies on secondary lymphedema (61). In other chronic inflammatory and Th2 cell-mediated diseases, such as asthma, atopic dermatitis, and obesity (19–21,62–65), evidence highlights the contribution of ILCs to aberrant stromal activation, fibrosis, and persistent inflammation driving these processes (20,66). Our data suggest that similar molecular mechanisms may contribute to lymphedema progression.

In addition to altered immune cell composition, our analyses support the presence of a highly active and remodeled immune microenvironment in BCRL tissues. Transcriptomic profiling, RNA velocity analysis, and cell-cell communication analyses collectively demonstrated enhanced inflammatory and co-stimulatory signaling, together with altered immune cell state dynamics across several innate and adaptive immune populations, particularly ILCs, Tregs, B cells, DCs, and monocytes. Cell communication analysis identified ILCs as central signaling hubs within BCRL tissues, exhibiting strong predicted interactions with Tregs, pDCs, cytotoxic lymphocytes and stromal populations through OX40, IL2, KIT, LT, CXCL12, MIF signaling pathways. Among these, CXCL12 has recently been identified as a candidate gene associated with CD4⁺ T-cell immune network remodeling in secondary lymphedema (67). Moreover, RNA velocity analysis revealed that immunological differences in BCRL extend beyond changes in cell abundance and involves altered immune cell state dynamics. Consistent with their increased abundance, ILCs displayed marked shifts in latent-time distributions and reduced trajectory coherence, supporting persistent activation and transcriptional remodeling. Similar alterations were also observed in CD4^+^ T cells, dendritic cells, B cells and monocytes despite the absence of major frequency changes in some of these populations, suggesting that the BCRL microenvironment broadly affects immune-cell activation and functional-state transitions.

Protein analysis of BCRL tissue lysates further supported the presence of an activated and immune transitional microenvironment, characterized by elevated MIF, TNFα (68,69), and IL1RA (70) levels. Importantly, histological analyses demonstrated increased expression of the OX40L/OX40 co-stimulatory pair in both dermal and subcutaneous adipose tissue compartments of BCRL tissues. As OX40 signaling regulates both T-cell activation and persistence in other chronic and Th2-driven diseases (71,72), it may drive a similar local response in lymphedema. To our knowledge, this study is the first to identify increased OX40 expression and predicted OX40 signaling activity in BCRL tissues, particularly between ILCs and Tregs. These findings suggest that OX40 signaling may contribute to the maintenance of the chronic inflammatory and tissue-remodeling microenvironment characteristic of BCRL. Given the ongoing clinical development of OX40-targeting therapies for other inflammatory diseases (73–75), our findings highlight this pathway as a potential therapeutic target for limiting inflammation and pathological tissue remodeling.

In conclusion, our study describes for the first time enriched ILCs activity and enhanced OX40 signaling as central features of the immune microenvironment in BCRL. The identification of ILCs in lymphedematous adipose tissue fills an important gap in the current understanding of lymphedema immunopathology. While previous work has established critical roles for Th2 cells, Tregs, and DCs, the upstream cellular mechanisms coordinating these sustained responses have remained elusive. As innate lymphoid cells are key regulators of inflammation, fibrosis, and tissue remodeling, their expansion in lymphedematous tissue provides a plausible mechanistic link between innate and adaptive immune responses. Thus, ILCs may represent a previously unrecognized cellular component that helps orchestrate the complex immune microenvironment characteristic of lymphedema through the OX40 co-stimulatory pathway. Our findings provide new insight into key processes underlying lymphedema pathogenesis, suggesting that while lymphatic dysfunction initiates the disease, its progression is driven by coordinated and sustained innate and adaptive immune signaling.

## MATERIAL AND METHODS

### Patient samples

**Table 1.** Demographic characteristics, breast cancer surgical approach, adjuvant oncological treatments, lymphedema surgery, and liposuction volume in 12 patients with BCRL and one patient* with cervical cancer-related lower extremity lymphedema.

| ID | Age | BMI (kg/m<sup>2</sup>) | Breast cancer surgery | Axillary LN surgery | Radiotherapy | Chemotherapy | BCRL duration (yrs) | ISL-Classification | Treatment year | Procedure | Liposuction volume (mL) | Used for |
| --- | --- | --- | --- | --- | --- | --- | --- | --- | --- | --- | --- | --- |
| BCRL-1 | 57 | 27.3 | MX | ALND | Yes | Yes | 16 | II | 2022 | Liposuction | 950 | 1 |
| BCRL-2 | 55 | 30.9 | BCS | ALND | Yes | Yes | 11 | II | 2023 | Liposuction | 600 | 1,2,3 |
| BCRL-3 | 61 | 28.9 | BCS | ALND | Yes | No | 8 | II | 2023 | Liposuction + VLNT | 800 | 1,2,3 |
| BCRL-4 | 57 | 29.7 | MX | ALND | Yes | No | 2 | II | 2023 | Liposuction | 800 | 1,2,4 |
| BCRL-5 | 61 | 31.2 | BCS | ALND | Yes | Yes | 10 | II | 2024 | Liposuction | 400 | 2,3 |
| BCRL-6 | 67 | 22.7 | MX | ALND | Yes | No | 1 | II | 2024 | Liposuction + LVA | 600 | 2, 3, 4 |
| BCRL-8 | 54 | 27.6 | MX | ALND | Yes | Yes | 3 | II | 2023 | Liposuction + DIEP + VLNT | 400 | 3 |
| BCRL-9 | 63 | 29.9 | MX | ALND | Yes | Yes | 4 | II | 2023 | Liposuction + VLNT | 1300 | 3 |
| BCRL-10 | 59 | 29.2 | MX | ALND | Yes | Yes | 2 | II | 2024 | Biopsy only | - | 4 |
| BCRL-11 | 54 | 29.3 | MX | ALND | Yes | Yes | 3 | II | 2024 | Liposuction + DIEP + VLNT | 660 | 4 |
| BCRL-12 | 43 | 30.0 | MX | ALND | N/A | N/A | 7 | II | 2024 | Liposuction | 1100 | 3,4 |
| BCRL-13 | 54 | 21.6 | MX | ALND | Yes | Yes | 1 | II | 2025 | Liposuction + DIEP + VLNT | 600 | 4 |
| BCRL-14 | 64 | 29.4 | BCS | ALND | Yes | Yes | 3 | II | 2026 | Liposuction | 900 | 5 |
| CRL-7* | 48 | 34.4 | ** |  | Yes | Yes | 7 | II | 2024 | Liposuction | 3000 | 2,3,4 |

BCRL= Breast Cancer Related Lymphedema, \*CRL= Cancer Related Lymphedema. ISL-Classification= International Society of Lymphology (76), BMI= Body mass index, MX= Mastectomy, BCS= Breast-conserving surgery, ALDN= Axillary Lymph node dissection, SLNB= Sentinel lymph node biopsy, DIEP+VLNT= Vascularized lymph node transfer with breast reconstruction, LVA= Lymphaticovenous Anastomosis. \*\*SLNB of the lower abdomen: pelvic lymph nodes were removed on the left side from the external iliac region as well as the obturator lymph nodes. Used for: (1) scRNA-seq, (2) Flow cytometry, (3) Cytokine multiplex, (4) OX40 tissue staining, (5) SVF cell staining.

Patients undergoing liposuction surgery for stage II BCRL at Turku University Hospital were recruited in the study. Clinical information was collected from the hospital’s electronic medical records and is shown in Table 1. BCRL-10 patient underwent adipose tissue sampling from the affected upper extremity without therapeutic liposuction. As controls, subcutaneous adipose tissue was obtained from healthy donors undergoing breast reduction mammoplasty **(BSAT)** using immediate liposuction of the removed tissue pieces (Supplementary Table 1). All recruited patients signed a written informed consent to allow the use of their tissue samples in the study. The collection of clinical samples has been approved by the ethics committee of the wellbeing services county of Southwest Finland (11/1801/2022, 23/1801/2018).

### Stromal vascular fraction isolation

Cells from the stromal and vascular fractions were isolated following a modified protocol modified from Felix I, et al. (77). Freshly collected lipoaspirate from BCRL patients or lipoaspirate from the subcutaneous layer of the reduction mammoplasty tissue piece was digested using 1 mg/mL collagenase D (Roche, Mannheim, Germany), 50 µg/mL DNase I (Roche) and 37,5 µg/mL Liberase (Roche) in DMEM-F12 medium for 1 hour under 37 °C shaking incubation. Post-incubation, cell suspension was centrifuged at 1000x g for 1,5 min at RT. After aspiration of the supernatants, red blood cells were lysed using BD Pharm Lyse*™* lysing buffer (BD Biosciences, Franklin Lakes, NJ, USA) for 3 min. The remaining SVF cell pellet was resuspended in PBS containing 5mM EDTA and filtered through a 70 µm cell strainer. Finally, the pellet was washed several times with PBS-EDTA and frozen at 196°C for later use.

### FACS sorting

BCRL immune cells were sorted from the isolated SVFs with Sony SH800 Cell Sorter (Sony Biotechnologies, CA, San Jose, USA). SVF containing cells were labelled with Fixable Viability Dye eFluor 780 (eBioscience, Thermo Fisher Scientific, Waltham, MA, USA; 65086514) and PE-CD45 (BioLegend, San Diego, CA, USA; HI30, 304058). Single cells were gated with the FSC-H versus FCS-W plot, followed by live and CD45^+^ plots. Live single CD45^+^ cells were sorted into RPMI medium containing 2 % FBS.

### Single cell RNA sequencing

Freshly sorted CD45^+^ cells were processed immediately according to 10X Genomics guidelines (10x Genomics, Pleasanton, CA, USA). With a targeted cell range of 5.000–10.000, the libraries were prepared using the Chromium Next GEM Single Cell 3’ Reagent Kit v3.1 in the Single Cell Omics core (Finland). Bioanalyzer assessment ensured high-quality samples, with an average library fragment size of 465 bp. Subsequent sequencing was performed at the Finnish Functional Genomics Center at Turku Bioscience (Finland) using the NovaSeq 6000 S2 v1.5 instrument (Illumina, San Diego, CA, USA) generating 28 + 90 bp reads.

### Public datasets

Additional lean and obese control scRNA-seq datasets were obtained from NCBI Gene Expression Omnibus (GEO) database under the accession number GSE155960(21). From this dataset, we downloaded samples from CD45^+^ lean donor 1 (**LSAT_1**, GSM4717158), donor 2 (**LSAT_2**, CGSM4717159) and donor 3 (**LSAT_3**, GSM4717160), and CD45^+^ obese donor 1 (**OSAT_1**, GSM4717161), donor 2 (**OSAT_2**, GSM4717162) and donor 3 (**OSAT_3**, GSM4717163).

### Data integration, clustering and cell type annotation

All scRNA-seq data were processed using RStudio (v.2023.09.0, Boston, MA, USA). Every filtered feature matrix from all downloaded controls and BCRL samples (**BCRL_1–4**) were transformed into individual Seurat objects using R package Seurat (v.4.4.0, New York, NY, USA). Cells with lower than 300 or higher than 5,000 expressed genes were removed. We further discarded cells with mitochondrial gene content higher than 10%. Finally, we obtained the following number of cells for the downstream analysis: 11.775, 7.780, 5.322 cells for LSAT-1, 2 and 3; 5.997, 5.871, 7.773 cells for OSAT-1, 2 and 3; and 4.673, 3.293, 4.382, 8.173 cells for BCRL-1, 2, 3 and 4. After removal of doublets and dead cells, all Seurat objects were merged and data underwent SC-Transform normalization. For correcting batch effects, we performed integration of the 10 samples. We merged and split the resulting data into individual donor samples, from which we obtained the integration features and integration anchors using the functions *SelectIntegrationFeatures* and *FindIntegrationAnchors,* respectively. A total of 65.039 cells were used for the *IntergateData* function. The integrated data was utilized as default data for downstream analysis.

Next, we performed data scaling via the *ScaleData* function and Principal component analysis (PCA). After applying UMAP dimensional reduction to the first 50 PCA components, cells were then clustered at a 0.6 resolution through *FindNeighbors* and *FindClusters* functions. Using these settings, we found 26 clusters. The markers of these clusters were obtained using the *FindAllMarkers*. Two clusters were removed due to signs indicative of impaired mitochondrial functioning (poor expression of genes related to oxidative phosphorylation, cholesterol biosynthesis and glycolysis). The cells in the residual 24 clusters were manually annotated based on the expressed markers.

To further explore the relationships between identified clusters, we performed hierarchical clustering. Specifically, we computed average expression levels for each gene across the identified clusters using the *AggregateExpression* function. We then applied hierarchical clustering (to group similar cell types based on their transcriptomic signatures and visualized the resulting dendrogram showing the hierarchical relationships between the cell types. To analyze and compare the contribution of each sample based on cell number and condition, the number of cells for each cell type from each sample was determined. This number was then divided by the total number of cells for each sample and transformed to percentage. For comparisons between conditions (LSAT, OSAT and BCRL), we aggregated the number of cells for each cell type from samples across all three conditions. We then divided this aggregated number by the total number of cells from all samples for each condition and expressed it as a percentage.

### Differential gene expression and gene enrichment analysis

To identify key transcriptional differences across conditions, we performed differential gene (DE) expression analysis using the total integrated single-cell dataset or a specific cluster subsetted data, comparing each condition (BCRL, OSAT and LSAT) against the other two. DE genes were obtained using *FindAllMarkers* function. Genes with an adjusted p-value (Benjamini–Hochberg correction) < *0,05, log₂ fold change* > 1, and expression in more than 10% of cells were considered significantly upregulated. The top 15 genes ranked by log₂ fold change were selected for visualization. DE genes from total single-cell data were then subjected to Gene Ontology (GO) and Reactome pathway enrichment analyses using the clusterProfiler package (v4.10.1). Gene symbols from the integrated dataset were converted to Entrez IDs using the *bitr* function. For GO analysis, we applied the *enrichGO* function with the org.Hs.eg.db annotation database, “BP” (biological process) as the ontology category, and the Benjamini–Hochberg (BH) method for multiple testing correction. For Reactome pathway analysis, enrichment was performed using the *enrichPathway* function from the ReactomePA package, with BH-adjusted p-values used to determine significance. Results from both analyses were further filtered to only include significant (*p> 0,05*) processes and pathways related to immunity, using the description *terms= “immune |cytokine |interleukin“*.

### RNA velocity analysis

Spliced and unspliced mRNA abundances were quantified from GRCh38-aligned BAM files using the velocity toolset (78). RNA velocity and cellular trajectories were computed using the dynamical modeling framework in scVelo (50) alongside Scanpy. After standard filtering, normalization, and moment computation, gene-specific velocities, latent time, and velocity confidence metrics were estimated using default parameters. Velocity-derived transcriptional states were projected onto the initial UMAP embedding and visualized by experimental condition (LSAT, BCRL and OSAT).

### Cell-Cell communication network analysis

To identify intercellular communication networks between the multiple immune cell populations identified in the previous analysis, we used CellChat (v.1.6.1, Irvine, CA, USA) (79). CellChatDB.human was used for the datasets. Briefly, BCRL and control LSAT and OSAT samples were first subsetted from the integrated Seurat object as described above and the corresponding CellChat objects were generated with the function *createCellChat*. Communication probability and inferring cell-cell communication network were predicted by *CommunProb* and *computeCommunProbPathway* (79). Statistically significant interactions were identified using permutation testing, and only communications with *p< 0,05* were retained for downstream analyses. We determined the major signaling sources and targets contributing to the outgoing or incoming signaling of all cell types using the function *netAnalysis_signalingRole_scatter*. Chord diagrams were used for visualization of interactions and incoming and outgoing pathways strengths between ILCs from all the 3 conditions and other cell types. A relative information flow analysis was performed using the merged BCRL, LSAT and OSAT CellChat objects to compare the contribution of each signaling pathway between these conditions. Top 10 ligands and receptors with the highest activity in these signaling pathways for each condition were ranked and visualized using the *netAnalysis_contribution* function.

### Spectral flow cytometry

SVF collected cells from subcutaneous adipose tissue derived from BCRL-affected arm (*n*= 6) and control BSAT (*n*= 7) were counted and total of 1 ×10^6^ were used for ILCs identification and quantification using BD FACSymphony™ A5 SE Cell Analyzer. Cells were stained with Fixable Viability Dye eFluor 780 to label dead cells. Unspecific binding to low-affinity Fc-receptors was blocked by incubating the cells with Human Fc Receptor Binding Inhibitor (BD Bioscience; 564220). Surface antigens were stained for 30 min on ice with conjugated fluorescently labelled antibodies against human CD45 (BD Biosciences, BUV805 HI30, 612892), Lin (Bio-Rad, Carlsbad, CA, USA; AlexaFluor 700, MCA2692A700T), CD11b (BD Biosciences; BV650 M1/70, 563402), CD94 (BD Biosciences; BUV395 HP-3D9, 743954), CD127 (BD Biosciences; BV421 HIL-7R-M21, 562437), CD117 (BD Bioscience; PE-Cy7, 104D2, 339217) and CD294 (Biolegend; PE, BM16, 350105). Samples were acquired using the FACSymphony A5 flow cytometer (BD Bioscience) with the following laser configuration: 355nm, 405nm, 488nm, 561nm and 637nm, and with FACSDiva™ Software (BD Bioscience, v.9.6). Data were analyzed using FlowJo™ (v.10.10.0, BD Bioscience).

### Cytokine and chemokine quantification

Quantification was performed on cryopreserved subcutaneous adipose tissue lipoaspirates obtained from BCRL (*n*= 8) and control BSAT (*n*= 8) tissues. Samples were lysed in ice-cold RIPA buffer (Thermo Scientific; 89900; 500 µL per 100 mg tissue) supplemented with protease and phosphatase inhibitors (Thermo Fisher Scientific; A32961) and homogenized using a TissueLyser (Qiagen, Helsinki, Finland). Cytokine and chemokine levels were measured using a custom Human ProcartaPlex Mix&Match 18-plex panel (Thermo Fisher Scientific; PPX-18-MXH6DCJ) targeting Th1- and Th2-associated inflammatory factors. Multiplex assay plates were prepared according to the manufacturer’s instructions using samples diluted 1:2 in assay buffer. Analyte concentrations were acquired on a Luminex 200 system (Luminex Corporation, Austin, TX, USA) using xPONENT 3.1 software (xMAP technology).

### Immunohistochemical tissue staining and quantification

Biopsies with subcutaneous adipose tissue from BCRL-affected (*n*= 7) and non-affected patient arms (*n*= 5) and healthy control biopsies with skin and subcutaneous adipose tissue from the breast area of breast reduction patients (BSAT, *n*= 6) were collected and placed in formalin in the operating room and embedded in paraffin after 24 h fixation. Tissue sections were cut at 4 µm thickness. Immunohistochemical staining was performed by the Histology core facility of the Institute of Biomedicine, University of Turku, Finland. Unspecific binding was blocked with Draco antibody diluent (WellMed AD125, WellMed BV, Holland, Netherlands). The primary antibodies used were OX40 (NeoBiotechnologies, Union City, CA, USA; OX40/3108, 7293-MSM2) and OX40L (BioLegend; 11C3.1, 326302) at dilutions 1:200 and 1:100, respectively, for 60 min at RT. Endogenous enzyme blocking was performed using 1% H2O2. Secondary antibodies used were BrightVision one step detection system goat anti-mouse HRP (WellMed T100HRP) for 30 min before detection with BrightDAB (WellMed BS04-110) for 10 min at RT and counterstaining with Mayer’s hematoxylin for one minute. After rinsing, sections were dehydrated and mounted using Pertex. The sections were imaged using a Pannoramic 250 scanner with a Plan-Apochromat 20x objective and 0.22 µm pixel size.

Quantitative analysis of OX40L and OX40 IHC staining was performed by two different methods. First, the overall number of OX40L- and OX40-positive cells was quantified with Visiopharm 2025.08.4 x64 (Visiopharm Integrator System, Hørsholm, Denmark), a digital image analysis software incorporating artificial intelligence–based algorithms. A pre-trained IHC Nuclei Detection AI app (10170) was applied with a three-pixel dilation to include the nuclear and immediately surrounding perinuclear area. A semi-quantitative grading system was then applied to assess staining intensity and classify cells into negative and OX40L- or OX40-positive groups. Regions-of-interest (ROIs) were manually drawn in the dermal and adipose tissue compartments (Supplementary figure 4) and areas containing blood vessels, hair follicles or glands, tissue artefacts, broken cells and large empty spaces were excluded from the analysis. Positively stained cells were divided in intensity grades (0–3), of which moderate-to-high intensity objects (grades 2–3) were included for downstream analysis.

To specifically focus on the OX40L and OX40-positive lymphoid cells in the tissues, a pathologist blinded to patient identity and sample conditions quantified the respective cell population. Lymphoid cell infiltrates were identified within the dermal and subcutaneous adipose tissue compartments using HE stained slides. Quantification was subsequently performed by counting OX40L- and OX40-positive lymphoid cells, followed by normalization to the analyzed tissue area. Results are expressed as the number of OX40L- or OX40-positive cells per mm². Adjacent HE stained tissue sections were used for detection and quantification.

### Immunofluorescence analysis

Specialty microscope slides with 5 mm wells (Epredia Netherlands B.V., Breda, Netherlands; X1XER302W) were coated overnight with anti-human CD117 (c-Kit) antibody (R&D Systems, Minneapolis, MN, USA; AF1356). Stromal vascular fraction (SVF) cells from BCRL subcutaneous adipose tissue were incubated on the slide for adhesion and fixed with 4% paraformaldehyde before blocking with TBST containing 1% BSA and 5% NGS (Jackson ImmunoResearch Laboratories, Cambridge, UK; 005-000-121). The cells were then incubated overnight with anti-human CD161 (Abcam, Cambridge, UK; ab259916) and OX40 antibodies diluted 1:50. Second-day staining was performed with secondary and conjugated antibodies including goat anti-rabbit Alexa Fluor 488 (Invitrogen, Thermo Fisher Scientific; A11034, 1:300), donkey anti-mouse Alexa Fluor 568 (Invitrogen; A10037, 1:300), and an Alexa Fluor 700-conjugated anti-lineage antibody cocktail (Bio-Rad; MCA2692A700T, 1:100). Nuclei were stained with DAPI (1:1000) for 5 min, and the slides were mounted using ProLong Gold Antifade Mountant (Thermo Fisher Scientific; P36930).

Imaging was performed using a Nikon Ti2 spinning disk confocal microscope equipped with a Kinetix camera and a Spectra III/Celesta/Ziva multi-laser illumination system. Images were acquired using a PLAN APO 60× SIL λS OFN25 DIC N2 objective with a numerical aperture of 1.3 and oil immersion (refractive index 1.406). Acquired images had a pixel size of 0.11 μm and a bit depth of 12. Spectral unmixing was performed using NIS-Elements AR software (version 5.11.03).

### Statistical methods

Statistical analyses were performed using GraphPad Prism (version 10.4.1, GraphPad Software, La Jolla, CA, USA). Details of computational and statistical methods applied to single-cell RNA sequencing analyses are described in the corresponding section above. For quantitative comparisons performed for spectral flow cytometry, immunohistochemistry and multiplex protein measurements, data distribution was first assessed for normality using the Shapiro-Wilk test. Depending on data distribution, either parametric or non-parametric statistical tests were applied. Comparisons between two groups were performed using unpaired two-tailed t tests or Mann-Whitney tests, as appropriate. For analyses involving multiple experimental groups, one-way ANOVA followed by Tukey’s multiple-comparison test or Kruskal-Wallis followed by Dunn’s multiple-comparison test was used. All data are presented as mean ± SD unless otherwise indicated. A *p value ≤ 0,05* was considered statistically significant.

## LIMITATIONS OF THE STUDY

Several limitations of our study must be acknowledged. First, we used external control scRNA-seq datasets owing to the limited availability of healthy adipose tissue samples in quantities sufficient for comprehensive scRNA-seq analysis. Nevertheless, dataset integration was successful, and all major immune-cell populations were consistently detected across samples. In addition, SVF samples from reduction mammoplasty specimens were included as healthy controls in flow cytometry validation experiments. This approach was adopted because reduction mammoplasty tissue represented the only readily available source of healthy subcutaneous adipose tissue. To ensure that control samples underwent collection procedures comparable to those used for BCRL specimens, subcutaneous adipose tissue was harvested by liposuction directly from the surgical specimen in the operating room. The procedure was performed under the guidance of a plastic surgeon to ensure sampling from the appropriate subcutaneous adipose tissue layer. It is also important to note that expression of cytokine transcripts was generally low across ILCs and other immune populations in the scRNA-seq dataset. This may reflect the inherently low abundance and transient expression of cytokine genes at the single-cell level, as well as technical limitations associated with scRNA-seq analyses and dataset integration.

Finally, this study has several limitations related to the investigation of human ILCs. ILCs represent a rare immune-cell population within adipose tissue, and their frequency showed substantial inter-patient variability in our cohort. Together with the limited availability of patient tissue and the absence of robust, standardized methods for isolating sufficient numbers of viable human ILCs from adipose tissue, these factors currently restrict comprehensive functional *in vitro* analyses. Consequently, although our findings identify ILCs as key contributors to the BCRL immune microenvironment, future studies combining optimized human ILC isolation strategies with functional assays will be essential to validate their role and further dissect the molecular mechanisms underlying their interactions with other immune-cell populations.

## AUTHOR CONTRIBUTIONS

GMC, HJ, SP, EP, MS, PR, and PH designed the research. GMC, MR, HJ, and HG conducted the experiments. MR, HG, EP, MS, PR, and PH provided the reagents. GMC, MR, SP, KO, DK, and PH acquired the data. GMC, MR, SP, KO, DK, and PH analyzed the data. GMC, MR, HJ, SP, HG, KO, DK, EP, MS, PR, and PH wrote the manuscript.

## FUNDING SUPPORT

- Sigrid Jusélius Foundation (to PH)
- Research Council of Finland (to PH)
- ImmunDocs Funded Project (to PH and MR)
- Inflames Research Support (to GM)
- Lounais-Suomen syöpäyhdistys (to GM)
- Valtion tutkimusrahoituksen Varsinais-Suomen (VTR to GM, PH and SP)
- Lääketieten säätiö (Finnish Medical Foundation, to PH and SP)
- Vappu Uuspään säätiö (to PH and SP)

## ACKNOWLEDGMENTS

Cytometry was performed in the Cytometry Core, and Imaging was performed at the Advanced Imaging Core Facility at Turku Bioscience Center, supported by Biocentre Finland, the Finnish Advanced Microscopy Node of Euro-BioImaging Finland (Turku, Finland), and Turku Bioimaging. This work was supported by the Research Council of Finland, FIRI 2023 grant decision numbers 359073 and 358879, and FIRI 2024 grant decision numbers 367582 and 367577. The histological methods were performed by the Histology core facility of the Institute of Biomedicine, University of Turku, Finland.

---

**Supplementary table 1.** Demographic characteristics of LSAT, OSAT and BSAT controls

**Supplementary table 2.** Cluster-specific markers from all cells (Integrated dataset with 65.039 total cells)

**Supplementary table 3.** Mean latent time values and differential progression analysis by cell group, comparing BCRL against LSAT and OSAT control conditions.

**Supplementary figure 1.**

**A. Heatmap showing the contribution of each sample to the major/linage cell groups identified in the integrated scRNA-seq dataset.**

**B. Integrated UMAP embedding split by experimental condition, BCRL, LSAT and OSAT.**

**Supplementary figure 2.**

**A. Expression of immune gene signatures in Tregs in BCRL versus control tissues**

**B. Expression of immune gene signatures in DCs in BCRL versus control tissues**

**Supplementary figure 3.**

**A. UMAP plots showing the latent time inferred from RNA velocity analysis (scVelo) representing the progression of cellular transcriptional states across LSAT, BCRL and OSAT conditions.** Cells are displayed in a shared transcriptional embedding and colored according to inferred latent time, from early (dark colors) to late (yellow colors) dynamic transcriptional states.

**B. Latent time distribution between immune cell groups from LSAT, BCRL and OSAT conditions.** Each violin shows the latent time distribution for cell group split by each studied condition

**Supplementary figure 4.**

**Representative image of the ROIs and cell segmentation used for the OX40 and OX40L quantification in Visiopharm.** The sample’s dermal ROI’s are annotated using green and blue solid outlines, and its adipose tissue regions using a red solid outline. Excluded areas are outlined with a gray solid outline. Cells annotated with blue are categorized as negative, while those annotated with yellow, orange and red are categorized as positive and grouped by increasing intensity.

## Notes

“The authors have declared that no conflict of interest exists.”

https://www.ncbi.nlm.nih.gov/geo/query/acc.cgi?acc=GSE346502

### Competing Interest Statement

The authors have declared no competing interest.

