## Supplementary information 1 for "Single-Cell Profiling Reveals Innate Lymphoid Cells and OX40 Activation in Breast Cancer-Related Lymphedema"

**Characterization of the cellular diversity and expression signatures of each identified cluster**

Individual clusters within the $CD4^+$ T cell group included, **c0** ($CD4^+$ T-a) marked by the expression of *NIBAN1*, *ANK3*, and *ITK*; **c1** ($CD4^+$ T-b) characterized by *RGCC*, *CD2*, and *ANXA1*; **c5** (Early Activated $CD4^+$ T) marked by markers *CCR6*, *LTB* (lymphotoxin-β) and *PTPN13;* and **c16** (Naïve $CD4^+$ T) marked by naïve markers *SELL*, *FHIT*, and *CCR7 (22,29).*

Mature NK cells in **c3** were characterized by their high expression of *cytotoxic* effector granules and receptor *genes including, PRF1*, *SPON2* and *CST7*; while those in **c7** were marked by *GNLY*, *KLRD1* and *KLRF1* (30) (Figures 2B, D). These mature NK cells categorized as NCAM or $CD56^{dim}$ cells (31). Resident and $CD56^{bright}$ NKs in **c12** (Figures 2B, D), in turn, showed lower cytotoxicity potential (*PRF1* and *GNLY*) but high levels of Th1 pro-inflammatory genes, including *XCL1*, *XCL2*, *IL12RB2*, *REL*, *NFKB1* and elevated NK specific tissue residency genes, *KLRC1*, *KLRC3* and *NCAM1* (21,30). $CD8^+$ T cells (Figures 2B, D) in **c2** presented signs of an activated state expressing effector genes, *GZMK* and *KLRK1*, and chemokine genes, *CCL5* and *CCL4*, whereas CD8+ T cells in **c20**, presented a more naïve-memory like signature with expression of quiescence factors, such as *LEF1* and *BACH2*, and effector receptor and factor genes, *KLRC2*, *KLRC3*, *IL21R* and *IFNG-AS1 (32)* (Figures 2B, D).

ATMs in **c6** were identified based on their high expression levels of classical macrophage markers, including *CD14* and *LYZ*, proinflammatory genes, *C5AR1*, *KYNU*, and tissue residency markers, *VCAM*, *CD163* and *PSAP (33,34)* (Figures 2B, D). In **c8**, we identified intermediate or M1/M2 macrophages characterized by high expression of CD14 and LYZ and low expression of tissue residency markers *CD163* and *MRC1* (Figures 2B, D). These cells also exhibited dual high expression of both pro-inflammatory markers, including several MHC class II genes and anti-inflammatory/immune-regulatory genes such as *CLEC10A*, *FCER1A*, and *IL1R2 (35,36)*. NCMs in **c14** exhibited very low *CD14* levels and strong *FCGR3A* expression (Figures 2B, D) as well as high expression of inducible markers associated with this subset, including *LST1*, *AIF1*, and *MS4A7(37).* PVMs in **c15** were characterized by high expression of immunoregulatory genes such as *SELENOP* and *C1QA* along with perivascular markers *CD163*, *MRC1*, and *LYVE1 (38).*

NK/T cells in **c4** expressed a potent activating and cytotoxic gene signature marked by *CD3G* and *TGFBR3* and high expression of several granzymes including, *GZMH*, *GZMB*, *GNLY*, *PRF1*, *NKG7* (Figures 2B, D). Similarly, NK/T cells in **c10** showed high expression of same cytotoxic genes and enhanced proinflammatory activity with expression of *CCL4*, *CCL5* and *IFNG (23,30)* (Figures 2B, D). Proliferating NK/T cells in **c18**, on the other hand, showed a clear signature associated with regulation of cell cycle and proliferation, expressing *STMN1*, *TUBB, MKI67* and *TYMS (23,39)* (Figures 2B, D). Specialized T cells included two different clusters. In **c11**, MAIT cells expressed *RORA*, *SLC4A10*, *PHACTHR2* and *KLRB1* and cytotoxic markers such as *CD8* and *GZMK (25)* and Treg cells in **c9** characterized by low expression of *FOXP3* and *IL7R*, but high expression of regulatory identity markers, including immunosuppressive genes *CTLA4*, *IKZF2*, and *IL2RA*, as well as differentiation and maintenance genes such as *TOX*, *ICOS*, *TIGIT*, *TNFRSF9* and *TNFRSF18 (40,41)* (Figures 2B, D).

B cells showed two expression signatures, mature B cells in **c13** expressing high levels of B cells activation genes *BANK1*, *CD79A*, *MS4A1* and the B cell antigen receptors *IGHM* and *IGHD (26)*; and plasma B cells in **c23**, showing high levels of B cell differentiation markers including, *MZB1*, *JCHAIN and TNFRSF17*, and antigen-specific *antibodies, such as IGKC and IGHA2 (42)* (Figures 2B, D). Similar to these, two clear clusters of DCs, **c21** and **c22**, were annotated based on expression of previously identified markers for cDC, including *THBD*

(*BATF3*), *WDFY4* and *CLEC9A*; and for plasmacytoid DC, such as *TCF4*, *CLEC4C* (*BDCA2*) and *IRF4 (27)* (Figures 2B, D).

The expression signature for ILCs in **c17** was featured by the lack of hematopoietic cell lineage surface markers, *CD3*, *CD14*, *CD16*, *CD19*, *CD20* and *CD56* and the high expression of intracellular markers *GATA3* and *RORC*, as well as the receptors *KIT* (cKIT), *KLRB1* and *IL7R (43-45)* (Figures 2B, D). The smallest cluster identified (**c19**) exhibited genes related to fibroblast activation and remodeling function, such as *GSN*, *IGFBP7* and *TIMP3*, *CAV1*, *ADIRF*, respectively (46) (Figures 2B, D). Although fibroblasts are not commonly defined as CD45+ cells, cells in c9 expressed enough levels of CD45 (Figure 2D) to be sorted.

**Supplementary table 1.** Demographic characteristics of LSAT, OSAT and BSAT controls

LSAT= Lean Subcutaneous Adipose Tissue

OSAT= Obese Subcutaneous Adipose Tissue

BSAT= Breast Subcutaneous Adipose Tissue

BMI= Body Mass Index

| **ID** | **Age** | **BMI (Kg/m$^2$)** |
|---|---|---|
| LSAT-1 | 39 | 20,5 |
| LSAT-2 | 32 | 24,8 |
| LSAT-3 | 43 | 16,9 |
| OSAT-1 | 39 | 33,4 |
| OSAT-2 | 36 | 30,0 |
| OSAT-3 | 35 | 31,6 |
| BSAT-1 | 41 | 25,2 |
| BSAT-2 | 30 | 28,2 |
| BSAT-3 | 54 | 26,4 |
| BSAT-4 | 58 | 20,9 |
| BSAT-5 | 63 | 26,1 |
| BSAT-6 | 31 | 29,4 |
| BSAT-7 | 52 | 24,2 |
| BSAT-8 | 68 | 29,1 |
| BSAT-9 | 56 | 27,5 |
| BSAT-10 | 24 | 27,3 |
| BSAT-11 | 41 | 26,1 |
| BSAT-12 | 50 | 32,7 |
| BSAT-13 | 58 | 24,9 |
| BSAT-14 | 45 | 28,3 |
| BSAT-15 | 76 | 30,1 |
| BSAT-16 | 41 | 27,9 |

**Supplementary table 2.**

Table of the cluster marker genes for each cell type identified. Genes with an adjusted p-value (Benjamini–Hochberg correction).

| row.name | p_val | avg_log2F | pct.1 | pct.2 | p_val_ | cluster | gene |
|---|---|---|---|---|---|---|---|
| IL7R | 0 | 1,26554873 | 0,913 | 0,449 | 0 | CD4+_T-a | IL7R |
| BACH2 | 0 | 1,50701245 | 0,771 | 0,353 | 0 | CD4+_T-a | BACH2 |
| PBX4 | 0 | 1,72543539 | 0,631 | 0,224 | 0 | CD4+_T-a | PBX4 |
| ANK3 | 0 | 2,48308725 | 0,536 | 0,137 | 0 | CD4+_T-a | ANK3 |
| PCNX1 | 0 | 1,81137045 | 0,654 | 0,273 | 0 | CD4+_T-a | PCNX1 |
| ITK | 0 | 1,5719091 | 0,693 | 0,318 | 0 | CD4+_T-a | ITK |
| NIBAN1 | 0 | 1,31943733 | 0,836 | 0,464 | 0 | CD4+_T-a | NIBAN1 |
| MAF | 0 | 1,69412475 | 0,616 | 0,254 | 0 | CD4+_T-a | MAF |
| RNF19A | 0 | 1,42446786 | 0,859 | 0,499 | 0 | CD4+_T-a | RNF19A |
| BCL11B | 0 | 1,20980609 | 0,687 | 0,337 | 0 | CD4+_T-a | BCL11B |
| RBMS1 | 0 | 1,47027804 | 0,605 | 0,26 | 0 | CD4+_T-a | RBMS1 |
| EML4 | 0 | 1,09103382 | 0,821 | 0,481 | 0 | CD4+_T-a | EML4 |
| FKBP5 | 0 | 1,36141518 | 0,836 | 0,498 | 0 | CD4+_T-a | FKBP5 |
| ARID5B | 0 | 1,13167182 | 0,737 | 0,402 | 0 | CD4+_T-a | ARID5B |
| INPP4B | 0 | 2,07875302 | 0,487 | 0,153 | 0 | CD4+_T-a | INPP4B |
| CD2 | 0 | 1,22773959 | 0,631 | 0,306 | 0 | CD4+_T-a | CD2 |
| CRYBG1 | 0 | 1,37148886 | 0,819 | 0,496 | 0 | CD4+_T-a | CRYBG1 |
| BICDL1 | 0 | 1,50929062 | 0,532 | 0,211 | 0 | CD4+_T-a | BICDL1 |
| RHOH | 0 | 1,19598121 | 0,642 | 0,33 | 0 | CD4+_T-a | RHOH |
| NR3C1 | 0 | 1,39205075 | 0,712 | 0,402 | 0 | CD4+_T-a | NR3C1 |
| ZNF831 | 0 | 1,49033306 | 0,52 | 0,214 | 0 | CD4+_T-a | ZNF831 |
| HERC1 | 0 | 1,30401199 | 0,622 | 0,318 | 0 | CD4+_T-a | HERC1 |
| DOCK10 | 0 | 1,17025419 | 0,6 | 0,31 | 0 | CD4+_T-a | DOCK10 |
| RORA | 0 | 0,95827684 | 0,692 | 0,403 | 0 | CD4+_T-a | RORA |
| CDC14A | 0 | 1,50604961 | 0,522 | 0,235 | 0 | CD4+_T-a | CDC14A |
| ANKRD28 | 0 | 1,45463833 | 0,616 | 0,329 | 0 | CD4+_T-a | ANKRD28 |
| ATXN1 | 0 | 1,08968633 | 0,642 | 0,356 | 0 | CD4+_T-a | ATXN1 |
| LEPROTL1 | 0 | 0,91606469 | 0,705 | 0,419 | 0 | CD4+_T-a | LEPROTL1 |
| ZC3HAV1 | 0 | 1,08629782 | 0,688 | 0,404 | 0 | CD4+_T-a | ZC3HAV1 |
| ANKRD12 | 0 | 1,1892447 | 0,772 | 0,491 | 0 | CD4+_T-a | ANKRD12 |
| FAM107B | 0 | 1,0499967 | 0,694 | 0,418 | 0 | CD4+_T-a | FAM107B |
| CYTH1 | 0 | 1,09135638 | 0,647 | 0,374 | 0 | CD4+_T-a | CYTH1 |
| CD6 | 0 | 1,53224871 | 0,433 | 0,163 | 0 | CD4+_T-a | CD6 |
| SMCHD1 | 0 | 1,02023156 | 0,892 | 0,623 | 0 | CD4+_T-a | SMCHD1 |
| PARP8 | 0 | 0,84276044 | 0,824 | 0,556 | 0 | CD4+_T-a | PARP8 |
| LINC-PINT | 0 | 1,05834376 | 0,58 | 0,313 | 0 | CD4+_T-a | LINC-PINT |
| AP001011.1 | 0 | 1,10471727 | 0,575 | 0,308 | 0 | CD4+_T-a | AP001011.1 |
| SAMSN1 | 0 | 0,52336594 | 0,824 | 0,557 | 0 | CD4+_T-a | SAMSN1 |
| ARHGAP15 | 0 | 1,03345811 | 0,745 | 0,48 | 0 | CD4+_T-a | ARHGAP15 |
| TNFSF8 | 0 | 1,73206604 | 0,399 | 0,134 | 0 | CD4+_T-a | TNFSF8 |
| RCAN3 | 0 | 1,45483685 | 0,425 | 0,161 | 0 | CD4+_T-a | RCAN3 |
| HNRNPLL | 0 | 1,69274345 | 0,4 | 0,136 | 0 | CD4+_T-a | HNRNPLL |
| RGCC | 0 | 0,74582412 | 0,692 | 0,428 | 0 | CD4+_T-a | RGCC |
| CNOT6L | 0 | 0,79376312 | 0,923 | 0,661 | 0 | CD4+_T-a | CNOT6L |
| LINC00513 | 0 | 1,042703 | 0,635 | 0,373 | 0 | CD4+_T-a | LINC00513 |
| SESN3 | 0 | 2,40754483 | 0,362 | 0,1 | 0 | CD4+_T-a | SESN3 |
| CDC42SE2 | 0 | 0,89519899 | 0,799 | 0,539 | 0 | CD4+_T-a | CDC42SE2 |
| SYNE2 | 0 | 0,86323834 | 0,687 | 0,429 | 0 | CD4+_T-a | SYNE2 |
| OSBPL8 | 0 | 1,10291412 | 0,636 | 0,378 | 0 | CD4+_T-a | OSBPL8 |
| PPP1CB | 0 | 0,9383046 | 0,724 | 0,466 | 0 | CD4+_T-a | PPP1CB |
| SLC2A3 | 0 | 1,09021684 | 0,613 | 0,356 | 0 | CD4+_T-a | SLC2A3 |
| PGAP1 | 0 | 2,39781249 | 0,349 | 0,095 | 0 | CD4+_T-a | PGAP1 |
| ABCC1 | 0 | 1,79996876 | 0,383 | 0,129 | 0 | CD4+_T-a | ABCC1 |
| STK4 | 0 | 0,87970539 | 0,788 | 0,536 | 0 | CD4+_T-a | STK4 |
| MAML2 | 0 | 1,26260455 | 0,544 | 0,294 | 0 | CD4+_T-a | MAML2 |
| KAT6A | 0 | 1,0363217 | 0,549 | 0,3 | 0 | CD4+_T-a | KAT6A |
| TC2N | 0 | 1,2660242 | 0,459 | 0,211 | 0 | CD4+_T-a | TC2N |
| EZR | 0 | 0,84106846 | 0,841 | 0,594 | 0 | CD4+_T-a | EZR |
| FOXP1 | 0 | 0,96048149 | 0,689 | 0,444 | 0 | CD4+_T-a | FOXP1 |
| AC016831.7 | 0 | 1,1882878 | 0,501 | 0,257 | 0 | CD4+_T-a | AC016831.7 |
| ANXA1 | 0 | 0,85606137 | 0,874 | 0,63 | 0 | CD4+_T-a | ANXA1 |
| CD96 | 0 | 1,06833695 | 0,579 | 0,335 | 0 | CD4+_T-a | CD96 |
| AKT3 | 0 | 1,14272429 | 0,554 | 0,311 | 0 | CD4+_T-a | AKT3 |
| FOXO1 | 0 | 0,98201903 | 0,558 | 0,316 | 0 | CD4+_T-a | FOXO1 |
| ZEB1 | 0 | 1,23291523 | 0,448 | 0,207 | 0 | CD4+_T-a | ZEB1 |
| PTGER4 | 0 | 1,16531109 | 0,479 | 0,238 | 0 | CD4+_T-a | PTGER4 |
| CASK | 0 | 1,43547898 | 0,414 | 0,173 | 0 | CD4+_T-a | CASK |
| ANKRD44 | 0 | 0,96922312 | 0,583 | 0,344 | 0 | CD4+_T-a | ANKRD44 |
| SARAF | 0 | 0,81536468 | 0,909 | 0,67 | 0 | CD4+_T-a | SARAF |
| ETS1 | 0 | 0,71649759 | 0,818 | 0,579 | 0 | CD4+_T-a | ETS1 |
| TP53BP2 | 0 | 1,55972668 | 0,392 | 0,154 | 0 | CD4+_T-a | TP53BP2 |
| STAT4 | 0 | 0,73677867 | 0,827 | 0,589 | 0 | CD4+_T-a | STAT4 |
| GSPT1 | 0 | 1,05250241 | 0,509 | 0,271 | 0 | CD4+_T-a | GSPT1 |
| LMNA | 0 | 0,69207089 | 0,715 | 0,478 | 0 | CD4+_T-a | LMNA |

| row.names | p_val | avg_log2FC | pct.1 | pct.2 | p_val_a | cluster | gene |
|---|---|---|---|---|---|---|---|
| IL7R1 | 0 | 1,24967085 | 0,902 | 0,459 | 0 | CD4+_T-b | IL7R |
| RGCC1 | 0 | 1,29536919 | 0,76 | 0,423 | 0 | CD4+_T-b | RGCC |
| CD21 | 0 | 1,26216769 | 0,633 | 0,312 | 0 | CD4+_T-b | CD2 |
| MAF2 | 0 | 1,40992256 | 0,565 | 0,268 | 0 | CD4+_T-b | MAF |
| LMNA1 | 0 | 0,93941557 | 0,761 | 0,475 | 0 | CD4+_T-b | LMNA |
| ANXA11 | 0 | 1,23960606 | 0,914 | 0,629 | 0 | CD4+_T-b | ANXA1 |
| LEPROTL11 | 0 | 0,91796485 | 0,689 | 0,426 | 0 | CD4+_T-b | LEPROTL1 |
| S100A10 | 0 | 0,78641578 | 0,872 | 0,617 | 0 | CD4+_T-b | S100A10 |
| JUNB1 | 0 | 1,16320995 | 0,873 | 0,62 | 0 | CD4+_T-b | JUNB |
| ITM2A1 | 0 | 1,42322837 | 0,437 | 0,192 | 0 | CD4+_T-b | ITM2A |
| CD40LG1 | 0 | 2,30113475 | 0,309 | 0,071 | 0 | CD4+_T-b | CD40LG |
| CRIP1 | 0 | 0,84634175 | 0,867 | 0,63 | 0 | CD4+_T-b | CRIP1 |
| IL32 | 0 | 0,66867216 | 0,712 | 0,476 | 0 | CD4+_T-b | IL32 |
| FOS | 0 | 0,90685945 | 0,71 | 0,477 | 0 | CD4+_T-b | FOS |
| TOMM7 | 0 | 0,80320157 | 0,76 | 0,529 | 0 | CD4+_T-b | TOMM7 |
| RPL36AL | 0 | 0,77111304 | 0,834 | 0,608 | 0 | CD4+_T-b | RPL36AL |
| SARAF1 | 0 | 0,87058432 | 0,899 | 0,676 | 0 | CD4+_T-b | SARAF |
| RNF19A1 | 0 | 0,59395026 | 0,742 | 0,523 | 0 | CD4+_T-b | RNF19A |
| ARID5B1 | 0 | 0,69501718 | 0,64 | 0,423 | 0 | CD4+_T-b | ARID5B |
| ZFP36 | 0 | 1,12598371 | 0,944 | 0,727 | 0 | CD4+_T-b | ZFP36 |
| MCL11 | 0 | 0,82636558 | 0,775 | 0,561 | 0 | CD4+_T-b | MCL1 |
| KLF61 | 0 | 0,84232931 | 0,717 | 0,505 | 0 | CD4+_T-b | KLF6 |
| ZFP36L21 | 0 | 1,00725678 | 0,952 | 0,743 | 0 | CD4+_T-b | ZFP36L2 |
| RGS11 | 0 | 0,86967879 | 0,534 | 0,329 | 0 | CD4+_T-b | RGS1 |
| PTGER41 | 0 | 1,06001511 | 0,45 | 0,247 | 0 | CD4+_T-b | PTGER4 |
| TSC22D3 | 0 | 1,09891859 | 0,943 | 0,741 | 0 | CD4+_T-b | TSC22D3 |
| FXYD5 | 0 | 0,78789476 | 0,57 | 0,371 | 0 | CD4+_T-b | FXYD5 |
| CD52 | 0 | 0,79530149 | 0,663 | 0,467 | 0 | CD4+_T-b | CD52 |
| TUBB4B | 0 | 0,9455327 | 0,495 | 0,3 | 0 | CD4+_T-b | TUBB4B |
| TXNIP1 | 0 | 0,8692521 | 0,765 | 0,578 | 0 | CD4+_T-b | TXNIP |
| UBE2S | 0 | 1,00202346 | 0,424 | 0,237 | 0 | CD4+_T-b | UBE2S |
| CXCR4 | 0 | 0,99469176 | 0,956 | 0,769 | 0 | CD4+_T-b | CXCR4 |
| SOCS11 | 0 | 1,14528008 | 0,383 | 0,202 | 0 | CD4+_T-b | SOCS1 |
| PHLDA11 | 0 | 1,32560211 | 0,323 | 0,143 | 0 | CD4+_T-b | PHLDA1 |
| CD99 | 0 | 0,74254719 | 0,658 | 0,479 | 0 | CD4+_T-b | CD99 |
| TRAC | 0 | 1,0038301 | 0,368 | 0,189 | 0 | CD4+_T-b | TRAC |
| PLP2 | 0 | 1,12693361 | 0,345 | 0,168 | 0 | CD4+_T-b | PLP2 |
| EEF1D | 0 | 0,60505102 | 0,784 | 0,615 | 0 | CD4+_T-b | EEF1D |
| TNFSF81 | 0 | 1,11545579 | 0,32 | 0,151 | 0 | CD4+_T-b | TNFSF8 |
| RPS5 | 0 | 0,59526528 | 0,92 | 0,754 | 0 | CD4+_T-b | RPS5 |
| RPSA | 0 | 0,58758634 | 0,908 | 0,746 | 0 | CD4+_T-b | RPSA |
| TRAT11 | 0 | 1,44360118 | 0,252 | 0,094 | 0 | CD4+_T-b | TRAT1 |
| RPL36A | 0 | 0,67293897 | 0,953 | 0,797 | 0 | CD4+_T-b | RPL36A |
| RPS26 | 0 | 0,86048225 | 0,874 | 0,731 | 0 | CD4+_T-b | RPS26 |
| JUND | 0 | 0,99762314 | 0,976 | 0,836 | 0 | CD4+_T-b | JUND |
| CLU1 | 0 | 1,91723905 | 0,182 | 0,042 | 0 | CD4+_T-b | CLU |
| RPL10A | 0 | 0,55009694 | 0,937 | 0,805 | 0 | CD4+_T-b | RPL10A |
| RPL38 | 0 | 0,67962673 | 0,968 | 0,836 | 0 | CD4+_T-b | RPL38 |
| UBC | 0 | 0,6084301 | 0,912 | 0,786 | 0 | CD4+_T-b | UBC |
| RPL24 | 0 | 0,59937028 | 0,967 | 0,849 | 0 | CD4+_T-b | RPL24 |
| RPL36 | 0 | 0,71167968 | 0,986 | 0,87 | 0 | CD4+_T-b | RPL36 |
| RPL35 | 0 | 0,59605121 | 0,952 | 0,839 | 0 | CD4+_T-b | RPL35 |
| RPS16 | 0 | 0,63975551 | 0,972 | 0,86 | 0 | CD4+_T-b | RPS16 |
| RPS25 | 0 | 0,76129734 | 0,989 | 0,879 | 0 | CD4+_T-b | RPS25 |
| BTG1 | 0 | 0,64550288 | 0,955 | 0,848 | 0 | CD4+_T-b | BTG1 |
| PDCD11 | 0 | 1,75110221 | 0,153 | 0,047 | 0 | CD4+_T-b | PDCD1 |
| RPL14 | 0 | 0,63405786 | 0,982 | 0,879 | 0 | CD4+_T-b | RPL14 |
| RPL23A | 0 | 0,63640152 | 0,984 | 0,883 | 0 | CD4+_T-b | RPL23A |
| H3F3B | 0 | 0,66561163 | 0,945 | 0,844 | 0 | CD4+_T-b | H3F3B |
| RPL35A | 0 | 0,65633857 | 0,989 | 0,889 | 0 | CD4+_T-b | RPL35A |
| RPS7 | 0 | 0,59159213 | 0,986 | 0,892 | 0 | CD4+_T-b | RPS7 |
| RPL9 | 0 | 0,51967137 | 0,976 | 0,883 | 0 | CD4+_T-b | RPL9 |
| RPL7A | 0 | 0,51861639 | 0,979 | 0,89 | 0 | CD4+_T-b | RPL7A |
| RPS3 | 0 | 0,62020077 | 0,993 | 0,905 | 0 | CD4+_T-b | RPS3 |
| RPS21 | 0 | 0,71280605 | 0,996 | 0,911 | 0 | CD4+_T-b | RPS21 |
| RPL17 | 0 | 0,50254118 | 0,983 | 0,9 | 0 | CD4+_T-b | RPL17 |
| RPL12 | 0 | 0,57651406 | 0,99 | 0,909 | 0 | CD4+_T-b | RPL12 |
| FAU | 0 | 0,6031567 | 0,991 | 0,912 | 0 | CD4+_T-b | FAU |
| RPL18A | 0 | 0,55571269 | 0,987 | 0,91 | 0 | CD4+_T-b | RPL18A |
| RPLP2 | 0 | 0,58328031 | 0,991 | 0,917 | 0 | CD4+_T-b | RPLP2 |
| RPS4X | 0 | 0,54937673 | 0,993 | 0,919 | 0 | CD4+_T-b | RPS4X |
| RPL19 | 0 | 0,59285015 | 0,995 | 0,925 | 0 | CD4+_T-b | RPL19 |
| RPS15 | 0 | 0,52772604 | 0,99 | 0,922 | 0 | CD4+_T-b | RPS15 |
| RPS19 | 0 | 0,61418644 | 0,996 | 0,929 | 0 | CD4+_T-b | RPS19 |

| row.names | p_val | avg_log2FC | pct.1 | pct.2 | p_val_a | cluster | gene |
|---|---|---|---|---|---|---|---|
| GZMK1 | 0 | 2,3995695 | 0,736 | 0,192 | 0 | CD8+-T | GZMK |
| CCL5 | 0 | 1,54042965 | 0,96 | 0,495 | 0 | CD8+-T | CCL5 |
| CD8A | 0 | 2,4954318 | 0,542 | 0,104 | 0 | CD8+-T | CD8A |
| KLRK1 | 0 | 1,96424171 | 0,538 | 0,153 | 0 | CD8+-T | KLRK1 |
| CCL4 | 0 | 1,19313628 | 0,691 | 0,355 | 0 | CD8+-T | CCL4 |
| CRTAM | 0 | 2,93532148 | 0,375 | 0,053 | 0 | CD8+-T | CRTAM |
| TUBA4A1 | 0 | 1,34913071 | 0,668 | 0,349 | 0 | CD8+-T | TUBA4A |
| RUNX3 | 0 | 1,04378253 | 0,741 | 0,433 | 0 | CD8+-T | RUNX3 |
| CD8B | 0 | 2,87721493 | 0,355 | 0,048 | 0 | CD8+-T | CD8B |
| DUSP2 | 0 | 1,0464037 | 0,813 | 0,514 | 0 | CD8+-T | DUSP2 |
| RNF19A2 | 0 | 0,87171295 | 0,807 | 0,515 | 0 | CD8+-T | RNF19A |
| CST7 | 0 | 0,54402925 | 0,706 | 0,427 | 0 | CD8+-T | CST7 |
| LYST | 0 | 1,63459887 | 0,487 | 0,21 | 0 | CD8+-T | LYST |
| PIK3R1 | 0 | 0,82039134 | 0,745 | 0,477 | 0 | CD8+-T | PIK3R1 |
| BICDL11 | 0 | 1,4406643 | 0,492 | 0,224 | 0 | CD8+-T | BICDL1 |
| PPP2R5C | 0 | 0,69647719 | 0,822 | 0,575 | 0 | CD8+-T | PPP2R5C |
| CBLB1 | 0 | 0,92549987 | 0,762 | 0,517 | 0 | CD8+-T | CBLB |
| ITGA4 | 0 | 1,10465876 | 0,542 | 0,298 | 0 | CD8+-T | ITGA4 |
| BCL11B1 | 0 | 0,95618058 | 0,6 | 0,358 | 0 | CD8+-T | BCL11B |
| NIBAN11 | 0 | 0,75819754 | 0,726 | 0,489 | 0 | CD8+-T | NIBAN1 |
| YBX3 | 0 | 1,39327804 | 0,363 | 0,141 | 0 | CD8+-T | YBX3 |
| AKNA | 0 | 0,75427061 | 0,546 | 0,326 | 0 | CD8+-T | AKNA |
| PARP81 | 0 | 0,65341359 | 0,786 | 0,568 | 0 | CD8+-T | PARP8 |
| CEMIP2 | 0 | 0,8681493 | 0,784 | 0,566 | 0 | CD8+-T | CEMIP2 |
| TERF2IP | 0 | 1,16300441 | 0,435 | 0,224 | 0 | CD8+-T | TERF2IP |
| FYN1 | 0 | 0,63010496 | 0,898 | 0,69 | 0 | CD8+-T | FYN |
| SRSF2 | 0 | 0,81625325 | 0,604 | 0,397 | 0 | CD8+-T | SRSF2 |
| WNK11 | 0 | 1,00281686 | 0,475 | 0,271 | 0 | CD8+-T | WNK1 |
| PTPN22 | 0 | 0,89625489 | 0,525 | 0,325 | 0 | CD8+-T | PTPN22 |
| THEMIS1 | 0 | 1,37867354 | 0,338 | 0,149 | 0 | CD8+-T | THEMIS |
| CCL4L2 | 0 | 1,90696285 | 0,296 | 0,107 | 0 | CD8+-T | CCL4L2 |
| TGFB1 | 0 | 0,69430706 | 0,702 | 0,514 | 0 | CD8+-T | TGFB1 |
| TC2N1 | 0 | 1,0125606 | 0,411 | 0,224 | 0 | CD8+-T | TC2N |
| GALNT11 | 0 | 1,28943367 | 0,327 | 0,142 | 0 | CD8+-T | GALNT11 |
| SH2D1A | 0 | 1,68795973 | 0,283 | 0,098 | 0 | CD8+-T | SH2D1A |
| DTHD1 | 0 | 1,52289623 | 0,288 | 0,107 | 0 | CD8+-T | DTHD1 |
| IL21R | 0 | 1,48490934 | 0,28 | 0,107 | 0 | CD8+-T | IL21R |
| XCL2 | 0 | 1,32667579 | 0,271 | 0,098 | 0 | CD8+-T | XCL2 |
| TRGC2 | 0 | 1,53836363 | 0,256 | 0,088 | 0 | CD8+-T | TRGC2 |
| GGA2 | 0 | 1,52252091 | 0,254 | 0,096 | 0 | CD8+-T | GGA2 |
| LYAR | 0 | 1,27257734 | 0,282 | 0,124 | 0 | CD8+-T | LYAR |
| PTPRC | 0 | 0,68288477 | 0,958 | 0,801 | 0 | CD8+-T | PTPRC |
| CXCR41 | 0 | 0,55697549 | 0,922 | 0,775 | 0 | CD8+-T | CXCR4 |
| WHRN | 0 | 1,60634123 | 0,196 | 0,069 | 0 | CD8+-T | WHRN |
| TNIP3 | 0 | 2,47081441 | 0,15 | 0,025 | 0 | CD8+-T | TNIP3 |
| TNFSF9 | 0 | 1,70574439 | 0,179 | 0,056 | 0 | CD8+-T | TNFSF9 |
| TNFRSF9 | 0 | 2,02442667 | 0,15 | 0,037 | 0 | CD8+-T | TNFRSF9 |
| AC243829.4 | 0 | 2,34975946 | 0,129 | 0,026 | 0 | CD8+-T | AC243829.4 |
| LAG3 | 0 | 2,11301148 | 0,129 | 0,031 | 0 | CD8+-T | LAG3 |
| FXYD21 | 0 | 2,30472179 | 0,112 | 0,024 | 0 | CD8+-T | FXYD2 |
| DRAIC | 0 | 2,99804819 | 0,071 | 0,009 | 0 | CD8+-T | DRAIC |
| LINC02446 | 6E-305 | 2,26594066 | 0,101 | 0,023 | 2E-300 | CD8+-T | LINC02446 |
| ATXN11 | 3E-304 | 0,8138523 | 0,571 | 0,373 | 8E-300 | CD8+-T | ATXN1 |
| HERPUD21 | 6E-296 | 1,04074678 | 0,385 | 0,209 | 1E-291 | CD8+-T | HERPUD2 |
| GABARAPL1 | 2E-292 | 0,95721327 | 0,409 | 0,228 | 5E-288 | CD8+-T | GABARAPL1 |
| SAMD3 | 8E-291 | 0,89142089 | 0,402 | 0,217 | 2E-286 | CD8+-T | SAMD3 |
| SRSF7 | 6E-272 | 0,64161911 | 0,692 | 0,51 | 2E-267 | CD8+-T | SRSF7 |
| PARD6G | 3E-268 | 1,51661268 | 0,161 | 0,056 | 8E-264 | CD8+-T | PARD6G |
| MCOLN2 | 6E-264 | 1,44277738 | 0,219 | 0,093 | 1E-259 | CD8+-T | MCOLN2 |
| LINC01871 | 2E-259 | 0,95191589 | 0,389 | 0,219 | 4E-255 | CD8+-T | LINC01871 |
| LEPROTL12 | 2E-258 | 0,60476426 | 0,637 | 0,435 | 4E-254 | CD8+-T | LEPROTL1 |
| UBE2S1 | 4E-244 | 0,87181697 | 0,408 | 0,24 | 1E-239 | CD8+-T | UBE2S |
| GLCCI1 | 5E-240 | 0,86980128 | 0,428 | 0,259 | 1E-235 | CD8+-T | GLCCI1 |
| RNF1251 | 4E-232 | 0,61444099 | 0,648 | 0,471 | 9E-228 | CD8+-T | RNF125 |
| PPP1R16B1 | 5E-225 | 0,72796522 | 0,484 | 0,312 | 1E-220 | CD8+-T | PPP1R16B |
| GTDC1 | 7E-225 | 1,04972124 | 0,281 | 0,143 | 2E-220 | CD8+-T | GTDC1 |
| RHOH1 | 3E-224 | 0,70744264 | 0,529 | 0,353 | 6E-220 | CD8+-T | RHOH |
| CMC1 | 3E-221 | 0,98809424 | 0,3 | 0,159 | 9E-217 | CD8+-T | CMC1 |
| JMJD6 | 2E-216 | 1,15523315 | 0,22 | 0,101 | 6E-212 | CD8+-T | JMJD6 |
| ADGRE5 | 2E-211 | 0,63309955 | 0,589 | 0,426 | 5E-207 | CD8+-T | ADGRE5 |
| STYK1 | 2E-204 | 2,9985431 | 0,039 | 0,005 | 6E-200 | CD8+-T | STYK1 |
| LITAF | 7E-203 | 0,56994888 | 0,582 | 0,408 | 2E-198 | CD8+-T | LITAF |
| CDC42SE21 | 1E-201 | 0,50027235 | 0,717 | 0,557 | 3E-197 | CD8+-T | CDC42SE2 |
| ATP8A1 | 5E-198 | 0,77041094 | 0,359 | 0,208 | 1E-193 | CD8+-T | ATP8A1 |

| row.names | p_val | avg_log2FC | pct.1 | pct.2 | p_val_ad | cluster | gene |
|---|---|---|---|---|---|---|---|
| GZMB | 0 | 3,0432461 | 0,946 | 0,214 | 0 | Mature NK-a | GZMB |
| GNLY | 0 | 2,69661049 | 0,951 | 0,234 | 0 | Mature NK-a | GNLY |
| PRF1 | 0 | 2,91769616 | 0,865 | 0,177 | 0 | Mature NK-a | PRF1 |
| SPON2 | 0 | 3,04692584 | 0,773 | 0,126 | 0 | Mature NK-a | SPON2 |
| FGFBP2 | 0 | 3,11557204 | 0,757 | 0,116 | 0 | Mature NK-a | FGFBP2 |
| KLRD1 | 0 | 1,96179638 | 0,852 | 0,228 | 0 | Mature NK-a | KLRD1 |
| NKG7 | 0 | 2,29502307 | 0,974 | 0,363 | 0 | Mature NK-a | NKG7 |
| KLRF1 | 0 | 3,0613156 | 0,674 | 0,082 | 0 | Mature NK-a | KLRF1 |
| AOAH | 0 | 2,32154754 | 0,85 | 0,302 | 0 | Mature NK-a | AOAH |
| CD247 | 0 | 2,18321736 | 0,939 | 0,394 | 0 | Mature NK-a | CD247 |
| C1orf21 | 0 | 2,60871517 | 0,671 | 0,134 | 0 | Mature NK-a | C1orf21 |
| FCGR3A | 0 | 2,79462131 | 0,631 | 0,098 | 0 | Mature NK-a | FCGR3A |
| METRNL | 0 | 2,09615115 | 0,894 | 0,365 | 0 | Mature NK-a | METRNL |
| CD7 | 0 | 2,03550017 | 0,779 | 0,255 | 0 | Mature NK-a | CD7 |
| CTSW | 0 | 2,07171347 | 0,747 | 0,226 | 0 | Mature NK-a | CTSW |
| CST71 | 0 | 1,98675688 | 0,925 | 0,417 | 0 | Mature NK-a | CST7 |
| CARD11 | 0 | 2,58803635 | 0,653 | 0,165 | 0 | Mature NK-a | CARD11 |
| MCTP2 | 0 | 2,54427464 | 0,62 | 0,135 | 0 | Mature NK-a | MCTP2 |
| TYROBP | 0 | 1,25625374 | 0,725 | 0,248 | 0 | Mature NK-a | TYROBP |
| MYOM2 | 0 | 3,7340065 | 0,517 | 0,053 | 0 | Mature NK-a | MYOM2 |
| TFDP2 | 0 | 2,37110249 | 0,593 | 0,162 | 0 | Mature NK-a | TFDP2 |
| SLA2 | 0 | 2,47116525 | 0,546 | 0,117 | 0 | Mature NK-a | SLA2 |
| PLAC8 | 0 | 2,12198367 | 0,572 | 0,155 | 0 | Mature NK-a | PLAC8 |
| CLIC3 | 0 | 3,2098007 | 0,474 | 0,062 | 0 | Mature NK-a | CLIC3 |
| AUTS2 | 0 | 1,63434998 | 0,664 | 0,256 | 0 | Mature NK-a | AUTS2 |
| SORL1 | 0 | 2,44123833 | 0,54 | 0,132 | 0 | Mature NK-a | SORL1 |
| ABHD17A | 0 | 2,02903155 | 0,568 | 0,166 | 0 | Mature NK-a | ABHD17A |
| ZEB2 | 0 | 0,94749789 | 0,809 | 0,412 | 0 | Mature NK-a | ZEB2 |
| TLE1 | 0 | 2,7159176 | 0,471 | 0,092 | 0 | Mature NK-a | TLE1 |
| CMIP | 0 | 1,61618032 | 0,657 | 0,28 | 0 | Mature NK-a | CMIP |
| GZMM1 | 0 | 1,7128943 | 0,603 | 0,229 | 0 | Mature NK-a | GZMM |
| EFHD2 | 0 | 1,74576091 | 0,587 | 0,214 | 0 | Mature NK-a | EFHD2 |
| AKNA1 | 0 | 1,40764864 | 0,692 | 0,32 | 0 | Mature NK-a | AKNA |
| ISG201 | 0 | 1,3247996 | 0,753 | 0,386 | 0 | Mature NK-a | ISG20 |
| KLRB1 | 0 | 0,78525189 | 0,688 | 0,323 | 0 | Mature NK-a | KLRB1 |
| VAV3 | 0 | 1,5607109 | 0,651 | 0,289 | 0 | Mature NK-a | VAV3 |
| PRKCH | 0 | 1,41398578 | 0,785 | 0,424 | 0 | Mature NK-a | PRKCH |
| RASA2 | 0 | 1,46356853 | 0,758 | 0,397 | 0 | Mature NK-a | RASA2 |
| HAVCR2 | 0 | 2,83577858 | 0,425 | 0,068 | 0 | Mature NK-a | HAVCR2 |
| PITPNC1 | 0 | 1,56890797 | 0,848 | 0,493 | 0 | Mature NK-a | PITPNC1 |
| LYN | 0 | 1,56751369 | 0,509 | 0,156 | 0 | Mature NK-a | LYN |
| DSTN | 0 | 1,72520743 | 0,551 | 0,2 | 0 | Mature NK-a | DSTN |
| SYNE1 | 0 | 1,77164009 | 0,517 | 0,177 | 0 | Mature NK-a | SYNE1 |
| AREG | 0 | 0,77019973 | 0,66 | 0,326 | 0 | Mature NK-a | AREG |
| LITAF1 | 0 | 1,1168789 | 0,733 | 0,4 | 0 | Mature NK-a | LITAF |
| BNC2 | 0 | 3,0277953 | 0,384 | 0,052 | 0 | Mature NK-a | BNC2 |
| ATP8A11 | 0 | 1,65134048 | 0,529 | 0,198 | 0 | Mature NK-a | ATP8A1 |
| PTPN12 | 0 | 1,97275707 | 0,458 | 0,128 | 0 | Mature NK-a | PTPN12 |
| ARPC2 | 0 | 1,19583397 | 0,811 | 0,483 | 0 | Mature NK-a | ARPC2 |
| PIP4K2A | 0 | 1,2462137 | 0,698 | 0,374 | 0 | Mature NK-a | PIP4K2A |
| NCALD | 0 | 1,59393764 | 0,519 | 0,202 | 0 | Mature NK-a | NCALD |
| IFITM2 | 0 | 1,12585051 | 0,744 | 0,428 | 0 | Mature NK-a | IFITM2 |
| YES1 | 0 | 1,92755118 | 0,446 | 0,13 | 0 | Mature NK-a | YES1 |
| GZMA | 0 | 1,19036246 | 0,64 | 0,325 | 0 | Mature NK-a | GZMA |
| GNG2 | 0 | 0,95919211 | 0,774 | 0,461 | 0 | Mature NK-a | GNG2 |
| GZMH | 0 | 1,25547004 | 0,516 | 0,204 | 0 | Mature NK-a | GZMH |
| GK5 | 0 | 2,73773496 | 0,375 | 0,065 | 0 | Mature NK-a | GK5 |
| PDE4D | 0 | 1,15478204 | 0,62 | 0,31 | 0 | Mature NK-a | PDE4D |
| CNOT2 | 0 | 1,43048037 | 0,585 | 0,275 | 0 | Mature NK-a | CNOT2 |
| HSH2D | 0 | 2,9712046 | 0,357 | 0,051 | 0 | Mature NK-a | HSH2D |
| CEMIP21 | 0 | 0,92579232 | 0,871 | 0,566 | 0 | Mature NK-a | CEMIP2 |
| APBA2 | 0 | 2,17672381 | 0,398 | 0,097 | 0 | Mature NK-a | APBA2 |
| SSH2 | 0 | 1,28036062 | 0,626 | 0,327 | 0 | Mature NK-a | SSH2 |
| LAIR2 | 0 | 3,0463524 | 0,342 | 0,05 | 0 | Mature NK-a | LAIR2 |
| PYHIN1 | 0 | 1,61147383 | 0,455 | 0,164 | 0 | Mature NK-a | PYHIN1 |
| DIP2A | 0 | 1,80062359 | 0,434 | 0,143 | 0 | Mature NK-a | DIP2A |
| CMC11 | 0 | 1,54338147 | 0,441 | 0,151 | 0 | Mature NK-a | CMC1 |
| SYTL3 | 0 | 1,21769684 | 0,917 | 0,63 | 0 | Mature NK-a | SYTL3 |
| RUNX31 | 0 | 0,80168106 | 0,732 | 0,446 | 0 | Mature NK-a | RUNX3 |
| CHST12 | 0 | 2,42956955 | 0,359 | 0,073 | 0 | Mature NK-a | CHST12 |
| UTRN | 0 | 1,15963311 | 0,625 | 0,34 | 0 | Mature NK-a | UTRN |
| PPP2R5C1 | 0 | 0,80525357 | 0,865 | 0,581 | 0 | Mature NK-a | PPP2R5C |
| PDGFD | 0 | 2,36628216 | 0,361 | 0,077 | 0 | Mature NK-a | PDGFD |
| SH2D2A | 0 | 1,92920265 | 0,401 | 0,119 | 0 | Mature NK-a | SH2D2A |

| row.names | p_val | avg_log2FC | pct.1 | pct.2 | p_val_ | cluster | gene |
|---|---|---|---|---|---|---|---|
| GNLY1 | 0 | 1,3205158 | 0,86 | 0,26 | 0 | T/NK-a | GNLY |
| GZMH1 | 0 | 2,32212871 | 0,779 | 0,195 | 0 | T/NK-a | GZMH |
| GZMB1 | 0 | 1,36464519 | 0,822 | 0,242 | 0 | T/NK-a | GZMB |
| NKG71 | 0 | 1,52965614 | 0,95 | 0,381 | 0 | T/NK-a | NKG7 |
| FGFBP21 | 0 | 2,00103172 | 0,647 | 0,141 | 0 | T/NK-a | FGFBP2 |
| C1orf211 | 0 | 2,35622914 | 0,651 | 0,15 | 0 | T/NK-a | C1orf21 |
| PRF11 | 0 | 1,41306167 | 0,644 | 0,211 | 0 | T/NK-a | PRF1 |
| ZEB21 | 0 | 1,08505905 | 0,853 | 0,42 | 0 | T/NK-a | ZEB2 |
| CST72 | 0 | 1,15869768 | 0,848 | 0,436 | 0 | T/NK-a | CST7 |
| KLRD11 | 0 | 1,03477073 | 0,654 | 0,258 | 0 | T/NK-a | KLRD1 |
| CCL51 | 0 | 0,74009572 | 0,922 | 0,529 | 0 | T/NK-a | CCL5 |
| METRNL1 | 0 | 1,16555932 | 0,778 | 0,387 | 0 | T/NK-a | METRNL |
| CMIP1 | 0 | 1,6658213 | 0,679 | 0,289 | 0 | T/NK-a | CMIP |
| AOAH1 | 0 | 1,46133218 | 0,702 | 0,327 | 0 | T/NK-a | AOAH |
| TGFBR31 | 0 | 1,92700411 | 0,564 | 0,2 | 0 | T/NK-a | TGFBR3 |
| AKNA2 | 0 | 1,23691721 | 0,688 | 0,331 | 0 | T/NK-a | AKNA |
| ISG202 | 0 | 1,19308848 | 0,747 | 0,397 | 0 | T/NK-a | ISG20 |
| CD3G | 0 | 1,54048791 | 0,579 | 0,233 | 0 | T/NK-a | CD3G |
| CD8A1 | 0 | 1,90776776 | 0,478 | 0,138 | 0 | T/NK-a | CD8A |
| PPP2R5C2 | 0 | 1,12037731 | 0,917 | 0,585 | 0 | T/NK-a | PPP2R5C |
| PDE3B1 | 0 | 1,55584482 | 0,688 | 0,359 | 0 | T/NK-a | PDE3B |
| SAMD32 | 0 | 1,62287123 | 0,546 | 0,219 | 0 | T/NK-a | SAMD3 |
| P2RY83 | 0 | 1,24856056 | 0,647 | 0,323 | 0 | T/NK-a | P2RY8 |
| ATP8A12 | 0 | 1,54455664 | 0,527 | 0,207 | 0 | T/NK-a | ATP8A1 |
| LINC018711 | 0 | 1,40896953 | 0,535 | 0,221 | 0 | T/NK-a | LINC01871 |
| PRKCH1 | 0 | 1,05371433 | 0,749 | 0,436 | 0 | T/NK-a | PRKCH |
| SYNE11 | 0 | 1,50352868 | 0,495 | 0,188 | 0 | T/NK-a | SYNE1 |
| SLA21 | 0 | 1,7859651 | 0,442 | 0,135 | 0 | T/NK-a | SLA2 |
| DSTN1 | 0 | 1,49655742 | 0,517 | 0,212 | 0 | T/NK-a | DSTN |
| CTSW1 | 0 | 0,96034372 | 0,555 | 0,253 | 0 | T/NK-a | CTSW |
| AUTS21 | 0 | 1,16338835 | 0,574 | 0,273 | 0 | T/NK-a | AUTS2 |
| PPP1R16B2 | 0 | 1,18821526 | 0,611 | 0,315 | 0 | T/NK-a | PPP1R16B |
| AC243829.21 | 0 | 3,02127515 | 0,336 | 0,046 | 0 | T/NK-a | AC243829.2 |
| CD3D2 | 0 | 1,18612457 | 0,582 | 0,292 | 0 | T/NK-a | CD3D |
| RASA21 | 0 | 1,01454899 | 0,699 | 0,411 | 0 | T/NK-a | RASA2 |
| SYTL31 | 0 | 0,96734912 | 0,921 | 0,638 | 0 | T/NK-a | SYTL3 |
| RALGAPA12 | 0 | 0,96065708 | 0,773 | 0,497 | 0 | T/NK-a | RALGAPA1 |
| NFATC22 | 0 | 1,43000653 | 0,455 | 0,19 | 0 | T/NK-a | NFATC2 |
| RABGAP1L | 0 | 1,05976369 | 0,696 | 0,432 | 0 | T/NK-a | RABGAP1L |
| SRPK21 | 0 | 1,36842568 | 0,482 | 0,22 | 0 | T/NK-a | SRPK2 |
| SPON21 | 0 | 0,92365793 | 0,423 | 0,167 | 0 | T/NK-a | SPON2 |
| ABHD17A1 | 0 | 1,20667917 | 0,437 | 0,186 | 0 | T/NK-a | ABHD17A |
| TFDP21 | 0 | 1,35048776 | 0,434 | 0,185 | 0 | T/NK-a | TFDP2 |
| PTPRJ1 | 0 | 1,29081099 | 0,467 | 0,219 | 0 | T/NK-a | PTPRJ |
| RASGRP11 | 0 | 1,34975674 | 0,437 | 0,197 | 0 | T/NK-a | RASGRP1 |
| DIP2A1 | 0 | 1,44030758 | 0,383 | 0,154 | 0 | T/NK-a | DIP2A |
| SGCD1 | 0 | 3,11885277 | 0,258 | 0,033 | 0 | T/NK-a | SGCD |
| TRGC21 | 0 | 1,85938632 | 0,316 | 0,095 | 0 | T/NK-a | TRGC2 |
| MYO1F1 | 0 | 1,5534548 | 0,351 | 0,134 | 0 | T/NK-a | MYO1F |
| TERF11 | 0 | 1,81208794 | 0,293 | 0,09 | 0 | T/NK-a | TERF1 |
| GGA21 | 0 | 1,62114435 | 0,297 | 0,104 | 0 | T/NK-a | GGA2 |
| GTF3C11 | 0 | 1,7519604 | 0,275 | 0,084 | 0 | T/NK-a | GTF3C1 |
| CD8B1 | 0 | 1,79540834 | 0,247 | 0,076 | 0 | T/NK-a | CD8B |
| CADM11 | 0 | 1,92687348 | 0,237 | 0,067 | 0 | T/NK-a | CADM1 |
| S1PR51 | 0 | 1,99754002 | 0,213 | 0,052 | 0 | T/NK-a | S1PR5 |
| TMCC31 | 0 | 2,02393285 | 0,213 | 0,057 | 0 | T/NK-a | TMCC3 |
| SETBP11 | 0 | 2,18243385 | 0,176 | 0,042 | 0 | T/NK-a | SETBP1 |
| SLC1A41 | 0 | 2,3050647 | 0,161 | 0,036 | 0 | T/NK-a | SLC1A4 |
| HLA-C1 | 0 | 0,69364891 | 0,954 | 0,836 | 0 | T/NK-a | HLA-C |
| LILRB11 | 0 | 2,51243423 | 0,123 | 0,022 | 0 | T/NK-a | LILRB1 |
| ATL11 | 0 | 3,18043595 | 0,087 | 0,01 | 0 | T/NK-a | ATL1 |
| BPTF1 | 1E-307 | 1,23421183 | 0,511 | 0,26 | ##### | T/NK-a | BPTF |
| ITGAL1 | 1E-306 | 1,51218298 | 0,333 | 0,126 | ##### | T/NK-a | ITGAL |
| HLA-E1 | 2E-293 | 0,64236858 | 0,885 | 0,686 | ##### | T/NK-a | HLA-E |
| CHST121 | 4E-292 | 1,62783315 | 0,262 | 0,087 | ##### | T/NK-a | CHST12 |
| TMEM1811 | 2E-290 | 1,4378477 | 0,357 | 0,146 | ##### | T/NK-a | TMEM181 |
| C12orf752 | 2E-290 | 1,27661398 | 0,363 | 0,147 | ##### | T/NK-a | C12orf75 |
| IQGAP22 | 1E-287 | 0,79870084 | 0,822 | 0,6 | ##### | T/NK-a | IQGAP2 |
| FAM53B1 | 1E-287 | 1,88019513 | 0,225 | 0,069 | ##### | T/NK-a | FAM53B |
| PRKCB1 | 2E-282 | 1,23341097 | 0,463 | 0,225 | ##### | T/NK-a | PRKCB |
| PTPRC1 | 1E-280 | 0,57425753 | 0,965 | 0,811 | ##### | T/NK-a | PTPRC |
| PLEKHA21 | 6E-279 | 1,11929971 | 0,509 | 0,263 | ##### | T/NK-a | PLEKHA2 |
| ADGRG11 | 3E-276 | 1,78913898 | 0,198 | 0,056 | ##### | T/NK-a | ADGRG1 |
| PITPNC11 | 5E-275 | 0,91302053 | 0,755 | 0,509 | ##### | T/NK-a | PITPNC1 |

| row.names | p_val | avg_log2FC | pct.1 | pct.2 | p_val_a | cluster | gene |
|---|---|---|---|---|---|---|---|
| CCR61 | 0 | 3,29446101 | 0,63 | 0,074 | 0 | Early activate | CCR6 |
| RBMS11 | 0 | 2,22908231 | 0,754 | 0,283 | 0 | Early activate | RBMS1 |
| IL7R2 | 0 | 1,09634671 | 0,949 | 0,49 | 0 | Early activate | IL7R |
| LTB1 | 0 | 2,35237841 | 0,625 | 0,177 | 0 | Early activate | LTB |
| TOB1 | 0 | 1,56105116 | 0,622 | 0,25 | 0 | Early activate | TOB1 |
| PTPN131 | 0 | 4,80626944 | 0,387 | 0,021 | 0 | Early activate | PTPN13 |
| RORA1 | 0 | 1,30136973 | 0,788 | 0,424 | 0 | Early activate | RORA |
| TTC39C1 | 0 | 2,10036654 | 0,487 | 0,138 | 0 | Early activate | TTC39C |
| MAP3K4 | 0 | 2,81057101 | 0,39 | 0,07 | 0 | Early activate | MAP3K4 |
| PDE4D1 | 0 | 1,12395575 | 0,629 | 0,319 | 0 | Early activate | PDE4D |
| KLRB11 | 0 | 1,1351706 | 0,641 | 0,338 | 0 | Early activate | KLRB1 |
| CAMK42 | 0 | 1,43885879 | 0,513 | 0,214 | 0 | Early activate | CAMK4 |
| MGAT4A1 | 0 | 1,70257364 | 0,469 | 0,175 | 0 | Early activate | MGAT4A |
| RUNX21 | 0 | 2,04322799 | 0,377 | 0,097 | 0 | Early activate | RUNX2 |
| EEF1G | 0 | 0,97787799 | 0,756 | 0,507 | 0 | Early activate | EEF1G |
| RPLP0 | 0 | 1,26237545 | 0,909 | 0,678 | 0 | Early activate | RPLP0 |
| AQP31 | 0 | 2,33441609 | 0,25 | 0,052 | 0 | Early activate | AQP3 |
| EEF1B2 | 0 | 0,90176633 | 0,841 | 0,646 | 0 | Early activate | EEF1B2 |
| AL136456.1 | 0 | 1,05036096 | 0,256 | 0,064 | 0 | Early activate | AL136456.1 |
| RPSA1 | 0 | 1,00915032 | 0,916 | 0,758 | 0 | Early activate | RPSA |
| RPL22 | 0 | 0,72339832 | 0,903 | 0,774 | 0 | Early activate | RPL22 |
| MCAM1 | 0 | 2,98668894 | 0,145 | 0,019 | 0 | Early activate | MCAM |
| RPS51 | 0 | 0,72079607 | 0,893 | 0,768 | 0 | Early activate | RPS5 |
| AL121935.11 | 0 | 2,9594188 | 0,136 | 0,018 | 0 | Early activate | AL121935.1 |
| KIF5C | 0 | 2,52289957 | 0,143 | 0,025 | 0 | Early activate | KIF5C |
| RPL36A1 | 0 | 0,85830568 | 0,928 | 0,81 | 0 | Early activate | RPL36A |
| RPL10A1 | 0 | 0,79158903 | 0,921 | 0,816 | 0 | Early activate | RPL10A |
| AL589693.11 | 0 | 3,00644433 | 0,106 | 0,012 | 0 | Early activate | AL589693.1 |
| RPL351 | 0 | 0,69992998 | 0,934 | 0,849 | 0 | Early activate | RPL35 |
| RPL27A | 0 | 0,6744559 | 0,906 | 0,824 | 0 | Early activate | RPL27A |
| RPL381 | 0 | 0,67670926 | 0,931 | 0,849 | 0 | Early activate | RPL38 |
| RPS161 | 0 | 0,63559949 | 0,943 | 0,87 | 0 | Early activate | RPS16 |
| RPL5 | 0 | 0,70672949 | 0,915 | 0,845 | 0 | Early activate | RPL5 |
| RPL241 | 0 | 0,6077004 | 0,929 | 0,86 | 0 | Early activate | RPL24 |
| RPL361 | 0 | 0,71947671 | 0,943 | 0,881 | 0 | Early activate | RPL36 |
| RPL29 | 0 | 0,69397069 | 0,938 | 0,878 | 0 | Early activate | RPL29 |
| RPS13 | 0 | 0,75262413 | 0,935 | 0,877 | 0 | Early activate | RPS13 |
| RPL141 | 0 | 0,72490906 | 0,944 | 0,889 | 0 | Early activate | RPL14 |
| RPS251 | 0 | 0,65642356 | 0,943 | 0,89 | 0 | Early activate | RPS25 |
| RPL18 | 0 | 0,79301389 | 0,942 | 0,895 | 0 | Early activate | RPL18 |
| RPL6 | 0 | 0,66726252 | 0,931 | 0,885 | 0 | Early activate | RPL6 |
| RPL91 | 0 | 0,76040155 | 0,938 | 0,892 | 0 | Early activate | RPL9 |
| RPL23A1 | 0 | 0,59596205 | 0,938 | 0,893 | 0 | Early activate | RPL23A |
| RPL35A1 | 0 | 0,64699279 | 0,944 | 0,9 | 0 | Early activate | RPL35A |
| RPL3 | 0 | 0,76355077 | 0,944 | 0,901 | 0 | Early activate | RPL3 |
| RPS6 | 0 | 0,79335964 | 0,949 | 0,91 | 0 | Early activate | RPS6 |
| RPS211 | 0 | 0,64692242 | 0,955 | 0,92 | 0 | Early activate | RPS21 |
| RPL121 | 0 | 0,75762707 | 0,951 | 0,917 | 0 | Early activate | RPL12 |
| RPL171 | 0 | 0,74008415 | 0,942 | 0,909 | 0 | Early activate | RPL17 |
| RPS31 | 0 | 0,59544802 | 0,946 | 0,914 | 0 | Early activate | RPS3 |
| RPL18A1 | 0 | 0,73104968 | 0,95 | 0,918 | 0 | Early activate | RPL18A |
| RPLP21 | 0 | 0,63674247 | 0,951 | 0,925 | 0 | Early activate | RPLP2 |
| RPS8 | 0 | 0,74943147 | 0,948 | 0,923 | 0 | Early activate | RPS8 |
| RPS4X1 | 0 | 0,73369757 | 0,951 | 0,928 | 0 | Early activate | RPS4X |
| RPS191 | 0 | 0,79125061 | 0,956 | 0,937 | 0 | Early activate | RPS19 |
| RPS2 | 0 | 0,76166013 | 0,949 | 0,932 | 0 | Early activate | RPS2 |
| RPS23 | 0 | 0,58359289 | 0,952 | 0,937 | 0 | Early activate | RPS23 |
| RPL191 | 0 | 0,68174483 | 0,948 | 0,934 | 0 | Early activate | RPL19 |
| RPS151 | 0 | 0,56614226 | 0,943 | 0,93 | 0 | Early activate | RPS15 |
| RPL411 | 0 | 0,50662942 | 0,962 | 0,974 | 0 | Early activate | RPL41 |
| RPL111 | 0 | 0,70381294 | 0,956 | 0,946 | 0 | Early activate | RPL11 |
| RPL101 | 0 | 0,67275293 | 0,96 | 0,97 | 0 | Early activate | RPL10 |
| RPL21 | 0 | 0,57650581 | 0,946 | 0,936 | 0 | Early activate | RPL21 |
| RPS3A | 0 | 0,6204383 | 0,95 | 0,941 | 0 | Early activate | RPS3A |
| RPL321 | 0 | 0,78056574 | 0,956 | 0,948 | 0 | Early activate | RPL32 |
| TPT1 | 0 | 0,79070932 | 0,958 | 0,966 | 0 | Early activate | TPT1 |
| EEF1A1 | 0 | 0,72206507 | 0,969 | 0,975 | 0 | Early activate | EEF1A1 |
| RPS15A1 | 0 | 0,53948458 | 0,953 | 0,947 | 0 | Early activate | RPS15A |
| RPS281 | 0 | 0,57744044 | 0,956 | 0,95 | 0 | Early activate | RPS28 |
| RPS141 | 0 | 0,62975062 | 0,952 | 0,947 | 0 | Early activate | RPS14 |
| RPLP1 | 0 | 0,95697052 | 0,96 | 0,965 | 0 | Early activate | RPLP1 |
| RPL341 | 0 | 0,76947016 | 0,959 | 0,955 | 0 | Early activate | RPL34 |
| RPS181 | 0 | 0,84735916 | 0,955 | 0,951 | 0 | Early activate | RPS18 |
| RPL301 | 0 | 0,63775412 | 0,954 | 0,95 | 0 | Early activate | RPL30 |

| row.names | p_val | avg_log2FC | pct.1 | pct.2 | p_val_ad | cluster | gene |
|---|---|---|---|---|---|---|---|
| LYZ | 0 | 5,10374149 | 0,979 | 0,067 | 0 | ATMs | LYZ |
| CST3 | 0 | 2,35486004 | 0,978 | 0,108 | 0 | ATMs | CST3 |
| C5AR1 | 0 | 4,64830293 | 0,897 | 0,042 | 0 | ATMs | C5AR1 |
| KYNU | 0 | 4,3766247 | 0,87 | 0,06 | 0 | ATMs | KYNU |
| IRAK3 | 0 | 3,6445595 | 0,898 | 0,088 | 0 | ATMs | IRAK3 |
| AIF1 | 0 | 3,64392052 | 0,868 | 0,07 | 0 | ATMs | AIF1 |
| PLAUR | 0 | 3,89820718 | 0,895 | 0,1 | 0 | ATMs | PLAUR |
| CTSS | 0 | 3,71235877 | 0,952 | 0,158 | 0 | ATMs | CTSS |
| HLA-DRA | 0 | 2,45135978 | 0,98 | 0,188 | 0 | ATMs | HLA-DRA |
| FCN1 | 0 | 5,58084119 | 0,808 | 0,024 | 0 | ATMs | FCN1 |
| PSAP | 0 | 3,31984443 | 0,917 | 0,142 | 0 | ATMs | PSAP |
| FCER1G1 | 0 | 2,65469385 | 0,922 | 0,171 | 0 | ATMs | FCER1G |
| RBM47 | 0 | 4,87757664 | 0,799 | 0,048 | 0 | ATMs | RBM47 |
| CTSB | 0 | 3,83194324 | 0,823 | 0,073 | 0 | ATMs | CTSB |
| VCAN | 0 | 7,23092734 | 0,757 | 0,012 | 0 | ATMs | VCAN |
| IER3 | 0 | 4,5909198 | 0,806 | 0,063 | 0 | ATMs | IER3 |
| ACSL1 | 0 | 4,27188415 | 0,806 | 0,064 | 0 | ATMs | ACSL1 |
| PID1 | 0 | 4,63042736 | 0,772 | 0,04 | 0 | ATMs | PID1 |
| CXCL8 | 0 | 4,44991767 | 0,797 | 0,065 | 0 | ATMs | CXCL8 |
| RNF144B | 0 | 3,41250252 | 0,801 | 0,073 | 0 | ATMs | RNF144B |
| MCTP1 | 0 | 4,10702928 | 0,788 | 0,06 | 0 | ATMs | MCTP1 |
| DOCK4 | 0 | 5,34166607 | 0,766 | 0,042 | 0 | ATMs | DOCK4 |
| TYROBP1 | 0 | 2,5228391 | 0,974 | 0,25 | 0 | ATMs | TYROBP |
| RAB31 | 0 | 4,09118415 | 0,767 | 0,056 | 0 | ATMs | RAB31 |
| SOD2 | 0 | 4,26261888 | 0,886 | 0,183 | 0 | ATMs | SOD2 |
| CXCL2 | 0 | 4,84610613 | 0,745 | 0,044 | 0 | ATMs | CXCL2 |
| FGD4 | 0 | 4,64828559 | 0,744 | 0,043 | 0 | ATMs | FGD4 |
| BCL2A1 | 0 | 4,00915761 | 0,773 | 0,075 | 0 | ATMs | BCL2A1 |
| CD163 | 0 | 4,33356522 | 0,722 | 0,025 | 0 | ATMs | CD163 |
| PLXDC2 | 0 | 4,08841981 | 0,752 | 0,061 | 0 | ATMs | PLXDC2 |
| NAMPT | 0 | 3,35603691 | 0,971 | 0,284 | 0 | ATMs | NAMPT |
| TLR2 | 0 | 4,23207003 | 0,726 | 0,041 | 0 | ATMs | TLR2 |
| MS4A6A | 0 | 3,74202154 | 0,736 | 0,053 | 0 | ATMs | MS4A6A |
| NLRP3 | 0 | 3,76790618 | 0,764 | 0,085 | 0 | ATMs | NLRP3 |
| EREG | 0 | 5,81288525 | 0,706 | 0,032 | 0 | ATMs | EREG |
| NPC2 | 0 | 3,15256743 | 0,76 | 0,098 | 0 | ATMs | NPC2 |
| ATP13A3 | 0 | 3,86239279 | 0,769 | 0,108 | 0 | ATMs | ATP13A3 |
| HLA-DRB1 | 0 | 2,1141475 | 0,929 | 0,268 | 0 | ATMs | HLA-DRB1 |
| HLA-DPA1 | 0 | 2,06823579 | 0,858 | 0,197 | 0 | ATMs | HLA-DPA1 |
| FAM49A | 0 | 3,18322729 | 0,759 | 0,101 | 0 | ATMs | FAM49A |
| IL1B | 0 | 4,91813828 | 0,702 | 0,044 | 0 | ATMs | IL1B |
| LRMDA | 0 | 4,09437966 | 0,705 | 0,049 | 0 | ATMs | LRMDA |
| DENND5A | 0 | 4,63490542 | 0,704 | 0,05 | 0 | ATMs | DENND5A |
| IFI30 | 0 | 3,50211622 | 0,717 | 0,067 | 0 | ATMs | IFI30 |
| THBS1 | 0 | 6,08084633 | 0,68 | 0,031 | 0 | ATMs | THBS1 |
| ETS2 | 0 | 3,4330601 | 0,721 | 0,076 | 0 | ATMs | ETS2 |
| FCGR2A | 0 | 4,0091565 | 0,683 | 0,044 | 0 | ATMs | FCGR2A |
| CD83 | 0 | 2,5747256 | 0,754 | 0,119 | 0 | ATMs | CD83 |
| FNDC3B | 0 | 3,54278005 | 0,806 | 0,175 | 0 | ATMs | FNDC3B |
| ANXA5 | 0 | 2,80190722 | 0,78 | 0,151 | 0 | ATMs | ANXA5 |
| CD74 | 0 | 2,05584236 | 0,97 | 0,341 | 0 | ATMs | CD74 |
| HLA-DPB1 | 0 | 1,91910581 | 0,846 | 0,22 | 0 | ATMs | HLA-DPB1 |
| SESTD1 | 0 | 4,50375456 | 0,676 | 0,052 | 0 | ATMs | SESTD1 |
| LST1 | 0 | 2,82590127 | 0,689 | 0,066 | 0 | ATMs | LST1 |
| LCP2 | 0 | 3,42781766 | 0,715 | 0,094 | 0 | ATMs | LCP2 |
| TET2 | 0 | 2,99465926 | 0,786 | 0,169 | 0 | ATMs | TET2 |
| EPB41L3 | 0 | 4,33537651 | 0,655 | 0,041 | 0 | ATMs | EPB41L3 |
| HLA-DQB1 | 0 | 1,76296709 | 0,781 | 0,175 | 0 | ATMs | HLA-DQB1 |
| MGAT1 | 0 | 3,21864967 | 0,699 | 0,094 | 0 | ATMs | MGAT1 |
| OLR1 | 0 | 5,99109544 | 0,62 | 0,016 | 0 | ATMs | OLR1 |
| NUMB | 0 | 3,76837448 | 0,7 | 0,097 | 0 | ATMs | NUMB |
| APLP2 | 0 | 2,84566933 | 0,729 | 0,13 | 0 | ATMs | APLP2 |
| SAT1 | 0 | 2,74851975 | 0,982 | 0,384 | 0 | ATMs | SAT1 |
| MXD1 | 0 | 3,58414573 | 0,685 | 0,091 | 0 | ATMs | MXD1 |
| AC020656.1 | 0 | 5,04605957 | 0,616 | 0,024 | 0 | ATMs | AC020656.1 |
| QKI | 0 | 2,46804241 | 0,802 | 0,212 | 0 | ATMs | QKI |
| ICAM1 | 0 | 4,16691459 | 0,635 | 0,047 | 0 | ATMs | ICAM1 |
| CXCL3 | 0 | 5,04030062 | 0,614 | 0,03 | 0 | ATMs | CXCL3 |
| HLA-DRB5 | 0 | 2,38107258 | 0,693 | 0,113 | 0 | ATMs | HLA-DRB5 |
| SIPA1L1 | 0 | 2,57106642 | 0,944 | 0,365 | 0 | ATMs | SIPA1L1 |
| FGL2 | 0 | 2,79949416 | 0,698 | 0,124 | 0 | ATMs | FGL2 |
| GAB2 | 0 | 3,14475161 | 0,681 | 0,11 | 0 | ATMs | GAB2 |
| LUCAT1 | 0 | 5,14253319 | 0,598 | 0,027 | 0 | ATMs | LUCAT1 |
| CYBB | 0 | 4,48427886 | 0,599 | 0,028 | 0 | ATMs | CYBB |

| row.names | p_val | avg_log2FC | pct.1 | pct.2 | p_val_adj | cluster | gene |
|---|---|---|---|---|---|---|---|
| GNLY2 | 0 | 2,34873749 | 0,956 | 0,261 | 0 | Mature NK-b | GNLY |
| KLRD12 | 0 | 2,49280814 | 0,936 | 0,247 | 0 | Mature NK-b | KLRD1 |
| GZMB2 | 0 | 1,80801434 | 0,897 | 0,245 | 0 | Mature NK-b | GZMB |
| KLRF11 | 0 | 3,18388019 | 0,733 | 0,101 | 0 | Mature NK-b | KLRF1 |
| SPON22 | 0 | 2,66118432 | 0,753 | 0,152 | 0 | Mature NK-b | SPON2 |
| NKG72 | 0 | 1,97479449 | 0,985 | 0,385 | 0 | Mature NK-b | NKG7 |
| FGFBP22 | 0 | 2,31275924 | 0,715 | 0,143 | 0 | Mature NK-b | FGFBP2 |
| CCL41 | 0 | 2,29478404 | 0,915 | 0,368 | 0 | Mature NK-b | CCL4 |
| PRF12 | 0 | 1,92688582 | 0,736 | 0,211 | 0 | Mature NK-b | PRF1 |
| FCGR3A2 | 0 | 2,60671675 | 0,641 | 0,117 | 0 | Mature NK-b | FCGR3A |
| CD2472 | 0 | 1,77426546 | 0,921 | 0,416 | 0 | Mature NK-b | CD247 |
| TYROBP2 | 0 | 1,27526886 | 0,753 | 0,265 | 0 | Mature NK-b | TYROBP |
| CTSW2 | 0 | 1,75616811 | 0,724 | 0,247 | 0 | Mature NK-b | CTSW |
| CST73 | 0 | 1,30585346 | 0,876 | 0,439 | 0 | Mature NK-b | CST7 |
| MCTP22 | 0 | 2,01574139 | 0,585 | 0,156 | 0 | Mature NK-b | MCTP2 |
| GZMA1 | 0 | 1,48097561 | 0,736 | 0,331 | 0 | Mature NK-b | GZMA |
| AOAH2 | 0 | 1,07704645 | 0,724 | 0,33 | 0 | Mature NK-b | AOAH |
| MYOM21 | 0 | 2,79808201 | 0,466 | 0,073 | 0 | Mature NK-b | MYOM2 |
| ZEB23 | 0 | 1,22572576 | 0,814 | 0,427 | 0 | Mature NK-b | ZEB2 |
| CD71 | 0 | 1,40652754 | 0,667 | 0,281 | 0 | Mature NK-b | CD7 |
| TXK1 | 0 | 2,74713644 | 0,462 | 0,082 | 0 | Mature NK-b | TXK |
| PIP4K2A2 | 0 | 1,5884374 | 0,761 | 0,383 | 0 | Mature NK-b | PIP4K2A |
| CMC12 | 0 | 1,81638375 | 0,532 | 0,157 | 0 | Mature NK-b | CMC1 |
| YES12 | 0 | 2,31546156 | 0,509 | 0,139 | 0 | Mature NK-b | YES1 |
| CCL31 | 0 | 2,7038063 | 0,452 | 0,085 | 0 | Mature NK-b | CCL3 |
| GZMH2 | 0 | 1,39534892 | 0,577 | 0,212 | 0 | Mature NK-b | GZMH |
| PLAC82 | 0 | 1,78296308 | 0,535 | 0,173 | 0 | Mature NK-b | PLAC8 |
| KLRK13 | 0 | 1,71546659 | 0,54 | 0,182 | 0 | Mature NK-b | KLRK1 |
| PYHIN12 | 0 | 1,83185151 | 0,527 | 0,171 | 0 | Mature NK-b | PYHIN1 |
| EFHD22 | 0 | 1,59277393 | 0,58 | 0,228 | 0 | Mature NK-b | EFHD2 |
| PTPN123 | 0 | 1,97469622 | 0,473 | 0,139 | 0 | Mature NK-b | PTPN12 |
| PRKCH2 | 0 | 1,25209033 | 0,769 | 0,438 | 0 | Mature NK-b | PRKCH |
| CARD112 | 0 | 1,61808354 | 0,521 | 0,191 | 0 | Mature NK-b | CARD11 |
| HOPX1 | 0 | 1,97858778 | 0,459 | 0,141 | 0 | Mature NK-b | HOPX |
| CLIC31 | 0 | 2,28925089 | 0,398 | 0,082 | 0 | Mature NK-b | CLIC3 |
| GZMM3 | 0 | 1,35859185 | 0,557 | 0,246 | 0 | Mature NK-b | GZMM |
| SYNE12 | 0 | 1,58325497 | 0,501 | 0,191 | 0 | Mature NK-b | SYNE1 |
| CCL4L21 | 0 | 2,43997538 | 0,425 | 0,115 | 0 | Mature NK-b | CCL4L2 |
| ABHD17A2 | 0 | 1,61631716 | 0,495 | 0,185 | 0 | Mature NK-b | ABHD17A |
| BNC21 | 0 | 2,55321772 | 0,371 | 0,066 | 0 | Mature NK-b | BNC2 |
| CCL52 | 0 | 0,94677575 | 0,841 | 0,538 | 0 | Mature NK-b | CCL5 |
| TFDP22 | 0 | 1,5189069 | 0,485 | 0,185 | 0 | Mature NK-b | TFDP2 |
| C1orf212 | 0 | 1,30125982 | 0,461 | 0,166 | 0 | Mature NK-b | C1orf21 |
| CEP782 | 0 | 2,43316144 | 0,371 | 0,076 | 0 | Mature NK-b | CEP78 |
| PDGFD2 | 0 | 2,22371246 | 0,374 | 0,087 | 0 | Mature NK-b | PDGFD |
| KLRC33 | 0 | 2,06709453 | 0,371 | 0,089 | 0 | Mature NK-b | KLRC3 |
| LYN2 | 0 | 1,2490314 | 0,452 | 0,173 | 0 | Mature NK-b | LYN |
| GNPTAB2 | 0 | 1,90956404 | 0,398 | 0,119 | 0 | Mature NK-b | GNPTAB |
| ID2 | 0 | 1,23836582 | 0,695 | 0,419 | 0 | Mature NK-b | ID2 |
| ATM1 | 0 | 1,77181371 | 0,407 | 0,143 | 0 | Mature NK-b | ATM |
| TRDC1 | 0 | 2,78020333 | 0,307 | 0,045 | 0 | Mature NK-b | TRDC |
| BIN22 | 0 | 1,570204 | 0,419 | 0,16 | 0 | Mature NK-b | BIN2 |
| PFN11 | 0 | 1,22835228 | 0,803 | 0,546 | 0 | Mature NK-b | PFN1 |
| GSAP2 | 0 | 2,55104671 | 0,314 | 0,062 | 0 | Mature NK-b | GSAP |
| CYBA1 | 0 | 0,93450957 | 0,851 | 0,607 | 0 | Mature NK-b | CYBA |
| HSH2D1 | 0 | 2,283801 | 0,305 | 0,065 | 0 | Mature NK-b | HSH2D |
| ARRDC31 | 0 | 2,05859319 | 0,349 | 0,11 | 0 | Mature NK-b | ARRDC3 |
| APBA22 | 0 | 1,71650479 | 0,35 | 0,111 | 0 | Mature NK-b | APBA2 |
| KLRC23 | 0 | 2,48585841 | 0,282 | 0,055 | 0 | Mature NK-b | KLRC2 |
| ADGRG12 | 0 | 2,37541515 | 0,271 | 0,054 | 0 | Mature NK-b | ADGRG1 |
| GK52 | 0 | 1,85846064 | 0,295 | 0,081 | 0 | Mature NK-b | GK5 |
| HAVCR21 | 0 | 1,85427126 | 0,301 | 0,088 | 0 | Mature NK-b | HAVCR2 |
| LINC002991 | 0 | 2,34006538 | 0,265 | 0,055 | 0 | Mature NK-b | LINC00299 |
| IGFBP71 | 0 | 1,96722545 | 0,252 | 0,045 | 0 | Mature NK-b | IGFBP7 |
| AC016074.2 | 0 | 2,72638592 | 0,253 | 0,046 | 0 | Mature NK-b | AC016074.2 |
| NCR11 | 0 | 2,72627918 | 0,246 | 0,039 | 0 | Mature NK-b | NCR1 |
| LINC00861 | 0 | 2,23434485 | 0,276 | 0,07 | 0 | Mature NK-b | LINC00861 |
| HLA-E2 | 0 | 0,79195765 | 0,889 | 0,688 | 0 | Mature NK-b | HLA-E |
| C5orf562 | 0 | 1,77176886 | 0,287 | 0,086 | 0 | Mature NK-b | C5orf56 |
| LAIR22 | 0 | 2,08056302 | 0,265 | 0,065 | 0 | Mature NK-b | LAIR2 |
| TTC382 | 0 | 2,46033792 | 0,234 | 0,042 | 0 | Mature NK-b | TTC38 |
| SH2D1B1 | 0 | 2,78665474 | 0,221 | 0,034 | 0 | Mature NK-b | SH2D1B |
| NCAM12 | 0 | 2,26362151 | 0,228 | 0,048 | 0 | Mature NK-b | NCAM1 |
| AKR1C31 | 0 | 2,64477409 | 0,208 | 0,033 | 0 | Mature NK-b | AKR1C3 |

| row.names | p_val | avg_log2FC | pct.1 | pct.2 | p_val_ac | cluster | gene |
|---|---|---|---|---|---|---|---|
| CST31 | 0 | 4,8221062 | 0,991 | 0,118 | 0 | Intermediate | CST3 |
| HLA-DQA11 | 0 | 4,89191092 | 0,985 | 0,117 | 0 | Intermediate | HLA-DQA1 |
| LYZ1 | 0 | 2,93535721 | 0,941 | 0,08 | 0 | Intermediate | LYZ |
| HLA-DMA1 | 0 | 4,0603108 | 0,944 | 0,09 | 0 | Intermediate | HLA-DMA |
| CLEC10A1 | 0 | 5,54259961 | 0,857 | 0,028 | 0 | Intermediate | CLEC10A |
| FCER1A | 0 | 8,04916955 | 0,829 | 0,007 | 0 | Intermediate | FCER1A |
| IL1R21 | 0 | 5,58340792 | 0,857 | 0,038 | 0 | Intermediate | IL1R2 |
| HLA-DQB11 | 0 | 4,45845137 | 0,983 | 0,173 | 0 | Intermediate | HLA-DQB1 |
| HLA-DRA1 | 0 | 4,5448234 | 0,995 | 0,197 | 0 | Intermediate | HLA-DRA |
| HLA-DPA11 | 0 | 4,39759502 | 0,99 | 0,199 | 0 | Intermediate | HLA-DPA1 |
| HLA-DPB11 | 0 | 4,44466807 | 0,991 | 0,221 | 0 | Intermediate | HLA-DPB1 |
| CSF2RA1 | 0 | 4,51536085 | 0,803 | 0,049 | 0 | Intermediate | CSF2RA |
| CD1C | 0 | 7,8261847 | 0,757 | 0,006 | 0 | Intermediate | CD1C |
| FCER1G3 | 0 | 2,5046192 | 0,922 | 0,18 | 0 | Intermediate | FCER1G |
| PLAUR1 | 0 | 2,87824026 | 0,845 | 0,112 | 0 | Intermediate | PLAUR |
| CEBPD1 | 0 | 3,16217285 | 0,907 | 0,188 | 0 | Intermediate | CEBPD |
| HLA-DRB11 | 0 | 4,23721088 | 0,992 | 0,273 | 0 | Intermediate | HLA-DRB1 |
| RNF144B1 | 0 | 2,70270644 | 0,791 | 0,082 | 0 | Intermediate | RNF144B |
| SGK11 | 0 | 3,87371361 | 0,803 | 0,1 | 0 | Intermediate | SGK1 |
| IFI301 | 0 | 3,27184495 | 0,771 | 0,072 | 0 | Intermediate | IFI30 |
| CXCL161 | 0 | 4,21611092 | 0,741 | 0,048 | 0 | Intermediate | CXCL16 |
| KLF41 | 0 | 3,40155463 | 0,765 | 0,077 | 0 | Intermediate | KLF4 |
| TYROBP3 | 0 | 1,8728991 | 0,945 | 0,26 | 0 | Intermediate | TYROBP |
| CXCL81 | 0 | 3,07539934 | 0,757 | 0,075 | 0 | Intermediate | CXCL8 |
| CD831 | 0 | 3,02855199 | 0,805 | 0,124 | 0 | Intermediate | CD83 |
| CPVL1 | 0 | 4,02647755 | 0,716 | 0,04 | 0 | Intermediate | CPVL |
| CTSZ1 | 0 | 3,49469671 | 0,749 | 0,08 | 0 | Intermediate | CTSZ |
| MS4A6A1 | 0 | 3,67180059 | 0,729 | 0,061 | 0 | Intermediate | MS4A6A |
| C1orf1621 | 0 | 3,54368739 | 0,741 | 0,078 | 0 | Intermediate | C1orf162 |
| HLA-DRB51 | 0 | 4,28607666 | 0,779 | 0,116 | 0 | Intermediate | HLA-DRB5 |
| AIF11 | 0 | 2,55887675 | 0,744 | 0,085 | 0 | Intermediate | AIF1 |
| CD741 | 0 | 3,73137523 | 0,996 | 0,347 | 0 | Intermediate | CD74 |
| NLRP31 | 0 | 3,03936008 | 0,739 | 0,094 | 0 | Intermediate | NLRP3 |
| INSIG11 | 0 | 3,5640498 | 0,774 | 0,135 | 0 | Intermediate | INSIG1 |
| GRASP2 | 0 | 2,62131108 | 0,784 | 0,157 | 0 | Intermediate | GRASP |
| KYNU1 | 0 | 2,89597732 | 0,7 | 0,078 | 0 | Intermediate | KYNU |
| FCGRT1 | 0 | 3,16318297 | 0,679 | 0,068 | 0 | Intermediate | FCGRT |
| CD861 | 0 | 4,04152571 | 0,639 | 0,042 | 0 | Intermediate | CD86 |
| LST11 | 0 | 2,41330704 | 0,67 | 0,074 | 0 | Intermediate | LST1 |
| ETS21 | 0 | 2,93075487 | 0,672 | 0,086 | 0 | Intermediate | ETS2 |
| TIMP11 | 0 | 2,56604006 | 0,747 | 0,165 | 0 | Intermediate | TIMP1 |
| NAMPT1 | 0 | 1,68972827 | 0,878 | 0,297 | 0 | Intermediate | NAMPT |
| IRAK31 | 0 | 2,19446458 | 0,682 | 0,108 | 0 | Intermediate | IRAK3 |
| GABARAP1 | 0 | 2,15492926 | 0,783 | 0,21 | 0 | Intermediate | GABARAP |
| STX111 | 0 | 2,67436513 | 0,687 | 0,118 | 0 | Intermediate | STX11 |
| ANXA51 | 0 | 2,30362662 | 0,726 | 0,161 | 0 | Intermediate | ANXA5 |
| SAT11 | 0 | 2,34858554 | 0,956 | 0,392 | 0 | Intermediate | SAT1 |
| CTSS1 | 0 | 1,52002898 | 0,741 | 0,178 | 0 | Intermediate | CTSS |
| PLSCR11 | 0 | 3,40914813 | 0,623 | 0,065 | 0 | Intermediate | PLSCR1 |
| COTL11 | 0 | 1,84659707 | 0,795 | 0,239 | 0 | Intermediate | COTL1 |
| FCGR2B1 | 0 | 4,6292376 | 0,577 | 0,025 | 0 | Intermediate | FCGR2B |
| GPR1831 | 0 | 2,67843486 | 0,92 | 0,37 | 0 | Intermediate | GPR183 |
| FCGR2A1 | 0 | 3,42170926 | 0,603 | 0,055 | 0 | Intermediate | FCGR2A |
| VEGFA1 | 0 | 3,59495849 | 0,595 | 0,048 | 0 | Intermediate | VEGFA |
| FAM49A1 | 0 | 2,50834637 | 0,66 | 0,114 | 0 | Intermediate | FAM49A |
| RILPL21 | 0 | 2,08837893 | 0,765 | 0,226 | 0 | Intermediate | RILPL2 |
| RAB311 | 0 | 2,83902115 | 0,608 | 0,072 | 0 | Intermediate | RAB31 |
| MAP3K83 | 0 | 1,98516237 | 0,795 | 0,268 | 0 | Intermediate | MAP3K8 |
| MIR181A1HG | 0 | 3,37095708 | 0,584 | 0,075 | 0 | Intermediate | MIR181A1HG |
| NME21 | 0 | 1,7745724 | 0,749 | 0,242 | 0 | Intermediate | NME2 |
| HLA-DMB1 | 0 | 3,81318952 | 0,543 | 0,04 | 0 | Intermediate | HLA-DMB |
| IL13RA11 | 0 | 3,89691263 | 0,539 | 0,036 | 0 | Intermediate | IL13RA1 |
| CFP1 | 0 | 4,28230595 | 0,529 | 0,03 | 0 | Intermediate | CFP |
| RGS21 | 0 | 2,47172337 | 0,675 | 0,181 | 0 | Intermediate | RGS2 |
| GSTP13 | 0 | 1,93019723 | 0,697 | 0,205 | 0 | Intermediate | GSTP1 |
| IER31 | 0 | 2,7068329 | 0,574 | 0,082 | 0 | Intermediate | IER3 |
| NR4A31 | 0 | 2,25930618 | 0,716 | 0,225 | 0 | Intermediate | NR4A3 |
| PHACTR11 | 0 | 3,4312488 | 0,542 | 0,052 | 0 | Intermediate | PHACTR1 |
| GNA121 | 0 | 3,5473115 | 0,545 | 0,059 | 0 | Intermediate | GNA12 |
| YBX32 | 0 | 2,14005011 | 0,633 | 0,147 | 0 | Intermediate | YBX3 |
| PLXDC21 | 0 | 2,58140579 | 0,564 | 0,078 | 0 | Intermediate | PLXDC2 |
| AP1S21 | 0 | 2,57131891 | 0,593 | 0,109 | 0 | Intermediate | AP1S2 |
| HLA-DQA21 | 0 | 4,46031867 | 0,534 | 0,053 | 0 | Intermediate | HLA-DQA2 |
| MCTP11 | 0 | 2,3123146 | 0,558 | 0,079 | 0 | Intermediate | MCTP1 |

| row.names | p_val | avg_log2FC | pct.1 | pct.2 | p_val_ac | cluster | gene |
|---|---|---|---|---|---|---|---|
| LINC026942 | 0 | 4,49929644 | 0,745 | 0,076 | 0 | Tregs | LINC02694 |
| AL136456.11 | 0 | 5,14101181 | 0,72 | 0,051 | 0 | Tregs | AL136456.1 |
| CTLA4 | 0 | 4,51225162 | 0,692 | 0,049 | 0 | Tregs | CTLA4 |
| IKZF22 | 0 | 3,52457613 | 0,735 | 0,097 | 0 | Tregs | IKZF2 |
| BATF2 | 0 | 3,26567833 | 0,743 | 0,14 | 0 | Tregs | BATF |
| STAM | 0 | 3,59969162 | 0,673 | 0,094 | 0 | Tregs | STAM |
| TBC1D4 | 0 | 4,69229177 | 0,596 | 0,035 | 0 | Tregs | TBC1D4 |
| UGP22 | 0 | 2,89990503 | 0,717 | 0,169 | 0 | Tregs | UGP2 |
| TOX | 0 | 3,35214861 | 0,697 | 0,15 | 0 | Tregs | TOX |
| TIGIT1 | 0 | 4,07293996 | 0,553 | 0,046 | 0 | Tregs | TIGIT |
| HPGD1 | 0 | 3,87989384 | 0,583 | 0,09 | 0 | Tregs | HPGD |
| ICOS2 | 0 | 2,63169046 | 0,615 | 0,143 | 0 | Tregs | ICOS |
| AC013652.12 | 0 | 3,62056907 | 0,519 | 0,052 | 0 | Tregs | AC013652.1 |
| LDLRAD44 | 0 | 2,20827183 | 0,656 | 0,218 | 0 | Tregs | LDLRAD4 |
| PHACTR21 | 0 | 2,13399923 | 0,62 | 0,202 | 0 | Tregs | PHACTR2 |
| ZNRF2 | 0 | 2,37880141 | 0,567 | 0,159 | 0 | Tregs | ZNRF2 |
| SKAP11 | 0 | 1,78438227 | 0,737 | 0,337 | 0 | Tregs | SKAP1 |
| TRAC2 | 0 | 2,0728069 | 0,592 | 0,198 | 0 | Tregs | TRAC |
| PHTF22 | 0 | 2,40631281 | 0,527 | 0,136 | 0 | Tregs | PHTF2 |
| CARD162 | 0 | 3,0942738 | 0,469 | 0,083 | 0 | Tregs | CARD16 |
| VAV33 | 0 | 1,81771218 | 0,69 | 0,308 | 0 | Tregs | VAV3 |
| DUSP162 | 0 | 2,41065323 | 0,521 | 0,14 | 0 | Tregs | DUSP16 |
| IL321 | 0 | 2,17471617 | 0,871 | 0,494 | 0 | Tregs | IL32 |
| RTKN2 | 0 | 4,86992824 | 0,395 | 0,02 | 0 | Tregs | RTKN2 |
| SLAMF11 | 0 | 3,04674927 | 0,436 | 0,065 | 0 | Tregs | SLAMF1 |
| BTG3 | 0 | 2,42463326 | 0,513 | 0,144 | 0 | Tregs | BTG3 |
| IL2RA1 | 0 | 3,82911797 | 0,385 | 0,035 | 0 | Tregs | IL2RA |
| KAT2B2 | 0 | 2,12063253 | 0,533 | 0,184 | 0 | Tregs | KAT2B |
| CASK1 | 0 | 1,75935008 | 0,54 | 0,196 | 0 | Tregs | CASK |
| PMAIP13 | 0 | 2,42144424 | 0,46 | 0,119 | 0 | Tregs | PMAIP1 |
| RAP1A1 | 0 | 1,68096668 | 0,65 | 0,311 | 0 | Tregs | RAP1A |
| PLCL12 | 0 | 3,28073711 | 0,397 | 0,06 | 0 | Tregs | PLCL1 |
| TNFRSF91 | 0 | 3,37568392 | 0,374 | 0,04 | 0 | Tregs | TNFRSF9 |
| ZNF2921 | 0 | 1,97255802 | 0,521 | 0,188 | 0 | Tregs | ZNF292 |
| SGMS11 | 0 | 2,56919381 | 0,436 | 0,107 | 0 | Tregs | SGMS1 |
| NCK21 | 0 | 1,83312534 | 0,498 | 0,173 | 0 | Tregs | NCK2 |
| LINC02195 | 0 | 6,44595284 | 0,325 | 0,005 | 0 | Tregs | LINC02195 |
| TNFRSF18 | 0 | 2,69919604 | 0,392 | 0,075 | 0 | Tregs | TNFRSF18 |
| TRAF31 | 0 | 2,19669273 | 0,449 | 0,136 | 0 | Tregs | TRAF3 |
| MAP3K51 | 0 | 1,99886883 | 0,482 | 0,175 | 0 | Tregs | MAP3K5 |
| MAST41 | 0 | 3,66274556 | 0,332 | 0,034 | 0 | Tregs | MAST4 |
| USP15 | 0 | 1,60200993 | 0,714 | 0,42 | 0 | Tregs | USP15 |
| GBP21 | 0 | 1,9075695 | 0,42 | 0,127 | 0 | Tregs | GBP2 |
| CTSC1 | 0 | 1,87172668 | 0,46 | 0,168 | 0 | Tregs | CTSC |
| HIVEP12 | 0 | 1,90219599 | 0,418 | 0,14 | 0 | Tregs | HIVEP1 |
| DUSP44 | 0 | 1,83795283 | 0,415 | 0,138 | 0 | Tregs | DUSP4 |
| GBP51 | 0 | 2,4185961 | 0,346 | 0,073 | 0 | Tregs | GBP5 |
| GLRX1 | 0 | 2,26771872 | 0,371 | 0,106 | 0 | Tregs | GLRX |
| CD272 | 0 | 3,28230077 | 0,297 | 0,035 | 0 | Tregs | CD27 |
| LAYN1 | 0 | 6,02407642 | 0,265 | 0,004 | 0 | Tregs | LAYN |
| TTN | 0 | 3,8470477 | 0,274 | 0,023 | 0 | Tregs | TTN |
| PVT12 | 0 | 2,71755929 | 0,306 | 0,055 | 0 | Tregs | PVT1 |
| AC093865.12 | 0 | 3,45035332 | 0,274 | 0,03 | 0 | Tregs | AC093865.1 |
| ZC2HC1A | 0 | 5,36951289 | 0,238 | 0,008 | 0 | Tregs | ZC2HC1A |
| SETD72 | 0 | 2,62540536 | 0,28 | 0,052 | 0 | Tregs | SETD7 |
| CORO1B2 | 0 | 2,53590217 | 0,284 | 0,06 | 0 | Tregs | CORO1B |
| LINC019431 | 0 | 3,62858839 | 0,241 | 0,024 | 0 | Tregs | LINC01943 |
| FOXP3 | 0 | 8,19935726 | 0,214 | 0,001 | 0 | Tregs | FOXP3 |
| TMEM154 | 0 | 2,61721798 | 0,261 | 0,049 | 0 | Tregs | TMEM154 |
| NAB11 | 0 | 3,06101127 | 0,244 | 0,035 | 0 | Tregs | NAB1 |
| PARD6G2 | 0 | 2,36187844 | 0,27 | 0,062 | 0 | Tregs | PARD6G |
| FANK1 | 0 | 8,09722448 | 0,199 | 0,001 | 0 | Tregs | FANK1 |
| CCNG21 | 0 | 3,08117371 | 0,224 | 0,03 | 0 | Tregs | CCNG2 |
| SPATS2L | 0 | 2,4635278 | 0,238 | 0,047 | 0 | Tregs | SPATS2L |
| ENTPD12 | 0 | 2,69283238 | 0,228 | 0,037 | 0 | Tregs | ENTPD1 |
| CCDC141 | 0 | 4,64409387 | 0,194 | 0,009 | 0 | Tregs | CCDC141 |
| ZC3H12D2 | 0 | 2,49378399 | 0,222 | 0,046 | 0 | Tregs | ZC3H12D |
| ICA1 | 0 | 5,40826652 | 0,166 | 0,005 | 0 | Tregs | ICA1 |
| S100A42 | 0 | 1,30023914 | 0,888 | 0,738 | 0 | Tregs | S100A4 |
| HTATIP21 | 0 | 2,78920728 | 0,178 | 0,028 | 0 | Tregs | HTATIP2 |
| CCR8 | 0 | 7,96415046 | 0,148 | 0,001 | 0 | Tregs | CCR8 |
| CD177 | 0 | 11,3502734 | 0,147 | 0 | 0 | Tregs | CD177 |
| DNPH1 | 0 | 3,43316355 | 0,151 | 0,015 | 0 | Tregs | DNPH1 |
| CPNE21 | 0 | 4,51715762 | 0,142 | 0,006 | 0 | Tregs | CPNE2 |

| row.names | p_val | avg_log2FC | pct.1 | pct.2 | p_val_adj | cluster | gene |
|---|---|---|---|---|---|---|---|
| NKG73 | 0 | 1,32900087 | 0,956 | 0,397 | 0 | T/NK-b | NKG7 |
| GNLY3 | 0 | 1,02456273 | 0,801 | 0,279 | 0 | T/NK-b | GNLY |
| CCL42 | 0 | 2,00631899 | 0,891 | 0,379 | 0 | T/NK-b | CCL4 |
| KLRD13 | 0 | 1,66043462 | 0,774 | 0,265 | 0 | T/NK-b | KLRD1 |
| GZMH3 | 0 | 2,064077 | 0,716 | 0,214 | 0 | T/NK-b | GZMH |
| GZMB3 | 0 | 0,66743064 | 0,699 | 0,263 | 0 | T/NK-b | GZMB |
| CCL53 | 0 | 1,07736171 | 0,948 | 0,539 | 0 | T/NK-b | CCL5 |
| FGFBP23 | 0 | 1,30844869 | 0,537 | 0,16 | 0 | T/NK-b | FGFBP2 |
| CD3G2 | 0 | 1,70443607 | 0,605 | 0,242 | 0 | T/NK-b | CD3G |
| CD8A2 | 0 | 1,82856437 | 0,485 | 0,147 | 0 | T/NK-b | CD8A |
| TRGC22 | 0 | 2,1728007 | 0,383 | 0,099 | 0 | T/NK-b | TRGC2 |
| AC243829.23 | 0 | 2,84499551 | 0,327 | 0,055 | 0 | T/NK-b | AC243829.2 |
| IFNG3 | 0 | 2,32356734 | 0,317 | 0,073 | 0 | T/NK-b | IFNG |
| SGCD3 | 4,147E-303 | 2,51820136 | 0,208 | 0,041 | 1,035E-298 | T/NK-b | SGCD |
| CCL4L22 | 1,205E-297 | 1,927508 | 0,383 | 0,122 | 3,006E-293 | T/NK-b | CCL4L2 |
| ZEB24 | 4,06E-294 | 1,04972828 | 0,811 | 0,434 | 1,013E-289 | T/NK-b | ZEB2 |
| ARRDC33 | 4,194E-257 | 1,85010742 | 0,347 | 0,114 | 1,047E-252 | T/NK-b | ARRDC3 |
| KLRK14 | 3,572E-252 | 1,37646559 | 0,48 | 0,191 | 8,915E-248 | T/NK-b | KLRK1 |
| ADGRG13 | 1,598E-239 | 2,00637052 | 0,233 | 0,059 | 3,989E-235 | T/NK-b | ADGRG1 |
| PPP2R5C4 | 1,155E-227 | 0,84127573 | 0,873 | 0,596 | 2,883E-223 | T/NK-b | PPP2R5C |
| CTSW3 | 4,027E-226 | 1,02636295 | 0,576 | 0,261 | 1,005E-221 | T/NK-b | CTSW |
| AC016074.21 | 5,533E-224 | 2,23793499 | 0,209 | 0,052 | 1,381E-219 | T/NK-b | AC016074.2 |
| AC022075.12 | 4,124E-216 | 2,35885243 | 0,163 | 0,034 | 1,029E-211 | T/NK-b | AC022075.1 |
| KLF21 | 1,022E-215 | 0,9644346 | 0,748 | 0,435 | 2,55E-211 | T/NK-b | KLF2 |
| CD8B2 | 4,531E-206 | 1,79739618 | 0,263 | 0,08 | 1,131E-201 | T/NK-b | CD8B |
| CD3D4 | 1,497E-194 | 1,15513927 | 0,577 | 0,301 | 3,736E-190 | T/NK-b | CD3D |
| SYNE13 | 1,382E-179 | 1,1711671 | 0,446 | 0,198 | 3,449E-175 | T/NK-b | SYNE1 |
| KLRG13 | 3,945E-175 | 1,7616582 | 0,252 | 0,083 | 9,845E-171 | T/NK-b | KLRG1 |
| TRG-AS13 | 1,39E-174 | 1,77287254 | 0,254 | 0,084 | 3,468E-170 | T/NK-b | TRG-AS1 |
| SPON23 | 1,811E-165 | 0,87854263 | 0,416 | 0,175 | 4,519E-161 | T/NK-b | SPON2 |
| PYHIN13 | 1,058E-164 | 1,14371722 | 0,413 | 0,182 | 2,639E-160 | T/NK-b | PYHIN1 |
| C1orf213 | 7,059E-162 | 0,96095758 | 0,408 | 0,173 | 1,762E-157 | T/NK-b | C1orf21 |
| PATL22 | 6,745E-154 | 2,20903037 | 0,127 | 0,028 | 1,683E-149 | T/NK-b | PATL2 |
| PRKCH3 | 6,821E-149 | 0,72773979 | 0,718 | 0,446 | 1,702E-144 | T/NK-b | PRKCH |
| AC015849.12 | 5,345E-145 | 2,52684215 | 0,094 | 0,017 | 1,334E-140 | T/NK-b | AC015849.1 |
| FCGR3A3 | 7,892E-143 | 1,07868 | 0,336 | 0,138 | 1,97E-138 | T/NK-b | FCGR3A |
| SAMD34 | 5,091E-139 | 0,91741261 | 0,467 | 0,232 | 1,27E-134 | T/NK-b | SAMD3 |
| AKNA4 | 3,395E-137 | 0,83409843 | 0,6 | 0,344 | 8,473E-133 | T/NK-b | AKNA |
| LINC023843 | 1,011E-133 | 1,91510191 | 0,142 | 0,038 | 2,523E-129 | T/NK-b | LINC02384 |
| CST74 | 7,031E-133 | 0,56203635 | 0,748 | 0,452 | 1,755E-128 | T/NK-b | CST7 |
| PTP4A22 | 1,48E-132 | 1,04176897 | 0,473 | 0,255 | 3,693E-128 | T/NK-b | PTP4A2 |
| LINC008611 | 2,409E-130 | 1,61240086 | 0,216 | 0,076 | 6,011E-126 | T/NK-b | LINC00861 |
| YES13 | 4,6E-124 | 1,11920013 | 0,338 | 0,152 | 1,148E-119 | T/NK-b | YES1 |
| PTPRC2 | 5,409E-123 | 0,52533507 | 0,964 | 0,816 | 1,35E-118 | T/NK-b | PTPRC |
| HCST2 | 1,88E-115 | 0,80282682 | 0,698 | 0,487 | 4,693E-111 | T/NK-b | HCST |
| AL158071.13 | 3,73E-115 | 2,61662528 | 0,069 | 0,012 | 9,309E-111 | T/NK-b | AL158071.1 |
| ID22 | 1,848E-114 | 0,76639209 | 0,653 | 0,425 | 4,611E-110 | T/NK-b | ID2 |
| ITGA43 | 1,385E-112 | 0,82645183 | 0,543 | 0,321 | 3,456E-108 | T/NK-b | ITGA4 |
| C12orf754 | 2,041E-112 | 1,05532628 | 0,331 | 0,154 | 5,094E-108 | T/NK-b | C12orf75 |
| TGFBR33 | 5,145E-112 | 0,86109825 | 0,419 | 0,216 | 1,284E-107 | T/NK-b | TGFBR3 |
| CD692 | 5,264E-111 | 0,50596574 | 0,716 | 0,475 | 1,314E-106 | T/NK-b | CD69 |
| AL096794.13 | 5,959E-110 | 2,2626792 | 0,093 | 0,021 | 1,487E-105 | T/NK-b | AL096794.1 |
| GZMA2 | 1,263E-108 | 0,7450255 | 0,567 | 0,345 | 3,151E-104 | T/NK-b | GZMA |
| CEP783 | 4,486E-106 | 1,32370354 | 0,221 | 0,087 | 1,12E-101 | T/NK-b | CEP78 |
| PIP4K2A3 | 8,727E-102 | 0,75426797 | 0,606 | 0,395 | 2,178E-97 | T/NK-b | PIP4K2A |
| ABHD17A3 | 2,316E-100 | 0,92997162 | 0,379 | 0,195 | 5,7789E-96 | T/NK-b | ABHD17A |
| ITGAL3 | 6,154E-99 | 1,10258595 | 0,29 | 0,134 | 1,5357E-94 | T/NK-b | ITGAL |
| ARGLU11 | 8,3429E-98 | 0,8716169 | 0,467 | 0,273 | 2,0821E-93 | T/NK-b | ARGLU1 |
| FCRL63 | 8,3262E-96 | 1,7186643 | 0,129 | 0,04 | 2,0779E-91 | T/NK-b | FCRL6 |
| TRGC14 | 1,4052E-95 | 1,45186872 | 0,204 | 0,081 | 3,5068E-91 | T/NK-b | TRGC1 |
| PRF13 | 3,1486E-93 | 0,55705046 | 0,439 | 0,231 | 7,8576E-89 | T/NK-b | PRF1 |
| BIN23 | 5,5133E-93 | 1,00386823 | 0,332 | 0,168 | 1,3759E-88 | T/NK-b | BIN2 |
| SLC10A11 | 6,904E-92 | 2,22493891 | 0,077 | 0,017 | 1,723E-87 | T/NK-b | SLC10A1 |
| TRBV282 | 1,1092E-91 | 2,09712178 | 0,087 | 0,021 | 2,768E-87 | T/NK-b | TRBV28 |
| IKZF23 | 8,7606E-91 | 1,63614962 | 0,25 | 0,116 | 2,1863E-86 | T/NK-b | IKZF2 |
| STPG41 | 2,259E-90 | 2,42366384 | 0,074 | 0,016 | 5,6375E-86 | T/NK-b | STPG4 |
| HLA-F3 | 3,2979E-90 | 0,8763441 | 0,432 | 0,25 | 8,2301E-86 | T/NK-b | HLA-F |
| GZMM4 | 1,0159E-89 | 0,74333944 | 0,45 | 0,255 | 2,5354E-85 | T/NK-b | GZMM |
| LUC7L32 | 1,0834E-89 | 1,00985611 | 0,352 | 0,187 | 2,7038E-85 | T/NK-b | LUC7L3 |
| RABGAP1L2 | 1,4156E-89 | 0,76342751 | 0,627 | 0,442 | 3,5328E-85 | T/NK-b | RABGAP1L |
| GALM3 | 1,1682E-88 | 1,20912133 | 0,28 | 0,135 | 2,9154E-84 | T/NK-b | GALM |
| TAFA12 | 1,5687E-88 | 1,66893527 | 0,133 | 0,043 | 3,9149E-84 | T/NK-b | TAFA1 |
| TNF1 | 1,6118E-88 | 1,49205516 | 0,172 | 0,065 | 4,0224E-84 | T/NK-b | TNF |
| TXNIP3 | 1,9812E-88 | 0,54271867 | 0,784 | 0,595 | 4,9443E-84 | T/NK-b | TXNIP |

| row.names | p_val | avg_log2FC | pct.1 | pct.2 | p_val_adj | cluster | gene |
|---|---|---|---|---|---|---|---|
| KLRB13 | 0 | 2,46225552 | 0,925 | 0,335 | 0 | MAIT | KLRB1 |
| GZMK2 | 0 | 1,32558408 | 0,732 | 0,244 | 0 | MAIT | GZMK |
| IL7R3 | 0 | 1,31731534 | 0,973 | 0,501 | 0 | MAIT | IL7R |
| RORA2 | 0 | 1,81756455 | 0,891 | 0,43 | 0 | MAIT | RORA |
| SLC4A101 | 0 | 5,54694469 | 0,427 | 0,011 | 0 | MAIT | SLC4A10 |
| PHACTR22 | 0 | 2,01243764 | 0,615 | 0,203 | 0 | MAIT | PHACTR2 |
| CCR63 | 0 | 2,27503797 | 0,477 | 0,094 | 0 | MAIT | CCR6 |
| ERN11 | 0 | 1,82372254 | 0,56 | 0,193 | 0 | MAIT | ERN1 |
| MYBL13 | 0 | 1,70507891 | 0,588 | 0,221 | 0 | MAIT | MYBL1 |
| RBMS12 | 0 | 1,37204983 | 0,659 | 0,299 | 0 | MAIT | RBMS1 |
| ZBTB163 | 0 | 1,67537069 | 0,55 | 0,191 | 0 | MAIT | ZBTB16 |
| TLE12 | 0 | 2,02398855 | 0,462 | 0,113 | 0 | MAIT | TLE1 |
| SPOCK23 | 0 | 1,39676565 | 0,636 | 0,288 | 0 | MAIT | SPOCK2 |
| CAMK44 | 0 | 1,48667221 | 0,567 | 0,22 | 0 | MAIT | CAMK4 |
| MGAT4A3 | 0 | 1,72791048 | 0,526 | 0,181 | 0 | MAIT | MGAT4A |
| TC2N3 | 0 | 1,65025878 | 0,575 | 0,236 | 0 | MAIT | TC2N |
| TTC39C3 | 0 | 1,85088373 | 0,455 | 0,148 | 0 | MAIT | TTC39C |
| SYTL23 | 0 | 1,74097166 | 0,455 | 0,154 | 0 | MAIT | SYTL2 |
| RORA-AS13 | 0 | 2,07844728 | 0,422 | 0,125 | 0 | MAIT | RORA-AS1 |
| ODF2L2 | 0 | 1,96232744 | 0,416 | 0,124 | 0 | MAIT | ODF2L |
| RUNX22 | 0 | 1,91620065 | 0,387 | 0,103 | 0 | MAIT | RUNX2 |
| KLRG14 | 0 | 2,2548091 | 0,353 | 0,079 | 0 | MAIT | KLRG1 |
| P2RY141 | 0 | 2,8621141 | 0,223 | 0,034 | 0 | MAIT | P2RY14 |
| KIF5C1 | 0 | 2,99392911 | 0,191 | 0,026 | 0 | MAIT | KIF5C |
| ADAM122 | 0 | 4,15121133 | 0,168 | 0,011 | 0 | MAIT | ADAM12 |
| NTN43 | 0 | 3,30321442 | 0,145 | 0,015 | 0 | MAIT | NTN4 |
| LINC016441 | 0 | 4,42627421 | 0,116 | 0,005 | 0 | MAIT | LINC01644 |
| ME11 | 0 | 3,58062929 | 0,111 | 0,008 | 0 | MAIT | ME1 |
| LTK1 | 0 | 4,21768759 | 0,094 | 0,005 | 0 | MAIT | LTK |
| IL23R2 | 0 | 4,36638345 | 0,083 | 0,004 | 0 | MAIT | IL23R |
| RORC1 | 0 | 4,43658658 | 0,068 | 0,003 | 0 | MAIT | RORC |
| AC010967.1 | 0 | 6,55503002 | 0,044 | 0 | 0 | MAIT | AC010967.1 |
| PDE4D2 | 2E-307 | 1,2872752 | 0,678 | 0,326 | 6E-303 | MAIT | PDE4D |
| DUSP163 | 6E-299 | 1,77253364 | 0,424 | 0,144 | 1,5E-294 | MAIT | DUSP16 |
| NCR32 | 2E-290 | 2,21048624 | 0,251 | 0,058 | 5,6E-286 | MAIT | NCR3 |
| WDFY21 | 3E-276 | 1,83880714 | 0,399 | 0,135 | 7,9E-272 | MAIT | WDFY2 |
| GPR654 | 7E-272 | 1,46519018 | 0,504 | 0,2 | 1,7E-267 | MAIT | GPR65 |
| CYTH32 | 1E-269 | 1,7048815 | 0,363 | 0,114 | 2,5E-265 | MAIT | CYTH3 |
| GTF3C14 | 4E-254 | 1,81269677 | 0,306 | 0,089 | 1,1E-249 | MAIT | GTF3C1 |
| PARP83 | 1E-252 | 0,997064 | 0,862 | 0,586 | 3,6E-248 | MAIT | PARP8 |
| EEF1A11 | 4E-242 | 0,50072519 | 0,984 | 0,974 | 1E-237 | MAIT | EEF1A1 |
| LTB3 | 4E-239 | 1,27995668 | 0,484 | 0,194 | 1E-234 | MAIT | LTB |
| RPLP01 | 6E-239 | 0,84937636 | 0,888 | 0,685 | 1,5E-234 | MAIT | RPLP0 |
| COLQ3 | 5E-235 | 2,57293466 | 0,149 | 0,027 | 1,1E-230 | MAIT | COLQ |
| MPZL33 | 1E-232 | 1,57057734 | 0,401 | 0,148 | 3,6E-228 | MAIT | MPZL3 |
| AL136456.12 | 4E-227 | 1,25104658 | 0,255 | 0,069 | 1,1E-222 | MAIT | AL136456.1 |
| SUSD41 | 8E-227 | 4,25401723 | 0,047 | 0,003 | 2,1E-222 | MAIT | SUSD4 |
| ROR2 | 1E-224 | 4,52133641 | 0,038 | 0,002 | 2,7E-220 | MAIT | ROR2 |
| SLC7A52 | 2E-220 | 1,43823387 | 0,436 | 0,175 | 4,1E-216 | MAIT | SLC7A5 |
| RPLP11 | 8E-218 | 0,54801659 | 0,977 | 0,964 | 2E-213 | MAIT | RPLP1 |
| LINC018712 | 4E-213 | 1,26755347 | 0,514 | 0,231 | 1,1E-208 | MAIT | LINC01871 |
| CEBPD2 | 2E-210 | 0,88692504 | 0,504 | 0,21 | 4,5E-206 | MAIT | CEBPD |
| DLEU13 | 2E-207 | 1,98381397 | 0,3 | 0,1 | 6,1E-203 | MAIT | DLEU1 |
| DPP43 | 2E-203 | 1,89174984 | 0,236 | 0,066 | 5E-199 | MAIT | DPP4 |
| HPGD2 | 2E-201 | 1,22004323 | 0,308 | 0,1 | 4,4E-197 | MAIT | HPGD |
| AQP33 | 9E-193 | 1,93711989 | 0,214 | 0,058 | 2,2E-188 | MAIT | AQP3 |
| PTPRM1 | 4E-191 | 2,59271 | 0,189 | 0,048 | 1,1E-186 | MAIT | PTPRM |
| NEK73 | 7E-188 | 1,27072097 | 0,524 | 0,262 | 1,8E-183 | MAIT | NEK7 |
| TNFAIP33 | 2E-185 | 0,95432373 | 0,832 | 0,618 | 3,9E-181 | MAIT | TNFAIP3 |
| NEO11 | 3E-185 | 2,27698113 | 0,148 | 0,032 | 6,7E-181 | MAIT | NEO1 |
| IKZF24 | 3E-184 | 1,09439264 | 0,323 | 0,113 | 6,5E-180 | MAIT | IKZF2 |
| A2M2 | 3E-184 | 1,75509203 | 0,234 | 0,069 | 7,4E-180 | MAIT | A2M |
| PBX43 | 1E-182 | 1,10783397 | 0,55 | 0,274 | 2,4E-178 | MAIT | PBX4 |
| RBM24 | 4E-175 | 8,78486077 | 0,013 | 0 | 9,3E-171 | MAIT | RBM24 |
| LINC011381 | 6E-172 | 1,56726554 | 0,294 | 0,105 | 1,6E-167 | MAIT | LINC01138 |
| MAP3K41 | 3E-171 | 1,74366292 | 0,253 | 0,083 | 8,7E-167 | MAIT | MAP3K4 |
| CERK1 | 7E-171 | 1,59800371 | 0,267 | 0,09 | 1,8E-166 | MAIT | CERK |
| TOB13 | 2E-163 | 1,06587318 | 0,518 | 0,263 | 3,8E-159 | MAIT | TOB1 |
| CELF21 | 7E-160 | 0,92153181 | 0,759 | 0,533 | 1,6E-155 | MAIT | CELF2 |
| CD8A3 | 1E-159 | 1,12338175 | 0,372 | 0,152 | 3,5E-155 | MAIT | CD8A |
| CA22 | 9E-159 | 3,21445103 | 0,054 | 0,005 | 2,3E-154 | MAIT | CA2 |
| SPRY12 | 3E-156 | 1,93103142 | 0,144 | 0,034 | 8,6E-152 | MAIT | SPRY1 |
| CDC14A2 | 2E-155 | 1,07556277 | 0,518 | 0,268 | 5,6E-151 | MAIT | CDC14A |
| TRGC23 | 2E-151 | 1,41718243 | 0,28 | 0,103 | 6E-147 | MAIT | TRGC2 |

| row.names | p_val | avg_log2FC | pct.1 | pct.2 | p_val_adj | cluster | gene |
|---|---|---|---|---|---|---|---|
| XCL11 | 0 | 4,34748516 | 0,685 | 0,058 | 0 | Resident NK | XCL1 |
| XCL23 | 0 | 3,69537502 | 0,713 | 0,1 | 0 | Resident NK | XCL2 |
| KLRD14 | 0 | 1,46618579 | 0,789 | 0,266 | 0 | Resident NK | KLRD1 |
| MCTP23 | 0 | 2,69627516 | 0,674 | 0,162 | 0 | Resident NK | MCTP2 |
| NKG74 | 0 | 0,99473946 | 0,912 | 0,4 | 0 | Resident NK | NKG7 |
| TYROBP4 | 0 | 1,39003637 | 0,77 | 0,274 | 0 | Resident NK | TYROBP |
| CTSW4 | 0 | 1,88686553 | 0,74 | 0,257 | 0 | Resident NK | CTSW |
| KLRC15 | 0 | 3,41481037 | 0,516 | 0,054 | 0 | Resident NK | KLRC1 |
| AREG4 | 0 | 1,47648261 | 0,781 | 0,341 | 0 | Resident NK | AREG |
| GZMK3 | 0 | 1,42034961 | 0,674 | 0,246 | 0 | Resident NK | GZMK |
| CMC14 | 0 | 2,33481939 | 0,579 | 0,164 | 0 | Resident NK | CMC1 |
| IL12RB23 | 0 | 2,33106492 | 0,522 | 0,124 | 0 | Resident NK | IL12RB2 |
| FCER1G4 | 0 | 1,14661449 | 0,596 | 0,201 | 0 | Resident NK | FCER1G |
| REL2 | 0 | 1,95160404 | 0,804 | 0,415 | 0 | Resident NK | REL |
| NFKB12 | 0 | 1,61313435 | 0,813 | 0,427 | 0 | Resident NK | NFKB1 |
| KLRC35 | 0 | 2,36254679 | 0,444 | 0,093 | 0 | Resident NK | KLRC3 |
| TNFRSF181 | 0 | 2,57135989 | 0,408 | 0,076 | 0 | Resident NK | TNFRSF18 |
| MATK4 | 0 | 2,26749228 | 0,402 | 0,093 | 0 | Resident NK | MATK |
| HOPX3 | 0 | 1,7779222 | 0,452 | 0,148 | 0 | Resident NK | HOPX |
| TXK3 | 0 | 2,31729535 | 0,387 | 0,092 | 0 | Resident NK | TXK |
| IL2RB2 | 0 | 2,25008492 | 0,381 | 0,089 | 0 | Resident NK | IL2RB |
| RIN33 | 0 | 1,96075313 | 0,398 | 0,109 | 0 | Resident NK | RIN3 |
| KCNQ51 | 0 | 3,03332977 | 0,342 | 0,055 | 0 | Resident NK | KCNQ5 |
| ATP8B43 | 0 | 3,2007616 | 0,319 | 0,042 | 0 | Resident NK | ATP8B4 |
| CRTAM2 | 0 | 2,46209803 | 0,338 | 0,086 | 0 | Resident NK | CRTAM |
| HIP12 | 0 | 2,61514269 | 0,303 | 0,064 | 0 | Resident NK | HIP1 |
| TIAM13 | 0 | 2,35681884 | 0,322 | 0,083 | 0 | Resident NK | TIAM1 |
| TRDC3 | 0 | 2,49824998 | 0,286 | 0,052 | 0 | Resident NK | TRDC |
| GOLIM41 | 0 | 2,79607883 | 0,268 | 0,04 | 0 | Resident NK | GOLIM4 |
| NCAM14 | 0 | 2,38506408 | 0,261 | 0,051 | 0 | Resident NK | NCAM1 |
| B3GNT72 | 0 | 3,09317338 | 0,239 | 0,029 | 0 | Resident NK | B3GNT7 |
| RASSF81 | 0 | 3,18840619 | 0,23 | 0,029 | 0 | Resident NK | RASSF8 |
| LAT23 | 0 | 2,89914063 | 0,225 | 0,032 | 0 | Resident NK | LAT2 |
| SLFN132 | 0 | 2,65834515 | 0,221 | 0,038 | 0 | Resident NK | SLFN13 |
| PHLDB23 | 0 | 3,41075115 | 0,211 | 0,03 | 0 | Resident NK | PHLDB2 |
| LDB21 | 0 | 4,1225742 | 0,16 | 0,008 | 0 | Resident NK | LDB2 |
| TNFRSF11A1 | 0 | 4,69888533 | 0,142 | 0,006 | 0 | Resident NK | TNFRSF11A |
| GNG42 | 0 | 3,21730043 | 0,152 | 0,017 | 0 | Resident NK | GNG4 |
| CSF2 | 0 | 6,47532697 | 0,132 | 0,003 | 0 | Resident NK | CSF2 |
| PPP1R9A1 | 0 | 4,780453 | 0,133 | 0,005 | 0 | Resident NK | PPP1R9A |
| TOX21 | 0 | 3,29808015 | 0,144 | 0,018 | 0 | Resident NK | TOX2 |
| CCNJL2 | 0 | 4,27432276 | 0,097 | 0,006 | 0 | Resident NK | CCNJL |
| SPTSSB | 0 | 6,10143681 | 0,069 | 0,001 | 0 | Resident NK | SPTSSB |
| BMP2 | 0 | 4,82964188 | 0,057 | 0,002 | 0 | Resident NK | BMP2 |
| MICAL24 | 1E-293 | 2,56601686 | 0,19 | 0,033 | 3E-289 | Resident NK | MICAL2 |
| OTULIN2 | 5E-282 | 2,11182772 | 0,331 | 0,094 | 1E-277 | Resident NK | OTULIN |
| TNFSF14 | 8E-277 | 2,7446241 | 0,174 | 0,03 | 2E-272 | Resident NK | TNFSF14 |
| IFITM22 | 2E-274 | 1,21624916 | 0,761 | 0,446 | 6E-270 | Resident NK | IFITM2 |
| GRASP3 | 5E-273 | 1,76228699 | 0,467 | 0,176 | 1E-268 | Resident NK | GRASP |
| COL4A4 | 7E-272 | 4,49707417 | 0,052 | 0,002 | 2E-267 | Resident NK | COL4A4 |
| ACOXL | 3E-271 | 4,59723364 | 0,054 | 0,003 | 8E-267 | Resident NK | ACOXL |
| CD73 | 5E-271 | 1,33813203 | 0,631 | 0,29 | 1E-266 | Resident NK | CD7 |
| IRF82 | 6E-271 | 2,06726487 | 0,249 | 0,057 | 1E-266 | Resident NK | IRF8 |
| GNLY4 | 4E-270 | 1,34912954 | 0,639 | 0,286 | 1E-265 | Resident NK | GNLY |
| IER2 | 2E-259 | 1,31296621 | 0,705 | 0,394 | 5E-255 | Resident NK | IER2 |
| CAPG2 | 2E-257 | 2,26178119 | 0,22 | 0,048 | 5E-253 | Resident NK | CAPG |
| ARHGAP312 | 4E-254 | 2,18621484 | 0,252 | 0,062 | 9E-250 | Resident NK | ARHGAP31 |
| DHRS33 | 4E-249 | 2,28055695 | 0,22 | 0,05 | 1E-244 | Resident NK | DHRS3 |
| BHLHE403 | 4E-249 | 1,56583423 | 0,495 | 0,203 | 1E-244 | Resident NK | BHLHE40 |
| MAFF3 | 2E-242 | 1,5838589 | 0,406 | 0,143 | 4E-238 | Resident NK | MAFF |
| NCALD3 | 7E-236 | 1,50065711 | 0,514 | 0,22 | 2E-231 | Resident NK | NCALD |
| HS6ST2 | 1E-233 | 4,83793408 | 0,039 | 0,001 | 3E-229 | Resident NK | HS6ST2 |
| MAP3K84 | 2E-229 | 1,21909574 | 0,601 | 0,282 | 4E-225 | Resident NK | MAP3K8 |
| ADGRE52 | 2E-229 | 1,17279897 | 0,733 | 0,437 | 6E-225 | Resident NK | ADGRE5 |
| KLRB14 | 6E-229 | 0,90877943 | 0,712 | 0,344 | 1E-224 | Resident NK | KLRB1 |
| PLCG22 | 2E-227 | 0,92267046 | 0,424 | 0,158 | 4E-223 | Resident NK | PLCG2 |
| IRAK32 | 1E-217 | 1,12051563 | 0,377 | 0,125 | 2E-213 | Resident NK | IRAK3 |
| CD632 | 2E-215 | 1,33418893 | 0,52 | 0,233 | 6E-211 | Resident NK | CD63 |
| SPRY21 | 2E-214 | 3,00940124 | 0,095 | 0,012 | 6E-210 | Resident NK | SPRY2 |
| LINC00996 | 6E-214 | 3,70913393 | 0,058 | 0,004 | 1E-209 | Resident NK | LINC00996 |
| APOBEC3G5 | 2E-213 | 1,90527235 | 0,293 | 0,09 | 5E-209 | Resident NK | APOBEC3G |
| AC017104.12 | 2E-207 | 3,21073205 | 0,082 | 0,009 | 4E-203 | Resident NK | AC017104.1 |
| KIR2DL42 | 4E-202 | 2,73898929 | 0,128 | 0,022 | 1E-197 | Resident NK | KIR2DL4 |
| DUSP21 | 2E-201 | 1,06715411 | 0,798 | 0,543 | 6E-197 | Resident NK | DUSP2 |

| row.names | p_val | avg_log2FC | pct.1 | pct.2 | p_val_a | cluster | gene |
|---|---|---|---|---|---|---|---|
| BANK1 | 0 | 8,39471776 | 0,835 | 0,006 | 0 | Mature B | BANK1 |
| CD79A | 0 | 8,88887175 | 0,789 | 0,003 | 0 | Mature B | CD79A |
| MS4A1 | 0 | 7,3472569 | 0,78 | 0,009 | 0 | Mature B | MS4A1 |
| HLA-DRA2 | 0 | 1,73357128 | 0,973 | 0,221 | 0 | Mature B | HLA-DRA |
| EBF1 | 0 | 7,01457332 | 0,715 | 0,007 | 0 | Mature B | EBF1 |
| ARHGAP241 | 0 | 5,25935526 | 0,736 | 0,035 | 0 | Mature B | ARHGAP24 |
| AFF31 | 0 | 4,64200427 | 0,751 | 0,059 | 0 | Mature B | AFF3 |
| HLA-DQA12 | 0 | 1,754576 | 0,829 | 0,146 | 0 | Mature B | HLA-DQA1 |
| GNG71 | 0 | 6,17310829 | 0,692 | 0,017 | 0 | Mature B | GNG7 |
| HLA-DPB12 | 0 | 1,13144117 | 0,89 | 0,246 | 0 | Mature B | HLA-DPB1 |
| HLA-DRB12 | 0 | 1,24409724 | 0,936 | 0,295 | 0 | Mature B | HLA-DRB1 |
| HLA-DPA12 | 0 | 1,01892828 | 0,861 | 0,225 | 0 | Mature B | HLA-DPA1 |
| CD742 | 0 | 2,19389977 | 0,989 | 0,367 | 0 | Mature B | CD74 |
| MEF2C2 | 0 | 4,00103799 | 0,669 | 0,061 | 0 | Mature B | MEF2C |
| HLA-DQB12 | 0 | 1,44574433 | 0,807 | 0,2 | 0 | Mature B | HLA-DQB1 |
| RALGPS2 | 0 | 5,04555519 | 0,633 | 0,033 | 0 | Mature B | RALGPS2 |
| IGKC | 0 | 1,79242229 | 0,614 | 0,019 | 0 | Mature B | IGKC |
| CD37 | 0 | 2,53584799 | 0,9 | 0,307 | 0 | Mature B | CD37 |
| LYN3 | 0 | 2,7360131 | 0,767 | 0,178 | 0 | Mature B | LYN |
| CD833 | 0 | 2,8617333 | 0,728 | 0,146 | 0 | Mature B | CD83 |
| ADAM282 | 0 | 4,93570629 | 0,603 | 0,031 | 0 | Mature B | ADAM28 |
| AC120193.11 | 0 | 6,20969745 | 0,585 | 0,014 | 0 | Mature B | AC120193.1 |
| TNFRSF13C | 0 | 7,1653106 | 0,569 | 0,005 | 0 | Mature B | TNFRSF13C |
| LINC00926 | 0 | 9,46355815 | 0,561 | 0,001 | 0 | Mature B | LINC00926 |
| LY91 | 0 | 5,443304 | 0,576 | 0,028 | 0 | Mature B | LY9 |
| PLEKHG12 | 0 | 4,45039201 | 0,601 | 0,056 | 0 | Mature B | PLEKHG1 |
| IGHM | 0 | 3,90008226 | 0,534 | 0,002 | 0 | Mature B | IGHM |
| PRDM21 | 0 | 2,48230062 | 0,702 | 0,214 | 0 | Mature B | PRDM2 |
| MGAT52 | 0 | 2,69810315 | 0,616 | 0,174 | 0 | Mature B | MGAT5 |
| BLK1 | 0 | 6,75554393 | 0,442 | 0,005 | 0 | Mature B | BLK |
| SWAP702 | 0 | 4,38822596 | 0,471 | 0,034 | 0 | Mature B | SWAP70 |
| CHPT12 | 0 | 2,98853027 | 0,491 | 0,086 | 0 | Mature B | CHPT1 |
| TCF42 | 0 | 3,51358298 | 0,456 | 0,053 | 0 | Mature B | TCF4 |
| ST6GAL11 | 0 | 3,44750802 | 0,457 | 0,061 | 0 | Mature B | ST6GAL1 |
| TRIO3 | 0 | 2,87252113 | 0,468 | 0,077 | 0 | Mature B | TRIO |
| BCL11A2 | 0 | 5,45644542 | 0,401 | 0,012 | 0 | Mature B | BCL11A |
| OSBPL10 | 0 | 6,80894522 | 0,392 | 0,005 | 0 | Mature B | OSBPL10 |
| CDK143 | 0 | 3,86273711 | 0,423 | 0,04 | 0 | Mature B | CDK14 |
| FCRL1 | 0 | 9,41309308 | 0,364 | 0,001 | 0 | Mature B | FCRL1 |
| PIKFYVE | 0 | 3,78207399 | 0,405 | 0,059 | 0 | Mature B | PIKFYVE |
| MARCH12 | 0 | 4,20039713 | 0,372 | 0,028 | 0 | Mature B | MARCH1 |
| CCSER11 | 0 | 3,34659408 | 0,398 | 0,055 | 0 | Mature B | CCSER1 |
| IGHD | 0 | 10,9360459 | 0,342 | 0 | 0 | Mature B | IGHD |
| RUBCNL1 | 0 | 5,86230184 | 0,349 | 0,007 | 0 | Mature B | RUBCNL |
| INPP5A2 | 0 | 3,04687718 | 0,401 | 0,07 | 0 | Mature B | INPP5A |
| SNED12 | 0 | 4,1291607 | 0,362 | 0,031 | 0 | Mature B | SNED1 |
| LARGE12 | 0 | 4,34059663 | 0,354 | 0,023 | 0 | Mature B | LARGE1 |
| COBLL1 | 0 | 5,42906091 | 0,333 | 0,007 | 0 | Mature B | COBLL1 |
| ST6GALNAC3 | 0 | 2,97028438 | 0,365 | 0,058 | 0 | Mature B | ST6GALNAC3 |
| SEL1L3 | 0 | 2,59103956 | 0,344 | 0,056 | 0 | Mature B | SEL1L3 |
| KHDRBS21 | 0 | 6,07038008 | 0,278 | 0,005 | 0 | Mature B | KHDRBS2 |
| LINC02397 | 0 | 9,22757666 | 0,267 | 0 | 0 | Mature B | LINC02397 |
| WDFY41 | 0 | 3,90242526 | 0,278 | 0,013 | 0 | Mature B | WDFY4 |
| JADE31 | 0 | 6,27538655 | 0,257 | 0,004 | 0 | Mature B | JADE3 |
| NIBAN3 | 0 | 6,05373836 | 0,256 | 0,004 | 0 | Mature B | NIBAN3 |
| IGLC2 | 0 | 2,6049177 | 0,253 | 0,001 | 0 | Mature B | IGLC2 |
| GRK32 | 0 | 2,99369663 | 0,28 | 0,036 | 0 | Mature B | GRK3 |
| C12orf421 | 0 | 5,26069489 | 0,25 | 0,009 | 0 | Mature B | C12orf42 |
| PAX5 | 0 | 10,5242541 | 0,24 | 0 | 0 | Mature B | PAX5 |
| COL19A1 | 0 | 9,81957944 | 0,218 | 0 | 0 | Mature B | COL19A1 |
| STAP12 | 0 | 5,5134067 | 0,218 | 0,005 | 0 | Mature B | STAP1 |
| ANKRD33B2 | 0 | 3,60990519 | 0,233 | 0,025 | 0 | Mature B | ANKRD33B |
| USP6NL1 | 0 | 3,63085289 | 0,227 | 0,02 | 0 | Mature B | USP6NL |
| CARMIL1 | 0 | 3,99053747 | 0,211 | 0,018 | 0 | Mature B | CARMIL1 |
| P2RX51 | 0 | 3,92977109 | 0,209 | 0,017 | 0 | Mature B | P2RX5 |
| LINC01781 | 0 | 10,3704487 | 0,19 | 0 | 0 | Mature B | LINC01781 |
| STEAP1B | 0 | 6,43093673 | 0,195 | 0,005 | 0 | Mature B | STEAP1B |
| SPIB1 | 0 | 4,2186367 | 0,2 | 0,01 | 0 | Mature B | SPIB |
| CD24 | 0 | 9,00787867 | 0,189 | 0 | 0 | Mature B | CD24 |
| TNFRSF13B1 | 0 | 5,89324242 | 0,185 | 0,003 | 0 | Mature B | TNFRSF13B |
| RPS112 | 0 | 1,3518163 | 0,959 | 0,785 | 0 | Mature B | RPS11 |
| RHEX | 0 | 3,6093265 | 0,167 | 0,008 | 0 | Mature B | RHEX |
| JCHAIN | 0 | 1,34765196 | 0,162 | 0,009 | 0 | Mature B | JCHAIN |
| PKIG | 0 | 4,04654029 | 0,157 | 0,01 | 0 | Mature B | PKIG |

| row.names | p_val | avg_log2FC | pct.1 | pct.2 | p_val_adj | cluster | gene |
|---|---|---|---|---|---|---|---|
| LST12 | 0 | 5,48302432 | 0,99 | 0,087 | 0 | NCMs | LST1 |
| C5AR12 | 0 | 4,33508044 | 0,973 | 0,077 | 0 | NCMs | C5AR1 |
| SERPINA11 | 0 | 5,76285301 | 0,922 | 0,029 | 0 | NCMs | SERPINA1 |
| RNF144B2 | 0 | 4,64359686 | 0,984 | 0,101 | 0 | NCMs | RNF144B |
| AIF12 | 0 | 4,48039165 | 0,975 | 0,102 | 0 | NCMs | AIF1 |
| ASAH12 | 0 | 4,36414441 | 0,963 | 0,092 | 0 | NCMs | ASAH1 |
| IRAK33 | 0 | 3,98191543 | 0,978 | 0,121 | 0 | NCMs | IRAK3 |
| MS4A72 | 0 | 4,57718104 | 0,909 | 0,058 | 0 | NCMs | MS4A7 |
| IFI302 | 0 | 4,1356924 | 0,933 | 0,091 | 0 | NCMs | IFI30 |
| PAPSS22 | 0 | 5,02727969 | 0,87 | 0,046 | 0 | NCMs | PAPSS2 |
| FAM49A3 | 0 | 3,6542525 | 0,947 | 0,126 | 0 | NCMs | FAM49A |
| KYNU2 | 0 | 3,36770374 | 0,911 | 0,094 | 0 | NCMs | KYNU |
| PSAP2 | 0 | 3,41155012 | 0,987 | 0,173 | 0 | NCMs | PSAP |
| LILRB22 | 0 | 5,74467413 | 0,836 | 0,023 | 0 | NCMs | LILRB2 |
| LILRA51 | 0 | 6,55211651 | 0,823 | 0,013 | 0 | NCMs | LILRA5 |
| CST32 | 0 | 0,89883351 | 0,949 | 0,145 | 0 | NCMs | CST3 |
| CTSS2 | 0 | 3,70567549 | 0,99 | 0,191 | 0 | NCMs | CTSS |
| WARS3 | 0 | 5,50455267 | 0,826 | 0,028 | 0 | NCMs | WARS |
| PLAUR2 | 0 | 3,35012867 | 0,93 | 0,133 | 0 | NCMs | PLAUR |
| LYN4 | 0 | 3,63521272 | 0,973 | 0,176 | 0 | NCMs | LYN |
| FCER1G5 | 0 | 3,50248951 | 0,983 | 0,202 | 0 | NCMs | FCER1G |
| BCL2A13 | 0 | 3,73462991 | 0,878 | 0,102 | 0 | NCMs | BCL2A1 |
| CLEC7A2 | 0 | 4,11353495 | 0,832 | 0,062 | 0 | NCMs | CLEC7A |
| SMIM251 | 0 | 6,81646029 | 0,775 | 0,01 | 0 | NCMs | SMIM25 |
| PILRA2 | 0 | 5,16479794 | 0,791 | 0,029 | 0 | NCMs | PILRA |
| IFITM32 | 0 | 4,16952648 | 0,837 | 0,081 | 0 | NCMs | IFITM3 |
| BASP13 | 0 | 3,97944921 | 0,809 | 0,063 | 0 | NCMs | BASP1 |
| TIMP12 | 0 | 4,12736655 | 0,927 | 0,181 | 0 | NCMs | TIMP1 |
| LIMS12 | 0 | 3,51821882 | 0,896 | 0,156 | 0 | NCMs | LIMS1 |
| ACSL12 | 0 | 3,43842081 | 0,834 | 0,095 | 0 | NCMs | ACSL1 |
| HLA-DRA3 | 0 | 0,69841857 | 0,96 | 0,222 | 0 | NCMs | HLA-DRA |
| COTL12 | 0 | 3,49123753 | 0,99 | 0,253 | 0 | NCMs | COTL1 |
| SOD22 | 0 | 3,18116835 | 0,948 | 0,212 | 0 | NCMs | SOD2 |
| TCF7L21 | 0 | 4,49535025 | 0,788 | 0,063 | 0 | NCMs | TCF7L2 |
| CD862 | 0 | 4,16280629 | 0,769 | 0,059 | 0 | NCMs | CD86 |
| RNF132 | 0 | 3,92477726 | 0,815 | 0,107 | 0 | NCMs | RNF13 |
| ZFAND52 | 0 | 3,34205941 | 0,933 | 0,227 | 0 | NCMs | ZFAND5 |
| FGD42 | 0 | 3,33086173 | 0,778 | 0,072 | 0 | NCMs | FGD4 |
| PAG11 | 0 | 3,4471004 | 0,87 | 0,164 | 0 | NCMs | PAG1 |
| TYROBP5 | 0 | 2,62312944 | 0,986 | 0,28 | 0 | NCMs | TYROBP |
| ETS22 | 0 | 3,21231458 | 0,805 | 0,102 | 0 | NCMs | ETS2 |
| FPR12 | 0 | 4,41791758 | 0,744 | 0,041 | 0 | NCMs | FPR1 |
| SLC11A11 | 0 | 3,9993699 | 0,739 | 0,039 | 0 | NCMs | SLC11A1 |
| CMTM63 | 0 | 2,93433911 | 0,87 | 0,17 | 0 | NCMs | CMTM6 |
| CEBPD3 | 0 | 2,36923143 | 0,9 | 0,21 | 0 | NCMs | CEBPD |
| TBC1D82 | 0 | 4,49726042 | 0,731 | 0,05 | 0 | NCMs | TBC1D8 |
| NAMPT2 | 0 | 3,337937 | 0,993 | 0,313 | 0 | NCMs | NAMPT |
| STXBP21 | 0 | 4,64180051 | 0,719 | 0,043 | 0 | NCMs | STXBP2 |
| GAB22 | 0 | 3,0226719 | 0,806 | 0,132 | 0 | NCMs | GAB2 |
| MAFB2 | 0 | 3,95499595 | 0,727 | 0,053 | 0 | NCMs | MAFB |
| IL1B2 | 0 | 3,85968545 | 0,732 | 0,071 | 0 | NCMs | IL1B |
| CYBB2 | 0 | 3,90168338 | 0,71 | 0,051 | 0 | NCMs | CYBB |
| TBXAS13 | 0 | 3,8569387 | 0,722 | 0,064 | 0 | NCMs | TBXAS1 |
| G0S22 | 0 | 3,88771069 | 0,71 | 0,056 | 0 | NCMs | G0S2 |
| FCGR3A4 | 0 | 2,83515557 | 0,787 | 0,135 | 0 | NCMs | FCGR3A |
| DMXL21 | 0 | 4,29119124 | 0,688 | 0,038 | 0 | NCMs | DMXL2 |
| DOCK41 | 0 | 3,15702633 | 0,72 | 0,073 | 0 | NCMs | DOCK4 |
| FCN12 | 0 | 3,41722707 | 0,705 | 0,059 | 0 | NCMs | FCN1 |
| DENND5A2 | 0 | 3,44074891 | 0,718 | 0,078 | 0 | NCMs | DENND5A |
| RIPK23 | 0 | 3,07796506 | 0,785 | 0,149 | 0 | NCMs | RIPK2 |
| TYMP2 | 0 | 3,08360001 | 0,733 | 0,098 | 0 | NCMs | TYMP |
| MCTP12 | 0 | 2,9468327 | 0,723 | 0,091 | 0 | NCMs | MCTP1 |
| NFKBIZ3 | 0 | 2,73758052 | 0,818 | 0,187 | 0 | NCMs | NFKBIZ |
| NPC22 | 0 | 2,56036675 | 0,756 | 0,126 | 0 | NCMs | NPC2 |
| FPR21 | 0 | 7,25837602 | 0,636 | 0,006 | 0 | NCMs | FPR2 |
| CHSY12 | 0 | 3,71193464 | 0,7 | 0,07 | 0 | NCMs | CHSY1 |
| TLR22 | 0 | 3,33962532 | 0,699 | 0,07 | 0 | NCMs | TLR2 |
| EPB41L32 | 0 | 3,31749364 | 0,692 | 0,066 | 0 | NCMs | EPB41L3 |
| STX112 | 0 | 2,83883589 | 0,756 | 0,134 | 0 | NCMs | STX11 |
| MTSS15 | 0 | 2,93693074 | 0,765 | 0,151 | 0 | NCMs | MTSS1 |
| HLA-DPA13 | 0 | 0,87306241 | 0,836 | 0,226 | 0 | NCMs | HLA-DPA1 |
| TKT2 | 0 | 3,11976956 | 0,705 | 0,096 | 0 | NCMs | TKT |
| PELI14 | 0 | 2,7301228 | 0,839 | 0,231 | 0 | NCMs | PELI1 |
| RILPL23 | 0 | 2,1423578 | 0,847 | 0,242 | 0 | NCMs | RILPL2 |

| row.names | p_val | avg_log2FC | pct.1 | pct.2 | p_val_ad | cluster | gene |
|---|---|---|---|---|---|---|---|
| SELENOP1 | 0 | 7,35341604 | 0,899 | 0,016 | 0 | PVMs | SELENOP |
| C1QA3 | 0 | 6,50574518 | 0,895 | 0,022 | 0 | PVMs | C1QA |
| RNASE11 | 0 | 7,99630672 | 0,869 | 0,016 | 0 | PVMs | RNASE1 |
| C1QC2 | 0 | 6,9042824 | 0,855 | 0,012 | 0 | PVMs | C1QC |
| CD1632 | 0 | 5,19923083 | 0,885 | 0,052 | 0 | PVMs | CD163 |
| MRC12 | 0 | 6,09144809 | 0,845 | 0,034 | 0 | PVMs | MRC1 |
| C1QB3 | 0 | 6,55646206 | 0,816 | 0,014 | 0 | PVMs | C1QB |
| FOLR21 | 0 | 6,78834549 | 0,799 | 0,013 | 0 | PVMs | FOLR2 |
| CST33 | 0 | 1,89921051 | 0,912 | 0,146 | 0 | PVMs | CST3 |
| STARD132 | 0 | 6,3702871 | 0,782 | 0,023 | 0 | PVMs | STARD13 |
| PDK42 | 0 | 6,03483949 | 0,769 | 0,04 | 0 | PVMs | PDK4 |
| LYVE11 | 0 | 7,57577777 | 0,736 | 0,007 | 0 | PVMs | LYVE1 |
| CTSB3 | 0 | 3,57273362 | 0,831 | 0,105 | 0 | PVMs | CTSB |
| F13A12 | 0 | 6,33368633 | 0,735 | 0,02 | 0 | PVMs | F13A1 |
| FCGRT3 | 0 | 3,99846315 | 0,796 | 0,085 | 0 | PVMs | FCGRT |
| MAFB3 | 0 | 4,34733033 | 0,756 | 0,053 | 0 | PVMs | MAFB |
| KLF43 | 0 | 3,8435975 | 0,798 | 0,097 | 0 | PVMs | KLF4 |
| LRMDA3 | 0 | 3,75906463 | 0,77 | 0,076 | 0 | PVMs | LRMDA |
| RBM473 | 0 | 3,35422255 | 0,774 | 0,081 | 0 | PVMs | RBM47 |
| MCTP13 | 0 | 3,65645642 | 0,784 | 0,091 | 0 | PVMs | MCTP1 |
| CD835 | 0 | 3,21184718 | 0,832 | 0,145 | 0 | PVMs | CD83 |
| FMN12 | 0 | 4,32106745 | 0,782 | 0,097 | 0 | PVMs | FMN1 |
| MS4A6A2 | 0 | 3,42938084 | 0,747 | 0,082 | 0 | PVMs | MS4A6A |
| LGMN1 | 0 | 6,75020859 | 0,681 | 0,016 | 0 | PVMs | LGMN |
| MS4A4A3 | 0 | 6,07684785 | 0,678 | 0,015 | 0 | PVMs | MS4A4A |
| NPC23 | 0 | 3,29062333 | 0,787 | 0,126 | 0 | PVMs | NPC2 |
| HLA-DRA4 | 0 | 1,53658577 | 0,883 | 0,223 | 0 | PVMs | HLA-DRA |
| PSAP3 | 0 | 2,65686612 | 0,831 | 0,176 | 0 | PVMs | PSAP |
| CD683 | 0 | 4,02747582 | 0,705 | 0,055 | 0 | PVMs | CD68 |
| MS4A73 | 0 | 3,92456454 | 0,706 | 0,061 | 0 | PVMs | MS4A7 |
| PID13 | 0 | 3,58938003 | 0,715 | 0,072 | 0 | PVMs | PID1 |
| MAN1A14 | 0 | 3,61307727 | 0,809 | 0,166 | 0 | PVMs | MAN1A1 |
| CPM2 | 0 | 4,72960086 | 0,68 | 0,039 | 0 | PVMs | CPM |
| ATF64 | 0 | 3,43121642 | 0,797 | 0,162 | 0 | PVMs | ATF6 |
| CXCL83 | 0 | 3,10924337 | 0,731 | 0,096 | 0 | PVMs | CXCL8 |
| EGR13 | 0 | 4,80399226 | 0,665 | 0,057 | 0 | PVMs | EGR1 |
| ABCA13 | 0 | 3,98725751 | 0,659 | 0,051 | 0 | PVMs | ABCA1 |
| EMP12 | 0 | 4,90521726 | 0,665 | 0,061 | 0 | PVMs | EMP1 |
| C5AR13 | 0 | 2,50266859 | 0,675 | 0,081 | 0 | PVMs | C5AR1 |
| ITSN13 | 0 | 4,86536728 | 0,626 | 0,034 | 0 | PVMs | ITSN1 |
| SGK12 | 0 | 3,17708382 | 0,714 | 0,123 | 0 | PVMs | SGK1 |
| FNIP23 | 0 | 3,44216944 | 0,683 | 0,094 | 0 | PVMs | FNIP2 |
| DAB21 | 0 | 6,59113426 | 0,6 | 0,011 | 0 | PVMs | DAB2 |
| PEAK15 | 0 | 3,37770336 | 0,717 | 0,131 | 0 | PVMs | PEAK1 |
| RASGEF1B4 | 0 | 2,71961296 | 0,781 | 0,196 | 0 | PVMs | RASGEF1B |
| CEBPD4 | 0 | 2,46118897 | 0,791 | 0,212 | 0 | PVMs | CEBPD |
| AIF13 | 0 | 2,382739 | 0,683 | 0,106 | 0 | PVMs | AIF1 |
| CXCL23 | 0 | 3,32362 | 0,648 | 0,075 | 0 | PVMs | CXCL2 |
| FGL23 | 0 | 3,71292123 | 0,719 | 0,148 | 0 | PVMs | FGL2 |
| BLVRB3 | 0 | 3,60662183 | 0,637 | 0,071 | 0 | PVMs | BLVRB |
| SH3BP53 | 0 | 2,99126028 | 0,689 | 0,124 | 0 | PVMs | SH3BP5 |
| IRAK34 | 0 | 2,39846814 | 0,689 | 0,125 | 0 | PVMs | IRAK3 |
| RAB313 | 0 | 3,09161493 | 0,651 | 0,088 | 0 | PVMs | RAB31 |
| HLA-DRB13 | 0 | 1,58707553 | 0,857 | 0,297 | 0 | PVMs | HLA-DRB1 |
| COLEC121 | 0 | 7,40101079 | 0,566 | 0,006 | 0 | PVMs | COLEC12 |
| RHOB3 | 0 | 4,00964275 | 0,625 | 0,066 | 0 | PVMs | RHOB |
| FCGR2A3 | 0 | 3,32107407 | 0,629 | 0,072 | 0 | PVMs | FCGR2A |
| RAB7A3 | 0 | 3,19789136 | 0,798 | 0,241 | 0 | PVMs | RAB7A |
| CD743 | 0 | 2,11154622 | 0,922 | 0,368 | 0 | PVMs | CD74 |
| SRGAP13 | 0 | 3,79813796 | 0,601 | 0,047 | 0 | PVMs | SRGAP1 |
| ETS23 | 0 | 2,98661358 | 0,649 | 0,104 | 0 | PVMs | ETS2 |
| CREB52 | 0 | 4,65026268 | 0,584 | 0,042 | 0 | PVMs | CREB5 |
| AFF42 | 0 | 2,81316293 | 0,727 | 0,189 | 0 | PVMs | AFF4 |
| CD143 | 0 | 4,81962696 | 0,563 | 0,026 | 0 | PVMs | CD14 |
| QKI3 | 0 | 2,3678161 | 0,774 | 0,238 | 0 | PVMs | QKI |
| ZFAND53 | 0 | 2,29523215 | 0,765 | 0,229 | 0 | PVMs | ZFAND5 |
| MFSD14 | 0 | 4,08000205 | 0,58 | 0,044 | 0 | PVMs | MFSD1 |
| CSF1R3 | 0 | 5,07159728 | 0,554 | 0,02 | 0 | PVMs | CSF1R |
| CFD2 | 0 | 1,89985182 | 0,581 | 0,05 | 0 | PVMs | CFD |
| HLA-DPA14 | 0 | 1,37006679 | 0,756 | 0,227 | 0 | PVMs | HLA-DPA1 |
| STAB11 | 0 | 7,02930982 | 0,53 | 0,006 | 0 | PVMs | STAB1 |
| RAPH13 | 0 | 3,96155458 | 0,573 | 0,054 | 0 | PVMs | RAPH1 |
| MSR13 | 0 | 4,49553354 | 0,554 | 0,036 | 0 | PVMs | MSR1 |
| LILRB51 | 0 | 8,45717811 | 0,517 | 0,002 | 0 | PVMs | LILRB5 |

| row.names | p_val | avg_log2FC | pct.1 | pct.2 | p_val_adj | cluster | gene |
|---|---|---|---|---|---|---|---|
| NDFIP13 | 0 | 2,33091292 | 0,85 | 0,307 | 0 | Naïve CD4+ T | NDFIP1 |
| LEF12 | 0 | 3,55575381 | 0,557 | 0,057 | 0 | Naïve CD4+ T | LEF1 |
| SELL4 | 0 | 3,43304926 | 0,524 | 0,06 | 0 | Naïve CD4+ T | SELL |
| PRKCQ-AS14 | 0 | 2,93541771 | 0,508 | 0,081 | 0 | Naïve CD4+ T | PRKCQ-AS1 |
| FHIT3 | 0 | 3,59884623 | 0,482 | 0,056 | 0 | Naïve CD4+ T | FHIT |
| TAF4B3 | 0 | 3,53235799 | 0,464 | 0,061 | 0 | Naïve CD4+ T | TAF4B |
| NELL22 | 0 | 3,23491713 | 0,463 | 0,066 | 0 | Naïve CD4+ T | NELL2 |
| CCR73 | 0 | 2,44129346 | 0,436 | 0,072 | 0 | Naïve CD4+ T | CCR7 |
| TSHZ22 | 0 | 3,66225514 | 0,377 | 0,034 | 0 | Naïve CD4+ T | TSHZ2 |
| SCML15 | 0 | 3,13739305 | 0,346 | 0,045 | 0 | Naïve CD4+ T | SCML1 |
| ADTRP4 | 0 | 5,36802947 | 0,177 | 0,005 | 0 | Naïve CD4+ T | ADTRP |
| RPL52 | 0 | 1,39245314 | 0,979 | 0,847 | 0 | Naïve CD4+ T | RPL5 |
| RPS133 | 0 | 1,44204667 | 0,986 | 0,879 | 0 | Naïve CD4+ T | RPS13 |
| RPL93 | 0 | 1,34690874 | 0,985 | 0,894 | 0 | Naïve CD4+ T | RPL9 |
| LEF1-AS1 | 0 | 5,83933519 | 0,072 | 0,001 | 0 | Naïve CD4+ T | LEF1-AS1 |
| RPS62 | 0 | 1,50804751 | 0,982 | 0,912 | 0 | Naïve CD4+ T | RPS6 |
| RPS82 | 0 | 1,44626988 | 0,986 | 0,924 | 0 | Naïve CD4+ T | RPS8 |
| RPL212 | 0 | 1,21521164 | 0,986 | 0,936 | 0 | Naïve CD4+ T | RPL21 |
| RPS3A2 | 0 | 1,32621367 | 0,986 | 0,941 | 0 | Naïve CD4+ T | RPS3A |
| RPL323 | 0 | 1,31727499 | 0,986 | 0,948 | 0 | Naïve CD4+ T | RPL32 |
| RPS123 | 0 | 1,30480354 | 0,987 | 0,957 | 0 | Naïve CD4+ T | RPS12 |
| RPL133 | 0 | 1,31957959 | 0,987 | 0,965 | 0 | Naïve CD4+ T | RPL13 |
| RPL303 | 1E-302 | 1,16555426 | 0,987 | 0,95 | 4E-298 | Naïve CD4+ T | RPL30 |
| RPS142 | 5E-301 | 1,14432928 | 0,986 | 0,947 | 1E-296 | Naïve CD4+ T | RPS14 |
| RPS232 | 3E-300 | 1,22939657 | 0,987 | 0,937 | 8E-296 | Naïve CD4+ T | RPS23 |
| RPL113 | 4E-294 | 1,13169925 | 0,986 | 0,946 | 9E-290 | Naïve CD4+ T | RPL11 |
| RPL222 | 4E-292 | 1,34101141 | 0,977 | 0,779 | 9E-288 | Naïve CD4+ T | RPL22 |
| RPL372 | 2E-283 | 1,0180567 | 0,986 | 0,95 | 4E-279 | Naïve CD4+ T | RPL37 |
| RPS53 | 2E-274 | 1,33180666 | 0,978 | 0,773 | 5E-270 | Naïve CD4+ T | RPS5 |
| RPL182 | 2E-274 | 1,17616303 | 0,986 | 0,896 | 6E-270 | Naïve CD4+ T | RPL18 |
| RPS253 | 2E-273 | 1,14840219 | 0,986 | 0,892 | 5E-269 | Naïve CD4+ T | RPS25 |
| EEF1B22 | 3E-272 | 1,63140118 | 0,952 | 0,653 | 8E-268 | Naïve CD4+ T | EEF1B2 |
| LDHB2 | 3E-271 | 2,05601684 | 0,661 | 0,181 | 7E-267 | Naïve CD4+ T | LDHB |
| RPS15A2 | 8E-270 | 0,97872203 | 0,986 | 0,947 | 2E-265 | Naïve CD4+ T | RPS15A |
| FAAH22 | 1E-269 | 2,95966087 | 0,384 | 0,066 | 3E-265 | Naïve CD4+ T | FAAH2 |
| RPL343 | 5E-269 | 1,01474203 | 0,986 | 0,955 | 1E-264 | Naïve CD4+ T | RPL34 |
| RETREG15 | 2E-268 | 2,97591961 | 0,323 | 0,047 | 4E-264 | Naïve CD4+ T | RETREG1 |
| RPL193 | 2E-268 | 1,11771278 | 0,986 | 0,934 | 4E-264 | Naïve CD4+ T | RPL19 |
| RPS27A3 | 3E-266 | 1,00645143 | 0,987 | 0,951 | 7E-262 | Naïve CD4+ T | RPS27A |
| RPL73 | 5E-265 | 1,43713647 | 0,949 | 0,684 | 1E-260 | Naïve CD4+ T | RPL7 |
| RPS183 | 5E-263 | 1,11253308 | 0,986 | 0,951 | 1E-258 | Naïve CD4+ T | RPS18 |
| RPS4X3 | 8E-263 | 1,12654465 | 0,987 | 0,928 | 2E-258 | Naïve CD4+ T | RPS4X |
| TPT11 | 8E-263 | 0,99168562 | 0,99 | 0,965 | 2E-258 | Naïve CD4+ T | TPT1 |
| RPL10A3 | 9E-262 | 1,26246557 | 0,979 | 0,82 | 2E-257 | Naïve CD4+ T | RPL10A |
| RPL234 | 6E-261 | 1,53480102 | 0,946 | 0,674 | 1E-256 | Naïve CD4+ T | RPL23 |
| RPL310 | 6E-256 | 1,18616103 | 0,983 | 0,903 | 1E-251 | Naïve CD4+ T | RPL3 |
| KLF23 | 3E-251 | 1,55444259 | 0,931 | 0,44 | 7E-247 | Naïve CD4+ T | KLF2 |
| RPL35A3 | 7E-248 | 1,05019436 | 0,987 | 0,901 | 2E-243 | Naïve CD4+ T | RPL35A |
| AP000787.13 | 1E-243 | 2,76931381 | 0,361 | 0,064 | 3E-239 | Naïve CD4+ T | AP000787.1 |
| LINC018914 | 8E-240 | 3,67923115 | 0,156 | 0,013 | 2E-235 | Naïve CD4+ T | LINC01891 |
| RPS32 | 1E-239 | 1,0655131 | 0,985 | 0,915 | 3E-235 | Naïve CD4+ T | RPS3 |
| LTB5 | 2E-239 | 1,5907005 | 0,676 | 0,198 | 4E-235 | Naïve CD4+ T | LTB |
| RPS283 | 3E-238 | 0,96331499 | 0,988 | 0,95 | 9E-234 | Naïve CD4+ T | RPS28 |
| RPL393 | 3E-238 | 0,94077857 | 0,986 | 0,951 | 9E-234 | Naïve CD4+ T | RPL39 |
| RPLP23 | 6E-237 | 0,94079973 | 0,987 | 0,926 | 1E-232 | Naïve CD4+ T | RPLP2 |
| RPL18A3 | 1E-236 | 1,07942403 | 0,986 | 0,919 | 3E-232 | Naïve CD4+ T | RPL18A |
| SARAF2 | 2E-235 | 1,30256229 | 0,979 | 0,702 | 5E-231 | Naïve CD4+ T | SARAF |
| SESN33 | 2E-235 | 2,41554413 | 0,526 | 0,133 | 6E-231 | Naïve CD4+ T | SESN3 |
| OXNAD12 | 2E-234 | 2,1110621 | 0,594 | 0,167 | 6E-230 | Naïve CD4+ T | OXNAD1 |
| RPL102 | 3E-232 | 0,87768491 | 0,988 | 0,969 | 7E-228 | Naïve CD4+ T | RPL10 |
| RPL292 | 4E-231 | 1,08688274 | 0,988 | 0,88 | 1E-226 | Naïve CD4+ T | RPL29 |
| EEF1A12 | 1E-224 | 0,79589554 | 0,988 | 0,974 | 4E-220 | Naïve CD4+ T | EEF1A1 |
| RPL27A2 | 6E-224 | 1,20445163 | 0,972 | 0,827 | 2E-219 | Naïve CD4+ T | RPL27A |
| GAS54 | 1E-221 | 1,86468333 | 0,751 | 0,295 | 3E-217 | Naïve CD4+ T | GAS5 |
| AL109930.11 | 8E-220 | 4,39915198 | 0,09 | 0,005 | 2E-215 | Naïve CD4+ T | AL109930.1 |
| RPS153 | 5E-218 | 0,91872878 | 0,986 | 0,93 | 1E-213 | Naïve CD4+ T | RPS15 |
| RPL363 | 7E-218 | 1,03315253 | 0,987 | 0,883 | 2E-213 | Naïve CD4+ T | RPL36 |
| RPS203 | 6E-216 | 1,33435016 | 0,932 | 0,707 | 2E-211 | Naïve CD4+ T | RPS20 |
| RPS272 | 1E-211 | 0,78014459 | 0,986 | 0,97 | 3E-207 | Naïve CD4+ T | RPS27 |
| EEF1G2 | 1E-210 | 1,46024564 | 0,893 | 0,517 | 2E-206 | Naïve CD4+ T | EEF1G |
| RPL312 | 5E-208 | 1,3367658 | 0,916 | 0,605 | 1E-203 | Naïve CD4+ T | RPL31 |
| RPS210 | 3E-204 | 0,92277782 | 0,985 | 0,932 | 6E-200 | Naïve CD4+ T | RPS2 |
| RPL13A2 | 7E-204 | 1,05683608 | 0,973 | 0,859 | 2E-199 | Naïve CD4+ T | RPL13A |
| RPS213 | 1E-203 | 0,89558929 | 0,988 | 0,921 | 3E-199 | Naïve CD4+ T | RPS21 |

| row.names | p_val | avg_log2FC | pct.1 | pct.2 | p_val_ad | cluster | gene |
|---|---|---|---|---|---|---|---|
| TNFRSF182 | 0 | 3,85719799 | 0,715 | 0,08 | 0 | ILCs | TNFRSF18 |
| AL136456.13 | 0 | 3,83459472 | 0,601 | 0,07 | 0 | ILCs | AL136456.1 |
| KIT3 | 0 | 7,08152179 | 0,526 | 0,006 | 0 | ILCs | KIT |
| TNFRSF43 | 0 | 3,60368466 | 0,481 | 0,049 | 0 | ILCs | TNFRSF4 |
| XCL12 | 0 | 3,17732877 | 0,497 | 0,074 | 0 | ILCs | XCL1 |
| AFF33 | 0 | 3,15790401 | 0,449 | 0,066 | 0 | ILCs | AFF3 |
| ATP8B45 | 0 | 3,52445664 | 0,391 | 0,048 | 0 | ILCs | ATP8B4 |
| FXYD72 | 0 | 3,89707967 | 0,338 | 0,031 | 0 | ILCs | FXYD7 |
| SPINK22 | 0 | 5,55512549 | 0,294 | 0,009 | 0 | ILCs | SPINK2 |
| AL031599.12 | 0 | 6,58284188 | 0,256 | 0,004 | 0 | ILCs | AL031599.1 |
| PCDH91 | 0 | 6,14537902 | 0,228 | 0,005 | 0 | ILCs | PCDH9 |
| GNG43 | 0 | 4,07070067 | 0,237 | 0,019 | 0 | ILCs | GNG4 |
| RBMS31 | 0 | 3,78545599 | 0,213 | 0,012 | 0 | ILCs | RBMS3 |
| MICU33 | 0 | 4,7694287 | 0,206 | 0,009 | 0 | ILCs | MICU3 |
| TNFSF112 | 0 | 6,30142895 | 0,194 | 0,003 | 0 | ILCs | TNFSF11 |
| PPP1R9A3 | 0 | 4,43464511 | 0,168 | 0,008 | 0 | ILCs | PPP1R9A |
| PRKG11 | 0 | 5,09393074 | 0,166 | 0,007 | 0 | ILCs | PRKG1 |
| SCX2 | 0 | 5,53640707 | 0,163 | 0,008 | 0 | ILCs | SCX |
| COL24A11 | 0 | 5,79002305 | 0,15 | 0,003 | 0 | ILCs | COL24A1 |
| GNAI12 | 0 | 5,81992454 | 0,137 | 0,003 | 0 | ILCs | GNAI1 |
| DLL11 | 0 | 5,54509803 | 0,128 | 0,003 | 0 | ILCs | DLL1 |
| SCN1B3 | 0 | 4,82326077 | 0,122 | 0,005 | 0 | ILCs | SCN1B |
| TNFSF41 | 0 | 5,54718536 | 0,112 | 0,003 | 0 | ILCs | TNFSF4 |
| MSRB31 | 0 | 5,5181697 | 0,1 | 0,003 | 0 | ILCs | MSRB3 |
| SEMA3A1 | 0 | 5,5398013 | 0,099 | 0,003 | 0 | ILCs | SEMA3A |
| FSTL41 | 0 | 5,82749776 | 0,088 | 0,001 | 0 | ILCs | FSTL4 |
| ABCB5 | 0 | 6,79273234 | 0,05 | 0 | 0 | ILCs | ABCB5 |
| TMPRSS11E | 0 | 6,80262309 | 0,051 | 0,001 | 0 | ILCs | TMPRSS11E |
| LINC02726 | 0 | 7,56446335 | 0,035 | 0 | 0 | ILCs | LINC02726 |
| AL157895.1 | 0 | 9,67994057 | 0,032 | 0 | 0 | ILCs | AL157895.1 |
| FGF20 | 0 | 10,8863914 | 0,028 | 0 | 0 | ILCs | FGF20 |
| KLRB15 | 2E-302 | 2,27804434 | 0,907 | 0,35 | 6E-298 | ILCs | KLRB1 |
| RASSF83 | 1E-292 | 3,41848405 | 0,29 | 0,032 | 4E-288 | ILCs | RASSF8 |
| FAM189A2 | 7E-277 | 6,85991924 | 0,037 | 0 | 2E-272 | ILCs | FAM189A2 |
| ZNF7042 | 3E-267 | 3,253719 | 0,274 | 0,031 | 7E-263 | ILCs | ZNF704 |
| IL1RL12 | 7E-260 | 5,47013597 | 0,084 | 0,003 | 2E-255 | ILCs | IL1RL1 |
| GAP43 | 4E-250 | 8,88639145 | 0,019 | 0 | 1E-245 | ILCs | GAP43 |
| IL4I13 | 9E-250 | 3,76839424 | 0,215 | 0,021 | 2E-245 | ILCs | IL4I1 |
| PLCB41 | 5E-241 | 5,22886032 | 0,054 | 0,001 | 1E-236 | ILCs | PLCB4 |
| CYSLTR14 | 1E-239 | 3,14056329 | 0,257 | 0,03 | 4E-235 | ILCs | CYSLTR1 |
| BCAS11 | 6E-239 | 5,44628193 | 0,056 | 0,001 | 2E-234 | ILCs | BCAS1 |
| PLA2G4A4 | 7E-237 | 4,42695983 | 0,138 | 0,009 | 2E-232 | ILCs | PLA2G4A |
| ITM2C | 4E-233 | 2,97950085 | 0,246 | 0,028 | 1E-228 | ILCs | ITM2C |
| LINC01229 | 6E-232 | 7,49046277 | 0,022 | 0 | 1E-227 | ILCs | LINC01229 |
| SSBP25 | 3E-231 | 2,55753829 | 0,659 | 0,199 | 7E-227 | ILCs | SSBP2 |
| ATRNL11 | 1E-225 | 6,8678555 | 0,041 | 0,001 | 3E-221 | ILCs | ATRNL1 |
| PDE7B6 | 2E-224 | 3,69967029 | 0,224 | 0,025 | 4E-220 | ILCs | PDE7B |
| HOXA55 | 3E-220 | 5,01849498 | 0,078 | 0,003 | 7E-216 | ILCs | HOXA5 |
| TTLL101 | 2E-213 | 5,11380194 | 0,062 | 0,002 | 6E-209 | ILCs | TTLL10 |
| FXYD52 | 6E-210 | 2,02522452 | 0,828 | 0,392 | 2E-205 | ILCs | FXYD5 |
| HS6ST21 | 2E-207 | 5,05478898 | 0,063 | 0,002 | 5E-203 | ILCs | HS6ST2 |
| HPGDS1 | 1E-201 | 5,00334759 | 0,069 | 0,003 | 3E-197 | ILCs | HPGDS |
| KIAA13243 | 1E-201 | 3,8947271 | 0,172 | 0,017 | 3E-197 | ILCs | KIAA1324 |
| LINC002994 | 2E-199 | 2,94325975 | 0,346 | 0,063 | 4E-195 | ILCs | LINC00299 |
| IL1R14 | 4E-199 | 2,59399914 | 0,29 | 0,044 | 1E-194 | ILCs | IL1R1 |
| JAG14 | 5E-199 | 3,97469987 | 0,124 | 0,009 | 1E-194 | ILCs | JAG1 |
| AL603783.1 | 5E-196 | 5,15783809 | 0,057 | 0,002 | 1E-191 | ILCs | AL603783.1 |
| AREG6 | 6E-191 | 1,90113233 | 0,828 | 0,35 | 2E-186 | ILCs | AREG |
| SRGAP33 | 4E-189 | 4,15529286 | 0,1 | 0,006 | 1E-184 | ILCs | SRGAP3 |
| ZG16B | 3E-188 | 5,81722942 | 0,038 | 0,001 | 6E-184 | ILCs | ZG16B |
| SPRY14 | 6E-187 | 3,0481093 | 0,249 | 0,035 | 1E-182 | ILCs | SPRY1 |
| SOX42 | 4E-183 | 2,9924043 | 0,221 | 0,029 | 9E-179 | ILCs | SOX4 |
| AGPAT54 | 3E-182 | 2,84129237 | 0,304 | 0,053 | 6E-178 | ILCs | AGPAT5 |
| SNAP91 | 6E-179 | 8,3299981 | 0,016 | 0 | 2E-174 | ILCs | SNAP91 |
| DACH14 | 9E-177 | 4,13776294 | 0,088 | 0,005 | 2E-172 | ILCs | DACH1 |
| STAC1 | 7E-171 | 5,18259272 | 0,046 | 0,001 | 2E-166 | ILCs | STAC |
| STAM3 | 6E-169 | 2,03190198 | 0,449 | 0,111 | 1E-164 | ILCs | STAM |
| GPC61 | 4E-168 | 4,91572494 | 0,059 | 0,002 | 9E-164 | ILCs | GPC6 |
| TNFRSF11A3 | 3E-158 | 3,70532589 | 0,112 | 0,009 | 8E-154 | ILCs | TNFRSF11A |
| SORCS1 | 4E-156 | 6,1445601 | 0,046 | 0,001 | 1E-151 | ILCs | SORCS1 |
| COL9A23 | 7E-151 | 4,17469261 | 0,076 | 0,004 | 2E-146 | ILCs | COL9A2 |
| IL2RA4 | 1E-147 | 2,61432798 | 0,257 | 0,046 | 3E-143 | ILCs | IL2RA |
| RUNX24 | 2E-144 | 2,2439351 | 0,416 | 0,11 | 4E-140 | ILCs | RUNX2 |
| TOX22 | 6E-144 | 3,16223046 | 0,165 | 0,02 | 1E-139 | ILCs | TOX2 |

| row.names | p_val | avg_log2FC | pct.1 | pct.2 | p_val_adj | cluster | gene |
|---|---|---|---|---|---|---|---|
| STMN11 | 0 | 5,95912861 | 0,969 | 0,075 | 0 | Proliferating | STMN1 |
| TUBB2 | 0 | 4,86473922 | 0,815 | 0,077 | 0 | Proliferating | TUBB |
| HMGN22 | 0 | 3,88755864 | 0,909 | 0,191 | 0 | Proliferating | HMGN2 |
| TYMS1 | 0 | 7,84761271 | 0,679 | 0,005 | 0 | Proliferating | TYMS |
| PCLAF1 | 0 | 8,2023989 | 0,668 | 0,003 | 0 | Proliferating | PCLAF |
| DEK | 0 | 2,79569279 | 0,875 | 0,226 | 0 | Proliferating | DEK |
| HMGB21 | 0 | 3,22927621 | 0,918 | 0,275 | 0 | Proliferating | HMGB2 |
| CENPF2 | 0 | 5,96477244 | 0,65 | 0,017 | 0 | Proliferating | CENPF |
| DUT1 | 0 | 4,48020912 | 0,686 | 0,064 | 0 | Proliferating | DUT |
| MKI67 | 0 | 8,71813859 | 0,599 | 0,002 | 0 | Proliferating | MKI67 |
| NUSAP11 | 0 | 5,12560856 | 0,621 | 0,026 | 0 | Proliferating | NUSAP1 |
| CKS1B | 0 | 5,44081176 | 0,595 | 0,019 | 0 | Proliferating | CKS1B |
| TOP2A1 | 0 | 6,6371089 | 0,577 | 0,009 | 0 | Proliferating | TOP2A |
| SMC42 | 0 | 3,13508746 | 0,646 | 0,086 | 0 | Proliferating | SMC4 |
| NUCB23 | 0 | 3,8430004 | 0,615 | 0,06 | 0 | Proliferating | NUCB2 |
| RRM21 | 0 | 7,97935555 | 0,55 | 0,003 | 0 | Proliferating | RRM2 |
| ASPM | 0 | 8,22700375 | 0,492 | 0,002 | 0 | Proliferating | ASPM |
| PCNA1 | 0 | 4,24860676 | 0,503 | 0,039 | 0 | Proliferating | PCNA |
| RANBP11 | 0 | 3,30950178 | 0,53 | 0,067 | 0 | Proliferating | RANBP1 |
| ATAD2 | 0 | 3,94083921 | 0,499 | 0,045 | 0 | Proliferating | ATAD2 |
| SMC21 | 0 | 5,13223138 | 0,445 | 0,015 | 0 | Proliferating | SMC2 |
| KNL11 | 0 | 6,32252899 | 0,434 | 0,006 | 0 | Proliferating | KNL1 |
| HIST1H1C1 | 0 | 3,61275366 | 0,47 | 0,057 | 0 | Proliferating | HIST1H1C |
| CLSPN | 0 | 5,69300255 | 0,403 | 0,01 | 0 | Proliferating | CLSPN |
| MCM71 | 0 | 4,28963791 | 0,408 | 0,024 | 0 | Proliferating | MCM7 |
| DIAPH3 | 0 | 8,14808377 | 0,372 | 0,001 | 0 | Proliferating | DIAPH3 |
| HELLS2 | 0 | 4,76711414 | 0,374 | 0,016 | 0 | Proliferating | HELLS |
| NUDT13 | 0 | 3,69210491 | 0,385 | 0,031 | 0 | Proliferating | NUDT1 |
| DTL1 | 0 | 6,79793273 | 0,336 | 0,003 | 0 | Proliferating | DTL |
| BIRC5 | 0 | 9,55672215 | 0,332 | 0 | 0 | Proliferating | BIRC5 |
| CDK12 | 0 | 5,77613414 | 0,332 | 0,006 | 0 | Proliferating | CDK1 |
| CENPU2 | 0 | 5,16395017 | 0,334 | 0,009 | 0 | Proliferating | CENPU |
| TK1 | 0 | 8,04559482 | 0,325 | 0,001 | 0 | Proliferating | TK1 |
| MAD2L11 | 0 | 5,71159389 | 0,327 | 0,006 | 0 | Proliferating | MAD2L1 |
| CENPM3 | 0 | 4,86550657 | 0,305 | 0,011 | 0 | Proliferating | CENPM |
| PRC1 | 0 | 4,80236568 | 0,301 | 0,011 | 0 | Proliferating | PRC1 |
| NCAPG23 | 0 | 4,72868698 | 0,292 | 0,011 | 0 | Proliferating | NCAPG2 |
| FANCI1 | 0 | 4,90836272 | 0,29 | 0,01 | 0 | Proliferating | FANCI |
| TPX2 | 0 | 6,93600548 | 0,281 | 0,003 | 0 | Proliferating | TPX2 |
| UBE2C | 0 | 9,48072949 | 0,276 | 0 | 0 | Proliferating | UBE2C |
| NUF2 | 0 | 6,32823201 | 0,265 | 0,003 | 0 | Proliferating | NUF2 |
| KIF11 | 0 | 5,87894495 | 0,261 | 0,005 | 0 | Proliferating | KIF11 |
| TMEM106C2 | 0 | 4,23487893 | 0,261 | 0,015 | 0 | Proliferating | TMEM106C |
| CENPW | 0 | 6,02525693 | 0,243 | 0,004 | 0 | Proliferating | CENPW |
| CENPE2 | 0 | 4,09834153 | 0,249 | 0,014 | 0 | Proliferating | CENPE |
| HIST1H1B | 0 | 7,41899503 | 0,232 | 0,002 | 0 | Proliferating | HIST1H1B |
| ORC6 | 0 | 5,60996128 | 0,229 | 0,005 | 0 | Proliferating | ORC6 |
| ZWINT | 0 | 7,74621856 | 0,218 | 0,001 | 0 | Proliferating | ZWINT |
| CDT1 | 0 | 8,06348201 | 0,218 | 0,001 | 0 | Proliferating | CDT1 |
| CENPH | 0 | 4,77743095 | 0,223 | 0,008 | 0 | Proliferating | CENPH |
| HIST1H3B | 0 | 7,83888341 | 0,214 | 0,001 | 0 | Proliferating | HIST1H3B |
| BRIP11 | 0 | 4,87118552 | 0,22 | 0,008 | 0 | Proliferating | BRIP1 |
| RAD51AP1 | 0 | 6,60902416 | 0,207 | 0,002 | 0 | Proliferating | RAD51AP1 |
| DHFR | 0 | 4,82685644 | 0,212 | 0,008 | 0 | Proliferating | DHFR |
| GTSE1 | 0 | 7,81614982 | 0,192 | 0,001 | 0 | Proliferating | GTSE1 |
| NCAPG | 0 | 6,78512293 | 0,192 | 0,002 | 0 | Proliferating | NCAPG |
| HMGB31 | 0 | 4,49127269 | 0,2 | 0,01 | 0 | Proliferating | HMGB3 |
| CDCA5 | 0 | 8,0878331 | 0,189 | 0,001 | 0 | Proliferating | CDCA5 |
| FAM111B1 | 0 | 7,39332498 | 0,18 | 0,001 | 0 | Proliferating | FAM111B |
| ASF1B | 0 | 6,3319503 | 0,18 | 0,002 | 0 | Proliferating | ASF1B |
| KIFC1 | 0 | 5,76445234 | 0,18 | 0,003 | 0 | Proliferating | KIFC1 |
| AURKB | 0 | 7,58958926 | 0,167 | 0,001 | 0 | Proliferating | AURKB |
| MELK | 0 | 6,72593333 | 0,167 | 0,002 | 0 | Proliferating | MELK |
| SPC25 | 0 | 9,73479839 | 0,163 | 0 | 0 | Proliferating | SPC25 |
| CDCA8 | 0 | 6,05742025 | 0,163 | 0,002 | 0 | Proliferating | CDCA8 |
| CHEK12 | 0 | 4,72289212 | 0,165 | 0,006 | 0 | Proliferating | CHEK1 |
| MYBL2 | 0 | 4,89674283 | 0,163 | 0,004 | 0 | Proliferating | MYBL2 |
| ECT2 | 0 | 5,03094804 | 0,163 | 0,005 | 0 | Proliferating | ECT2 |
| CIT | 0 | 6,17797434 | 0,16 | 0,002 | 0 | Proliferating | CIT |
| ANLN | 0 | 6,97343558 | 0,158 | 0,001 | 0 | Proliferating | ANLN |
| GGH | 0 | 5,32045466 | 0,16 | 0,004 | 0 | Proliferating | GGH |
| CDKN3 | 0 | 5,47787888 | 0,16 | 0,004 | 0 | Proliferating | CDKN3 |
| E2F1 | 0 | 5,63035567 | 0,154 | 0,003 | 0 | Proliferating | E2F1 |
| TROAP | 0 | 8,29773458 | 0,147 | 0 | 0 | Proliferating | TROAP |

| row.names | p_val | avg_log2FC | pct.1 | pct.2 | p_val_adj | cluster | gene |
|---|---|---|---|---|---|---|---|
| GSN2 | 0 | 8,31766807 | 0,871 | 0,037 | 0 | Fibroblasts | GSN |
| TIMP31 | 0 | 9,47458934 | 0,819 | 0,002 | 0 | Fibroblasts | TIMP3 |
| IFITM34 | 0 | 4,45560346 | 0,896 | 0,086 | 0 | Fibroblasts | IFITM3 |
| IGFBP73 | 0 | 6,05627343 | 0,799 | 0,051 | 0 | Fibroblasts | IGFBP7 |
| CAV11 | 0 | 10,5254354 | 0,738 | 0,001 | 0 | Fibroblasts | CAV1 |
| ADIRF1 | 0 | 9,40547125 | 0,709 | 0,003 | 0 | Fibroblasts | ADIRF |
| MGP | 0 | 11,6739947 | 0,704 | 0,002 | 0 | Fibroblasts | MGP |
| LGALS35 | 0 | 3,6563349 | 0,826 | 0,149 | 0 | Fibroblasts | LGALS3 |
| IGFBP44 | 0 | 6,09644038 | 0,691 | 0,022 | 0 | Fibroblasts | IGFBP4 |
| SELENOP2 | 0 | 3,92326163 | 0,682 | 0,024 | 0 | Fibroblasts | SELENOP |
| CXCL121 | 0 | 9,58527297 | 0,648 | 0,003 | 0 | Fibroblasts | CXCL12 |
| PLAC9 | 0 | 9,66814192 | 0,643 | 0,002 | 0 | Fibroblasts | PLAC9 |
| APOD | 0 | 12,2386986 | 0,639 | 0,002 | 0 | Fibroblasts | APOD |
| SERPING11 | 0 | 9,01414362 | 0,639 | 0,003 | 0 | Fibroblasts | SERPING1 |
| S100A133 | 0 | 7,80419041 | 0,641 | 0,006 | 0 | Fibroblasts | S100A13 |
| SPARC1 | 0 | 9,58488447 | 0,63 | 0,001 | 0 | Fibroblasts | SPARC |
| CALD12 | 0 | 8,38803931 | 0,621 | 0,004 | 0 | Fibroblasts | CALD1 |
| DCN | 0 | 12,8455935 | 0,605 | 0,002 | 0 | Fibroblasts | DCN |
| LAPTM4A4 | 0 | 3,58995901 | 0,718 | 0,116 | 0 | Fibroblasts | LAPTM4A |
| COL1A2 | 0 | 11,8846022 | 0,6 | 0,001 | 0 | Fibroblasts | COL1A2 |
| COL6A23 | 0 | 6,42594093 | 0,628 | 0,035 | 0 | Fibroblasts | COL6A2 |
| LHFPL64 | 0 | 4,73753286 | 0,607 | 0,029 | 0 | Fibroblasts | LHFPL6 |
| CFD3 | 0 | 8,51725472 | 0,632 | 0,054 | 0 | Fibroblasts | CFD |
| NNMT1 | 0 | 7,72686041 | 0,582 | 0,007 | 0 | Fibroblasts | NNMT |
| NFIB1 | 0 | 8,79871992 | 0,569 | 0,001 | 0 | Fibroblasts | NFIB |
| COL3A11 | 0 | 11,4655283 | 0,562 | 0,001 | 0 | Fibroblasts | COL3A1 |
| LUM | 0 | 12,6741787 | 0,56 | 0 | 0 | Fibroblasts | LUM |
| SOD3 | 0 | 12,7900303 | 0,558 | 0 | 0 | Fibroblasts | SOD3 |
| CFH1 | 0 | 7,50119012 | 0,567 | 0,013 | 0 | Fibroblasts | CFH |
| CCDC801 | 0 | 10,6586883 | 0,553 | 0,001 | 0 | Fibroblasts | CCDC80 |
| PRRX11 | 0 | 9,94786032 | 0,546 | 0,001 | 0 | Fibroblasts | PRRX1 |
| COL6A11 | 0 | 8,83824828 | 0,549 | 0,004 | 0 | Fibroblasts | COL6A1 |
| SERPINF11 | 0 | 7,13313239 | 0,551 | 0,01 | 0 | Fibroblasts | SERPINF1 |
| GNG111 | 0 | 9,92623607 | 0,54 | 0,002 | 0 | Fibroblasts | GNG11 |
| SELENOM2 | 0 | 4,53284251 | 0,582 | 0,044 | 0 | Fibroblasts | SELENOM |
| C1R | 0 | 10,918728 | 0,537 | 0,001 | 0 | Fibroblasts | C1R |
| C31 | 0 | 10,1954889 | 0,537 | 0,002 | 0 | Fibroblasts | C3 |
| MT1M1 | 0 | 8,23870514 | 0,533 | 0,002 | 0 | Fibroblasts | MT1M |
| COL6A31 | 0 | 9,92885383 | 0,53 | 0,002 | 0 | Fibroblasts | COL6A3 |
| C1S | 0 | 6,76655687 | 0,542 | 0,015 | 0 | Fibroblasts | C1S |
| MT1E3 | 0 | 4,12837738 | 0,564 | 0,039 | 0 | Fibroblasts | MT1E |
| PDK43 | 0 | 3,16354826 | 0,567 | 0,047 | 0 | Fibroblasts | PDK4 |
| RBMS32 | 0 | 7,00362594 | 0,53 | 0,011 | 0 | Fibroblasts | RBMS3 |
| PLPP32 | 0 | 8,10962855 | 0,519 | 0,004 | 0 | Fibroblasts | PLPP3 |
| EBF12 | 0 | 5,17569947 | 0,528 | 0,014 | 0 | Fibroblasts | EBF1 |
| TCF44 | 0 | 3,32821122 | 0,569 | 0,055 | 0 | Fibroblasts | TCF4 |
| NFIA2 | 0 | 4,47330565 | 0,555 | 0,042 | 0 | Fibroblasts | NFIA |
| TIMP24 | 0 | 4,28151306 | 0,562 | 0,051 | 0 | Fibroblasts | TIMP2 |
| SPARCL11 | 0 | 10,1322849 | 0,512 | 0,002 | 0 | Fibroblasts | SPARCL1 |
| CAVIN12 | 0 | 8,26706589 | 0,512 | 0,002 | 0 | Fibroblasts | CAVIN1 |
| FSTL11 | 0 | 10,6050797 | 0,51 | 0,001 | 0 | Fibroblasts | FSTL1 |
| CD594 | 0 | 4,25438172 | 0,551 | 0,043 | 0 | Fibroblasts | CD59 |
| CXCL14 | 0 | 12,6994675 | 0,499 | 0,001 | 0 | Fibroblasts | CXCL14 |
| NUPR11 | 0 | 10,1192816 | 0,497 | 0,001 | 0 | Fibroblasts | NUPR1 |
| NOVA11 | 0 | 11,1969831 | 0,494 | 0 | 0 | Fibroblasts | NOVA1 |
| COL1A1 | 0 | 9,46126279 | 0,497 | 0,003 | 0 | Fibroblasts | COL1A1 |
| ADH1B | 0 | 11,6840231 | 0,49 | 0 | 0 | Fibroblasts | ADH1B |
| IGFBP61 | 0 | 9,70080704 | 0,49 | 0,002 | 0 | Fibroblasts | IGFBP6 |
| IFI271 | 0 | 8,70194372 | 0,488 | 0,004 | 0 | Fibroblasts | IFI27 |
| FABP42 | 0 | 7,68829176 | 0,49 | 0,009 | 0 | Fibroblasts | FABP4 |
| CCN5 | 0 | 12,7410476 | 0,481 | 0 | 0 | Fibroblasts | CCN5 |
| MFAP4 | 0 | 11,7674817 | 0,479 | 0 | 0 | Fibroblasts | MFAP4 |
| FBLN2 | 0 | 8,56958534 | 0,481 | 0,003 | 0 | Fibroblasts | FBLN2 |
| EMP13 | 0 | 3,49489254 | 0,542 | 0,066 | 0 | Fibroblasts | EMP1 |
| LRP15 | 0 | 5,2670911 | 0,497 | 0,022 | 0 | Fibroblasts | LRP1 |
| EPHX11 | 0 | 7,52179652 | 0,474 | 0,004 | 0 | Fibroblasts | EPHX1 |
| HSPB14 | 0 | 3,43141397 | 0,526 | 0,058 | 0 | Fibroblasts | HSPB1 |
| MGST11 | 0 | 7,7554269 | 0,467 | 0,004 | 0 | Fibroblasts | MGST1 |
| GPC3 | 0 | 11,2008184 | 0,463 | 0,001 | 0 | Fibroblasts | GPC3 |
| DLC11 | 0 | 7,66157783 | 0,463 | 0,003 | 0 | Fibroblasts | DLC1 |
| SLIT31 | 0 | 7,97040289 | 0,458 | 0,002 | 0 | Fibroblasts | SLIT3 |
| FBLN1 | 0 | 11,5994206 | 0,456 | 0 | 0 | Fibroblasts | FBLN1 |
| GPX31 | 0 | 8,77028269 | 0,456 | 0,003 | 0 | Fibroblasts | GPX3 |
| RAMP2 | 0 | 11,409098 | 0,445 | 0 | 0 | Fibroblasts | RAMP2 |

| row.names | p_val | avg_log2FC | pct.1 | pct.2 | p_val_adj | cluster | gene |
|---|---|---|---|---|---|---|---|
| FXYD22 | 0 | 4,12534545 | 0,445 | 0,033 | 0 | Naïve CD8+ T | FXYD2 |
| COBL | 0 | 8,75725379 | 0,186 | 0,001 | 0 | Naïve CD8+ T | COBL |
| ZNF6833 | 0 | 4,64158257 | 0,188 | 0,008 | 0 | Naïve CD8+ T | ZNF683 |
| KLRC37 | 3,3E-307 | 2,93894887 | 0,647 | 0,1 | 8,322E-303 | Naïve CD8+ T | KLRC3 |
| LEF13 | 1,5E-267 | 3,1545047 | 0,471 | 0,061 | 3,622E-263 | Naïve CD8+ T | LEF1 |
| LINC024464 | 1,6E-234 | 3,72115855 | 0,311 | 0,031 | 4,093E-230 | Naïve CD8+ T | LINC02446 |
| RASSF86 | 2,4E-216 | 3,46396883 | 0,313 | 0,033 | 6,065E-212 | Naïve CD8+ T | RASSF8 |
| TFCP2L13 | 3,8E-198 | 4,64077024 | 0,108 | 0,004 | 9,459E-194 | Naïve CD8+ T | TFCP2L1 |
| KLRC26 | 2,2E-195 | 2,89399815 | 0,424 | 0,064 | 5,493E-191 | Naïve CD8+ T | KLRC2 |
| USP6NL4 | 2,9E-178 | 3,52076599 | 0,231 | 0,022 | 7,172E-174 | Naïve CD8+ T | USP6NL |
| CHPT13 | 2,6E-170 | 2,70473609 | 0,473 | 0,089 | 6,545E-166 | Naïve CD8+ T | CHPT1 |
| SPRY25 | 1,4E-168 | 3,78880852 | 0,174 | 0,013 | 3,509E-164 | Naïve CD8+ T | SPRY2 |
| CCDC571 | 1,1E-166 | 2,95625436 | 0,36 | 0,054 | 2,683E-162 | Naïve CD8+ T | CCDC57 |
| AKAP56 | 5,8E-147 | 3,57505045 | 0,167 | 0,014 | 1,435E-142 | Naïve CD8+ T | AKAP5 |
| IFNG-AS15 | 8,4E-147 | 3,09578139 | 0,332 | 0,052 | 2,097E-142 | Naïve CD8+ T | IFNG-AS1 |
| MME1 | 4,1E-145 | 5,1873579 | 0,056 | 0,002 | 1,012E-140 | Naïve CD8+ T | MME |
| KLRK17 | 2,5E-134 | 1,90448768 | 0,671 | 0,198 | 6,255E-130 | Naïve CD8+ T | KLRK1 |
| CCM24 | 1,2E-127 | 2,42858452 | 0,562 | 0,158 | 3,101E-123 | Naïve CD8+ T | CCM2 |
| GPCPD13 | 2,6E-126 | 2,03496078 | 0,802 | 0,362 | 6,575E-122 | Naïve CD8+ T | GPCPD1 |
| IL21R4 | 1,2E-121 | 2,30254439 | 0,496 | 0,126 | 2,878E-117 | Naïve CD8+ T | IL21R |
| CD76 | 1,9E-121 | 1,68319462 | 0,786 | 0,298 | 4,696E-117 | Naïve CD8+ T | CD7 |
| EZR3 | 1,8E-118 | 1,46530349 | 0,951 | 0,628 | 4,424E-114 | Naïve CD8+ T | EZR |
| VAV35 | 5,1E-117 | 1,76251965 | 0,788 | 0,318 | 1,275E-112 | Naïve CD8+ T | VAV3 |
| ZFPM15 | 4,4E-110 | 2,6542325 | 0,308 | 0,057 | 1,109E-105 | Naïve CD8+ T | ZFPM1 |
| TRGC24 | 3,8E-103 | 2,07441068 | 0,431 | 0,107 | 9,541E-99 | Naïve CD8+ T | TRGC2 |
| PDE9A4 | 8,65E-97 | 3,98740664 | 0,087 | 0,006 | 2,1584E-92 | Naïve CD8+ T | PDE9A |
| BACH27 | 7,12E-96 | 1,48704877 | 0,835 | 0,411 | 1,7757E-91 | Naïve CD8+ T | BACH2 |
| IKZF25 | 2,5E-95 | 1,75926362 | 0,449 | 0,118 | 6,2343E-91 | Naïve CD8+ T | IKZF2 |
| LMO75 | 2,65E-95 | 3,07743464 | 0,179 | 0,023 | 6,6192E-91 | Naïve CD8+ T | LMO7 |
| SHLD12 | 6,61E-93 | 2,55487131 | 0,254 | 0,045 | 1,6507E-88 | Naïve CD8+ T | SHLD1 |
| PABPC14 | 7,01E-90 | 1,24233661 | 0,934 | 0,721 | 1,7489E-85 | Naïve CD8+ T | PABPC1 |
| SCML17 | 4,77E-88 | 2,54490469 | 0,254 | 0,047 | 1,1903E-83 | Naïve CD8+ T | SCML1 |
| KLF24 | 6,06E-84 | 1,51109296 | 0,845 | 0,443 | 1,5117E-79 | Naïve CD8+ T | KLF2 |
| CD556 | 6,31E-84 | 1,36290673 | 0,826 | 0,434 | 1,5759E-79 | Naïve CD8+ T | CD55 |
| LITAF5 | 2,08E-81 | 1,39725766 | 0,798 | 0,427 | 5,1816E-77 | Naïve CD8+ T | LITAF |
| CLASP23 | 1,11E-80 | 2,00353213 | 0,546 | 0,203 | 2,7581E-76 | Naïve CD8+ T | CLASP2 |
| KLRC46 | 4,62E-80 | 2,59896282 | 0,193 | 0,031 | 1,1532E-75 | Naïve CD8+ T | KLRC4 |
| ITPKB2 | 2,74E-79 | 1,94247068 | 0,482 | 0,16 | 6,8408E-75 | Naïve CD8+ T | ITPKB |
| CASS43 | 4,11E-78 | 2,15591623 | 0,339 | 0,085 | 1,0245E-73 | Naïve CD8+ T | CASS4 |
| LYAR5 | 4,9E-75 | 1,83147326 | 0,447 | 0,142 | 1,2241E-70 | Naïve CD8+ T | LYAR |
| EPHA45 | 4,2E-73 | 2,1203517 | 0,332 | 0,086 | 1,0486E-68 | Naïve CD8+ T | EPHA4 |
| ZFP36L23 | 1,69E-72 | 0,92563343 | 0,993 | 0,769 | 4,2159E-68 | Naïve CD8+ T | ZFP36L2 |
| TCF76 | 4,44E-71 | 2,13646689 | 0,289 | 0,069 | 1,109E-66 | Naïve CD8+ T | TCF7 |
| PHF202 | 1,08E-70 | 1,36586075 | 0,793 | 0,467 | 2,6911E-66 | Naïve CD8+ T | PHF20 |
| CEMIP23 | 3,09E-70 | 1,49082034 | 0,88 | 0,591 | 7,7191E-66 | Naïve CD8+ T | CEMIP2 |
| FAM13A6 | 1,03E-67 | 2,42664329 | 0,226 | 0,047 | 2,5701E-63 | Naïve CD8+ T | FAM13A |
| PIK3R13 | 3,79E-65 | 1,21592939 | 0,816 | 0,509 | 9,4492E-61 | Naïve CD8+ T | PIK3R1 |
| GEM7 | 1,78E-62 | 3,11354362 | 0,118 | 0,015 | 4,4455E-58 | Naïve CD8+ T | GEM |
| ZHX26 | 3,31E-60 | 1,7778801 | 0,569 | 0,264 | 8,2675E-56 | Naïve CD8+ T | ZHX2 |
| ZNF7045 | 3,8E-60 | 2,47793965 | 0,176 | 0,033 | 9,4883E-56 | Naïve CD8+ T | ZNF704 |
| MAP3K17 | 1,26E-58 | 1,88704119 | 0,365 | 0,116 | 3,1537E-54 | Naïve CD8+ T | MAP3K1 |
| SEMA4D3 | 2,31E-58 | 1,600239 | 0,487 | 0,194 | 5,7668E-54 | Naïve CD8+ T | SEMA4D |
| SPTBN15 | 4,58E-58 | 2,01023916 | 0,28 | 0,074 | 1,1421E-53 | Naïve CD8+ T | SPTBN1 |
| KLF34 | 4,6E-58 | 1,36942285 | 0,588 | 0,263 | 1,1479E-53 | Naïve CD8+ T | KLF3 |
| JUND2 | 2,92E-57 | 0,79166154 | 0,974 | 0,853 | 7,2784E-53 | Naïve CD8+ T | JUND |
| ESYT23 | 7,67E-57 | 1,33678456 | 0,692 | 0,376 | 1,9147E-52 | Naïve CD8+ T | ESYT2 |
| CAST2 | 1,3E-56 | 1,44069182 | 0,699 | 0,419 | 3,2403E-52 | Naïve CD8+ T | CAST |
| TXK6 | 2,1E-56 | 1,85151305 | 0,332 | 0,1 | 5,2307E-52 | Naïve CD8+ T | TXK |
| PDE3B4 | 6,94E-56 | 1,13057706 | 0,734 | 0,377 | 1,7314E-51 | Naïve CD8+ T | PDE3B |
| LMBR11 | 1,46E-55 | 1,79838787 | 0,409 | 0,148 | 3,6465E-51 | Naïve CD8+ T | LMBR1 |
| ABLIM16 | 3,11E-55 | 1,77439649 | 0,464 | 0,181 | 7,7735E-51 | Naïve CD8+ T | ABLIM1 |
| DPF36 | 1,43E-54 | 3,43354324 | 0,08 | 0,009 | 3,5576E-50 | Naïve CD8+ T | DPF3 |
| C19orf334 | 1,44E-54 | 3,91945837 | 0,047 | 0,003 | 3,5845E-50 | Naïve CD8+ T | C19orf33 |
| AKT31 | 5,82E-54 | 1,29816991 | 0,678 | 0,345 | 1,4514E-49 | Naïve CD8+ T | AKT3 |
| SH3BGRL23 | 1,58E-53 | 5,12485772 | 0,024 | 0,001 | 3,9469E-49 | Naïve CD8+ T | SH3BGRL2 |
| AL109935.12 | 5,37E-51 | 3,23610638 | 0,08 | 0,009 | 1,3399E-46 | Naïve CD8+ T | AL109935.1 |
| TMEM1634 | 6,4E-51 | 2,69902082 | 0,12 | 0,019 | 1,598E-46 | Naïve CD8+ T | TMEM163 |
| GNG45 | 7,92E-51 | 3,02812266 | 0,125 | 0,02 | 1,9766E-46 | Naïve CD8+ T | GNG4 |
| DRICH1 | 2,79E-50 | 5,11073661 | 0,021 | 0,001 | 6,957E-46 | Naïve CD8+ T | DRICH1 |
| TNFSF83 | 4,99E-50 | 1,45681881 | 0,44 | 0,171 | 1,245E-45 | Naïve CD8+ T | TNFSF8 |
| TUBA4A3 | 9,97E-50 | 1,17735817 | 0,694 | 0,387 | 2,4894E-45 | Naïve CD8+ T | TUBA4A |
| PPP2R5C6 | 1,97E-48 | 0,8758558 | 0,899 | 0,604 | 4,9265E-44 | Naïve CD8+ T | PPP2R5C |
| RALGPS24 | 4,36E-48 | 1,99095369 | 0,184 | 0,041 | 1,0876E-43 | Naïve CD8+ T | RALGPS2 |
| EML44 | 5,79E-48 | 0,98763931 | 0,816 | 0,528 | 1,4446E-43 | Naïve CD8+ T | EML4 |

| row.names | p_val | avg_log2FC | pct.1 | pct.2 | p_val_adj | cluster | gene |
|---|---|---|---|---|---|---|---|
| C1orf544 | 0 | 7,05165614 | 0,928 | 0,018 | 0 | cDCs | C1orf54 |
| CPVL3 | 0 | 5,7683928 | 0,957 | 0,065 | 0 | cDCs | CPVL |
| WDFY43 | 0 | 7,33709314 | 0,903 | 0,012 | 0 | cDCs | WDFY4 |
| CPNE31 | 0 | 5,16602192 | 0,914 | 0,054 | 0 | cDCs | CPNE3 |
| LYZ2 | 0 | 2,49362009 | 0,973 | 0,113 | 0 | cDCs | LYZ |
| HLA-DQA14 | 0 | 3,66886414 | 0,984 | 0,151 | 0 | cDCs | HLA-DQA1 |
| CST35 | 0 | 4,08472253 | 0,984 | 0,152 | 0 | cDCs | CST3 |
| SNX36 | 0 | 4,54368512 | 0,941 | 0,11 | 0 | cDCs | SNX3 |
| CLEC9A | 0 | 11,6076129 | 0,815 | 0 | 0 | cDCs | CLEC9A |
| C1orf1623 | 0 | 3,67750224 | 0,887 | 0,103 | 0 | cDCs | C1orf162 |
| BASP14 | 0 | 3,87177489 | 0,853 | 0,07 | 0 | cDCs | BASP1 |
| HLA-DMA4 | 0 | 3,00759717 | 0,906 | 0,124 | 0 | cDCs | HLA-DMA |
| HLA-DQB14 | 0 | 3,74555506 | 0,979 | 0,204 | 0 | cDCs | HLA-DQB1 |
| DNASE1L31 | 0 | 10,1286518 | 0,772 | 0,001 | 0 | cDCs | DNASE1L3 |
| CXCL164 | 0 | 3,80122508 | 0,845 | 0,074 | 0 | cDCs | CXCL16 |
| CCSER13 | 0 | 5,09962399 | 0,826 | 0,056 | 0 | cDCs | CCSER1 |
| HLA-DRA5 | 0 | 3,57461912 | 0,995 | 0,228 | 0 | cDCs | HLA-DRA |
| HLA-DPA15 | 0 | 4,32093409 | 0,989 | 0,23 | 0 | cDCs | HLA-DPA1 |
| LGALS22 | 0 | 5,01111985 | 0,794 | 0,036 | 0 | cDCs | LGALS2 |
| SHTN13 | 0 | 5,56620267 | 0,78 | 0,024 | 0 | cDCs | SHTN1 |
| RGS104 | 0 | 3,2939389 | 0,882 | 0,132 | 0 | cDCs | RGS10 |
| HDAC95 | 0 | 4,70079376 | 0,812 | 0,066 | 0 | cDCs | HDAC9 |
| HLA-DPB14 | 0 | 4,29661413 | 0,992 | 0,251 | 0 | cDCs | HLA-DPB1 |
| CTSZ4 | 0 | 3,27092189 | 0,839 | 0,106 | 0 | cDCs | CTSZ |
| SIPA1L32 | 0 | 5,06807137 | 0,767 | 0,056 | 0 | cDCs | SIPA1L3 |
| NRARP3 | 0 | 5,83011586 | 0,732 | 0,025 | 0 | cDCs | NRARP |
| CSF2RA4 | 0 | 3,52712051 | 0,777 | 0,079 | 0 | cDCs | CSF2RA |
| IDO11 | 0 | 9,10254998 | 0,7 | 0,002 | 0 | cDCs | IDO1 |
| TACSTD2 | 0 | 11,1180123 | 0,694 | 0,001 | 0 | cDCs | TACSTD2 |
| RAB11FIP15 | 0 | 3,22858231 | 0,866 | 0,177 | 0 | cDCs | RAB11FIP1 |
| HLA-DRB14 | 0 | 3,56044991 | 0,989 | 0,301 | 0 | cDCs | HLA-DRB1 |
| RNF144B4 | 0 | 2,95454599 | 0,788 | 0,11 | 0 | cDCs | RNF144B |
| CDK2AP14 | 0 | 4,672021 | 0,705 | 0,032 | 0 | cDCs | CDK2AP1 |
| HLA-DRB54 | 0 | 3,61577852 | 0,799 | 0,142 | 0 | cDCs | HLA-DRB5 |
| CLEC7A4 | 0 | 3,43302354 | 0,721 | 0,07 | 0 | cDCs | CLEC7A |
| CADM14 | 0 | 3,60394949 | 0,721 | 0,074 | 0 | cDCs | CADM1 |
| UPF22 | 0 | 3,43671241 | 0,745 | 0,104 | 0 | cDCs | UPF2 |
| CD744 | 0 | 3,96963624 | 0,997 | 0,372 | 0 | cDCs | CD74 |
| PLEKHA53 | 0 | 4,57299435 | 0,694 | 0,083 | 0 | cDCs | PLEKHA5 |
| MIR181A1HG | 0 | 3,13750766 | 0,697 | 0,094 | 0 | cDCs | MIR181A1HG |
| CCDC64 | 0 | 3,20236235 | 0,697 | 0,094 | 0 | cDCs | CCDC6 |
| RAB7B3 | 0 | 7,97018301 | 0,606 | 0,004 | 0 | cDCs | RAB7B |
| DAPP14 | 0 | 4,38338151 | 0,63 | 0,041 | 0 | cDCs | DAPP1 |
| PKP24 | 0 | 3,90327087 | 0,641 | 0,052 | 0 | cDCs | PKP2 |
| ZBTB464 | 0 | 6,58726429 | 0,595 | 0,01 | 0 | cDCs | ZBTB46 |
| DST6 | 0 | 5,02595616 | 0,606 | 0,023 | 0 | cDCs | DST |
| NEGR13 | 0 | 5,11008153 | 0,582 | 0,015 | 0 | cDCs | NEGR1 |
| RNASE63 | 0 | 4,5475141 | 0,603 | 0,037 | 0 | cDCs | RNASE6 |
| IL1R23 | 0 | 2,57498035 | 0,633 | 0,071 | 0 | cDCs | IL1R2 |
| NET12 | 0 | 5,00782943 | 0,587 | 0,025 | 0 | cDCs | NET1 |
| ZFHX36 | 0 | 4,14671918 | 0,598 | 0,043 | 0 | cDCs | ZFHX3 |
| THRB5 | 0 | 5,06361234 | 0,563 | 0,024 | 0 | cDCs | THRB |
| GNA125 | 0 | 3,35108911 | 0,614 | 0,077 | 0 | cDCs | GNA12 |
| PPT14 | 0 | 3,60986891 | 0,579 | 0,052 | 0 | cDCs | PPT1 |
| IRF87 | 0 | 3,54258154 | 0,587 | 0,06 | 0 | cDCs | IRF8 |
| PPP1R14B5 | 0 | 3,32070241 | 0,576 | 0,05 | 0 | cDCs | PPP1R14B |
| CTTNBP2NL5 | 0 | 3,90589025 | 0,563 | 0,043 | 0 | cDCs | CTTNBP2NL |
| AC099560.11 | 0 | 8,86299377 | 0,52 | 0,002 | 0 | cDCs | AC099560.1 |
| VDR3 | 0 | 4,59860932 | 0,547 | 0,03 | 0 | cDCs | VDR |
| MEF2C6 | 0 | 3,46404029 | 0,584 | 0,067 | 0 | cDCs | MEF2C |
| GPR1572 | 0 | 5,5683713 | 0,517 | 0,015 | 0 | cDCs | GPR157 |
| RUBCNL2 | 0 | 5,81650524 | 0,491 | 0,009 | 0 | cDCs | RUBCNL |
| PKIB2 | 0 | 4,86363021 | 0,496 | 0,02 | 0 | cDCs | PKIB |
| KIF16B3 | 0 | 4,38258333 | 0,485 | 0,03 | 0 | cDCs | KIF16B |
| SDK14 | 0 | 6,6556441 | 0,456 | 0,009 | 0 | cDCs | SDK1 |
| FRY6 | 0 | 4,17943546 | 0,499 | 0,055 | 0 | cDCs | FRY |
| PADI21 | 0 | 6,78561124 | 0,445 | 0,006 | 0 | cDCs | PADI2 |
| S100B6 | 0 | 6,47640993 | 0,44 | 0,009 | 0 | cDCs | S100B |
| CALCRL4 | 0 | 5,67681915 | 0,442 | 0,014 | 0 | cDCs | CALCRL |
| KLHL67 | 0 | 3,8460617 | 0,461 | 0,037 | 0 | cDCs | KLHL6 |
| SLC41A23 | 0 | 3,67748663 | 0,466 | 0,044 | 0 | cDCs | SLC41A2 |
| LDLRAD35 | 0 | 4,42333264 | 0,445 | 0,025 | 0 | cDCs | LDLRAD3 |
| CLIC24 | 0 | 4,80315133 | 0,434 | 0,018 | 0 | cDCs | CLIC2 |
| THBD4 | 0 | 3,71005204 | 0,45 | 0,035 | 0 | cDCs | THBD |

| row.names | p_val | avg_log2FC | pct.1 | pct.2 | p_val_adj | cluster | gene |
|---|---|---|---|---|---|---|---|
| RHEX2 | 0 | 8,58759687 | 0,963 | 0,005 | 0 | pDCs | RHEX |
| JCHAIN1 | 0 | 4,72103613 | 0,955 | 0,006 | 0 | pDCs | JCHAIN |
| IL3RA4 | 0 | 7,29575065 | 0,958 | 0,016 | 0 | pDCs | IL3RA |
| PPP1R14B6 | 0 | 5,90600414 | 0,941 | 0,048 | 0 | pDCs | PPP1R14B |
| SEL1L31 | 0 | 6,36472402 | 0,946 | 0,055 | 0 | pDCs | SEL1L3 |
| TCF46 | 0 | 5,19748563 | 0,93 | 0,054 | 0 | pDCs | TCF4 |
| IRF42 | 0 | 5,87124004 | 0,961 | 0,089 | 0 | pDCs | IRF4 |
| SOX46 | 0 | 7,01539606 | 0,887 | 0,026 | 0 | pDCs | SOX4 |
| ITM2C3 | 0 | 6,1435828 | 0,885 | 0,026 | 0 | pDCs | ITM2C |
| FCHSD23 | 0 | 5,69174812 | 0,958 | 0,117 | 0 | pDCs | FCHSD2 |
| RUBCN2 | 0 | 5,22175988 | 0,896 | 0,057 | 0 | pDCs | RUBCN |
| SLC15A43 | 0 | 5,08119955 | 0,893 | 0,063 | 0 | pDCs | SLC15A4 |
| PTPRS1 | 0 | 8,03399705 | 0,834 | 0,004 | 0 | pDCs | PTPRS |
| APP5 | 0 | 5,1037857 | 0,887 | 0,057 | 0 | pDCs | APP |
| CD2AP5 | 0 | 4,82845356 | 0,907 | 0,08 | 0 | pDCs | CD2AP |
| RRBP18 | 0 | 4,53179266 | 0,901 | 0,09 | 0 | pDCs | RRBP1 |
| EGLN31 | 0 | 7,62225409 | 0,817 | 0,007 | 0 | pDCs | EGLN3 |
| FAM160A14 | 0 | 6,2554568 | 0,834 | 0,029 | 0 | pDCs | FAM160A1 |
| CORO1C4 | 0 | 5,00809299 | 0,854 | 0,052 | 0 | pDCs | CORO1C |
| POLB1 | 0 | 5,27601217 | 0,868 | 0,07 | 0 | pDCs | POLB |
| IRF88 | 0 | 4,92416805 | 0,856 | 0,059 | 0 | pDCs | IRF8 |
| AFF35 | 0 | 4,19693055 | 0,856 | 0,065 | 0 | pDCs | AFF3 |
| C12orf756 | 0 | 4,05348634 | 0,944 | 0,156 | 0 | pDCs | C12orf75 |
| PLD42 | 0 | 8,43194649 | 0,789 | 0,004 | 0 | pDCs | PLD4 |
| CTSB4 | 0 | 3,60423249 | 0,893 | 0,111 | 0 | pDCs | CTSB |
| FYTTD11 | 0 | 4,57468648 | 0,854 | 0,079 | 0 | pDCs | FYTTD1 |
| N4BP2L11 | 0 | 4,60082496 | 0,839 | 0,067 | 0 | pDCs | N4BP2L1 |
| NPC26 | 0 | 3,39334129 | 0,893 | 0,131 | 0 | pDCs | NPC2 |
| SEC61B | 0 | 4,06194487 | 0,975 | 0,22 | 0 | pDCs | SEC61B |
| AC021594.24 | 0 | 8,437234 | 0,755 | 0,004 | 0 | pDCs | AC021594.2 |
| P2RY62 | 0 | 9,40056185 | 0,752 | 0,002 | 0 | pDCs | P2RY6 |
| RNASET27 | 0 | 3,40393318 | 0,882 | 0,137 | 0 | pDCs | RNASET2 |
| IRF73 | 0 | 5,11601018 | 0,792 | 0,048 | 0 | pDCs | IRF7 |
| SULF24 | 0 | 4,70744437 | 0,792 | 0,049 | 0 | pDCs | SULF2 |
| GRASP9 | 0 | 3,40556811 | 0,918 | 0,181 | 0 | pDCs | GRASP |
| PMEPA14 | 0 | 4,63352335 | 0,783 | 0,052 | 0 | pDCs | PMEPA1 |
| THEMIS27 | 0 | 3,66614085 | 0,8 | 0,081 | 0 | pDCs | THEMIS2 |
| INPP4A5 | 0 | 3,53583341 | 0,856 | 0,142 | 0 | pDCs | INPP4A |
| AC023590.12 | 0 | 8,1234646 | 0,713 | 0,006 | 0 | pDCs | AC023590.1 |
| GNA126 | 0 | 4,17829081 | 0,78 | 0,077 | 0 | pDCs | GNA12 |
| GAB19 | 0 | 5,61738729 | 0,715 | 0,026 | 0 | pDCs | GAB1 |
| SERPINF13 | 0 | 5,82777192 | 0,696 | 0,01 | 0 | pDCs | SERPINF1 |
| CDYL6 | 0 | 3,93723811 | 0,775 | 0,096 | 0 | pDCs | CDYL |
| LINC013743 | 0 | 7,62336565 | 0,685 | 0,006 | 0 | pDCs | LINC01374 |
| BCL2L115 | 0 | 3,10513844 | 0,803 | 0,132 | 0 | pDCs | BCL2L11 |
| PBX32 | 0 | 4,47551797 | 0,749 | 0,088 | 0 | pDCs | PBX3 |
| TGFBR14 | 0 | 3,94863369 | 0,738 | 0,084 | 0 | pDCs | TGFBR1 |
| TSPAN133 | 0 | 8,07728438 | 0,654 | 0,003 | 0 | pDCs | TSPAN13 |
| RASD12 | 0 | 6,18396015 | 0,67 | 0,02 | 0 | pDCs | RASD1 |
| BID4 | 0 | 3,38614654 | 0,746 | 0,1 | 0 | pDCs | BID |
| NUTM2B-AS1 | 0 | 3,71594112 | 0,738 | 0,094 | 0 | pDCs | NUTM2B-AS1 |
| MZB11 | 0 | 7,08257872 | 0,645 | 0,004 | 0 | pDCs | MZB1 |
| VEGFB5 | 0 | 4,41213543 | 0,679 | 0,038 | 0 | pDCs | VEGFB |
| CCDC506 | 0 | 5,21011178 | 0,668 | 0,029 | 0 | pDCs | CCDC50 |
| EIF2AK44 | 0 | 4,07099664 | 0,682 | 0,051 | 0 | pDCs | EIF2AK4 |
| GALNT24 | 0 | 4,31518404 | 0,685 | 0,055 | 0 | pDCs | GALNT2 |
| CYBB4 | 0 | 3,65523244 | 0,687 | 0,057 | 0 | pDCs | CYBB |
| SLC41A24 | 0 | 5,00852346 | 0,67 | 0,043 | 0 | pDCs | SLC41A2 |
| LILRA4 | 0 | 10,7948483 | 0,625 | 0 | 0 | pDCs | LILRA4 |
| GNA157 | 0 | 3,92682968 | 0,679 | 0,063 | 0 | pDCs | GNA15 |
| TENT4A1 | 0 | 4,46235027 | 0,662 | 0,048 | 0 | pDCs | TENT4A |
| MYBL21 | 0 | 9,23233647 | 0,611 | 0,002 | 0 | pDCs | MYBL2 |
| TEX146 | 0 | 3,52483684 | 0,704 | 0,096 | 0 | pDCs | TEX14 |
| CLIC33 | 0 | 3,28360184 | 0,696 | 0,095 | 0 | pDCs | CLIC3 |
| SH2B35 | 0 | 3,5791388 | 0,668 | 0,072 | 0 | pDCs | SH2B3 |
| COBLL13 | 0 | 6,31296912 | 0,597 | 0,009 | 0 | pDCs | COBLL1 |
| SMPD31 | 0 | 7,71163491 | 0,586 | 0,003 | 0 | pDCs | SMPD3 |
| SMIM35 | 0 | 3,95633045 | 0,628 | 0,046 | 0 | pDCs | SMIM3 |
| SCT | 0 | 10,5431667 | 0,58 | 0 | 0 | pDCs | SCT |
| LINC014781 | 0 | 11,2395719 | 0,58 | 0,001 | 0 | pDCs | LINC01478 |
| PARVB4 | 0 | 3,76520994 | 0,634 | 0,056 | 0 | pDCs | PARVB |
| MOB1B3 | 0 | 4,8176712 | 0,623 | 0,046 | 0 | pDCs | MOB1B |
| C12orf452 | 0 | 5,12126012 | 0,606 | 0,029 | 0 | pDCs | C12orf45 |
| PHEX1 | 0 | 8,77939041 | 0,58 | 0,003 | 0 | pDCs | PHEX |

| row.names | p_val | avg_log2FC | pct.1 | pct.2 | p_val_adj | cluster | gene |
|---|---|---|---|---|---|---|---|
| MZB12 | 0 | 8,76130697 | 0,946 | 0,006 | 0 | Plasma B | MZB1 |
| JCHAIN2 | 0 | 11,8582525 | 0,932 | 0,01 | 0 | Plasma B | JCHAIN |
| TXNDC53 | 0 | 7,81034316 | 0,932 | 0,013 | 0 | Plasma B | TXNDC5 |
| IGKC1 | 0 | 13,2331097 | 0,784 | 0,027 | 0 | Plasma B | IGKC |
| DERL31 | 0 | 8,58527661 | 0,757 | 0,003 | 0 | Plasma B | DERL3 |
| COBLL14 | 0 | 6,31412488 | 0,662 | 0,011 | 0 | Plasma B | COBLL1 |
| CD79A1 | 0 | 5,63496373 | 0,635 | 0,014 | 0 | Plasma B | CD79A |
| IGLC11 | 0 | 11,6503863 | 0,554 | 0,002 | 0 | Plasma B | IGLC1 |
| IGHA1 | 0 | 13,7262563 | 0,554 | 0,006 | 0 | Plasma B | IGHA1 |
| POU2AF11 | 0 | 8,72089074 | 0,486 | 0,002 | 0 | Plasma B | POU2AF1 |
| IGHG3 | 0 | 14,1159213 | 0,392 | 0,001 | 0 | Plasma B | IGHG3 |
| IGHG11 | 0 | 13,3351644 | 0,365 | 0,002 | 0 | Plasma B | IGHG1 |
| IGHA21 | 0 | 12,8912632 | 0,311 | 0,001 | 0 | Plasma B | IGHA2 |
| CADPS22 | 0 | 7,63902358 | 0,27 | 0,002 | 0 | Plasma B | CADPS2 |
| IGHG41 | 0 | 13,1460336 | 0,257 | 0,001 | 0 | Plasma B | IGHG4 |
| IGHGP | 0 | 14,5705641 | 0,257 | 0,001 | 0 | Plasma B | IGHGP |
| FCRL51 | 0 | 7,57628783 | 0,23 | 0,001 | 0 | Plasma B | FCRL5 |
| IGLC31 | 0 | 12,2469851 | 0,23 | 0,002 | 0 | Plasma B | IGLC3 |
| TNFRSF17 | 0 | 12,2373533 | 0,216 | 0 | 0 | Plasma B | TNFRSF17 |
| IGHG21 | 0 | 14,0160814 | 0,162 | 0 | 0 | Plasma B | IGHG2 |
| SDC1 | 0 | 13,4783614 | 0,149 | 0 | 0 | Plasma B | SDC1 |
| AC026369.32 | 0 | 7,41535161 | 0,149 | 0,001 | 0 | Plasma B | AC026369.3 |
| IGKV1-5 | 0 | 14,3926315 | 0,108 | 0 | 0 | Plasma B | IGKV1-5 |
| AL391056.1 | 0 | 8,77792169 | 0,095 | 0 | 0 | Plasma B | AL391056.1 |
| TUBB2B | 0 | 9,31849007 | 0,081 | 0 | 0 | Plasma B | TUBB2B |
| IGLV3-1 | 0 | 12,7779217 | 0,068 | 0 | 0 | Plasma B | IGLV3-1 |
| ZNF215 | 0 | 9,19295919 | 0,068 | 0 | 0 | Plasma B | ZNF215 |
| IGHV3-7 | 0 | 14,450347 | 0,054 | 0 | 0 | Plasma B | IGHV3-7 |
| IGKV4-1 | 0 | 13,9873751 | 0,041 | 0 | 0 | Plasma B | IGKV4-1 |
| IGHV4-34 | 0 | 12,1929592 | 0,041 | 0 | 0 | Plasma B | IGHV4-34 |
| PPARGC1A | 0 | 11,1929592 | 0,041 | 0 | 0 | Plasma B | PPARGC1A |
| IGHV3-23 | 0 | 12,3628842 | 0,041 | 0 | 0 | Plasma B | IGHV3-23 |
| IGKV3-20 | 0 | 13,0998498 | 0,041 | 0 | 0 | Plasma B | IGKV3-20 |
| IGHV3-21 | 0 | 13,6848123 | 0,041 | 0 | 0 | Plasma B | IGHV3-21 |
| PLA2G2D | 0 | 11,3628842 | 0,027 | 0 | 0 | Plasma B | PLA2G2D |
| GNG75 | 1E-297 | 5,24017382 | 0,716 | 0,026 | 3,6E-293 | Plasma B | GNG7 |
| PRDX43 | 1E-257 | 5,15502973 | 0,622 | 0,023 | 2,7E-253 | Plasma B | PRDX4 |
| AC012236.1 | 3E-256 | 10,3628842 | 0,027 | 0 | 7,4E-252 | Plasma B | AC012236.1 |
| AC084734.1 | 3E-256 | 10,3628842 | 0,027 | 0 | 7,4E-252 | Plasma B | AC084734.1 |
| IGLV6-57 | 3E-256 | 10,3628842 | 0,027 | 0 | 7,4E-252 | Plasma B | IGLV6-57 |
| ITM2C4 | 4E-249 | 4,90088708 | 0,703 | 0,03 | 1E-244 | Plasma B | ITM2C |
| TNFRSF13B2 | 2E-244 | 5,64041816 | 0,311 | 0,006 | 4,2E-240 | Plasma B | TNFRSF13B |
| IGLC21 | 2E-243 | 13,0479503 | 0,284 | 0,005 | 4,4E-239 | Plasma B | IGLC2 |
| AC002072.1 | 2E-239 | 7,86038385 | 0,095 | 0 | 3,9E-235 | Plasma B | AC002072.1 |
| JSRP1 | 4E-237 | 6,77792169 | 0,189 | 0,002 | 9,9E-233 | Plasma B | JSRP1 |
| LINC014803 | 4E-221 | 7,17126412 | 0,203 | 0,003 | 1E-216 | Plasma B | LINC01480 |
| BLNK3 | 2E-208 | 6,40636282 | 0,203 | 0,003 | 5,2E-204 | Plasma B | BLNK |
| AC007009.1 | 6E-193 | 10,7779217 | 0,014 | 0 | 1,6E-188 | Plasma B | AC007009.1 |
| IGHEP1 | 2E-192 | 15,6023501 | 0,027 | 0 | 4,1E-188 | Plasma B | IGHEP1 |
| AC025434.1 | 6E-173 | 8,77792169 | 0,041 | 0 | 1,5E-168 | Plasma B | AC025434.1 |
| GREB11 | 3E-164 | 7,10807029 | 0,108 | 0,001 | 7,3E-160 | Plasma B | GREB1 |
| SEC11C4 | 3E-159 | 4,58785316 | 0,919 | 0,087 | 8,1E-155 | Plasma B | SEC11C |
| CARMIL15 | 1E-140 | 5,69139172 | 0,432 | 0,02 | 3,2E-136 | Plasma B | CARMIL1 |
| IGHM1 | 3E-134 | 11,8008221 | 0,297 | 0,01 | 8,5E-130 | Plasma B | IGHM |
| FAM30A2 | 2E-132 | 6,60000389 | 0,149 | 0,002 | 3,9E-128 | Plasma B | FAM30A |
| AL589693.14 | 2E-122 | 4,85426621 | 0,378 | 0,017 | 4,9E-118 | Plasma B | AL589693.1 |
| CHMP4C | 2E-115 | 7,87103109 | 0,041 | 0 | 3,8E-111 | Plasma B | CHMP4C |
| CST6 | 2E-108 | 8,19295919 | 0,041 | 0 | 5,5E-104 | Plasma B | CST6 |
| RALGPS27 | 2,5E-98 | 4,16321184 | 0,527 | 0,041 | 6,32E-94 | Plasma B | RALGPS2 |
| AC097375.3 | 3E-97 | 9,77792169 | 0,014 | 0 | 7,56E-93 | Plasma B | AC097375.3 |
| WASHC5-AS1 | 3E-97 | 9,77792169 | 0,014 | 0 | 7,56E-93 | Plasma B | WASHC5-AS1 |
| RAB306 | 1,9E-90 | 4,41292487 | 0,351 | 0,02 | 4,8E-86 | Plasma B | RAB30 |
| TRIB17 | 2,7E-87 | 4,18979885 | 0,392 | 0,026 | 6,81E-83 | Plasma B | TRIB1 |
| WNT16 | 3,4E-86 | 9,60799669 | 0,027 | 0 | 8,61E-82 | Plasma B | WNT16 |
| RHEX3 | 4,8E-85 | 5,22286067 | 0,243 | 0,01 | 1,2E-80 | Plasma B | RHEX |
| AC007569.15 | 1,1E-82 | 5,36288419 | 0,162 | 0,005 | 2,84E-78 | Plasma B | AC007569.1 |
| CD277 | 8,8E-81 | 3,63697575 | 0,5 | 0,044 | 2,2E-76 | Plasma B | CD27 |
| CREB3L24 | 8,1E-80 | 3,94742143 | 0,486 | 0,043 | 2,01E-75 | Plasma B | CREB3L2 |
| IGHE | 1,1E-77 | 16,2639339 | 0,027 | 0 | 2,74E-73 | Plasma B | IGHE |
| GAB110 | 1,3E-76 | 4,04364446 | 0,392 | 0,029 | 3,33E-72 | Plasma B | GAB1 |
| AP002852.1 | 3,2E-75 | 6,33497819 | 0,081 | 0,001 | 7,97E-71 | Plasma B | AP002852.1 |
| TLCD41 | 1,8E-72 | 8,48841507 | 0,041 | 0 | 4,47E-68 | Plasma B | TLCD4 |
| IGLV10-54 | 1,8E-72 | 7,64041816 | 0,041 | 0 | 4,47E-68 | Plasma B | IGLV10-54 |
| AL133467.1 | 1,9E-72 | 7,31849007 | 0,041 | 0 | 4,63E-68 | Plasma B | AL133467.1 |

**Supplementary table 3.** Comparison of RNA velocity metrics across major immune cell populations in BCRL, LSAT and OSAT adipose tissue.

BCRL= Breast Cancer Related Lymphedema

LSAT= Lean Subcutaneous Adipose Tissue. Selected as control or reference condition

OSAT= Obese Subcutaneous Adipose Tissue

| Comparison | Cell group | Metric | Mean_Target | Mean_Ref | P-Value | P-Adj |
|---|---|---|---|---|---|---|
| **BCRL_vs_LSAT** | B cells | velocity_length | 19.69742857142857 | 21.857470588235294 | 1.6451945425380892e-17 | 3.785403372211532e-17 |
| **BCRL_vs_LSAT** | B cells | velocity_confidence | 0.865063081682308 | 0.9052971608582683 | 1.2490599916229916e-16 | 2.8239617201911114e-16 |
| **BCRL_vs_LSAT** | B cells | latent_time | 0.618277735172645 | 0.566185720761974 | 4.566898655283106e-12 | 9.575755244948448e-12 |
| **BCRL_vs_LSAT** | Dendritic cells | velocity_length | 18.157094339622642 | 19.600227272727274 | 1.7145314912672394e-07 | 2.8946635566849494e-07 |
| **BCRL_vs_LSAT** | Dendritic cells | velocity_confidence | 0.8487993072019203 | 0.8794900146274047 | 0.2646434954934189 | 0.29786713778480056 |
| **BCRL_vs_LSAT** | Dendritic cells | latent_time | 0.6788203523232879 | 0.5011628624076312 | 7.206711260594983e-27 | 2.2850547899447507e-26 |
| **BCRL_vs_LSAT** | Fibroblasts | velocity_length | 19.484107648725214 | 20.776756756756757 | 0.007718983959350261 | 0.01008510467050788 |
| **BCRL_vs_LSAT** | Fibroblasts | velocity_confidence | 0.8336571247838032 | 0.8902159844323749 | 4.216417001155208e-06 | 6.448637766472671e-06 |
| **BCRL_vs_LSAT** | Fibroblasts | latent_time | 0.7947786277453786 | 0.7816918342867992 | 0.0032635160384154923 | 0.004396446476621907 |
| **BCRL_vs_LSAT** | Innate lymphoid cells | velocity_length | 22.345579868708974 | 21.81745098039216 | 0.9186903071324417 | 0.9186903071324417 |
| **BCRL_vs_LSAT** | Innate lymphoid cells | velocity_confidence | 0.8265786974368277 | 0.8703947446192369 | 0.00011289574006190853 | 0.00015866428333024982 |
| **BCRL_vs_LSAT** | Innate lymphoid cells | latent_time | 0.41681981360141235 | 0.3653762755358181 | 1.3543584623480613e-06 | 2.134140607336339e-06 |
| **BCRL_vs_LSAT** | Monocytes | velocity_length | 18.517712975098295 | 20.6109833857207 | 3.526805242469344e-107 | 1.9103528396708947e-106 |
| **BCRL_vs_LSAT** | Monocytes | velocity_confidence | 0.8012501533485608 | 0.8909853031542655 | 7.759944826425024e-260 | 6.725285516235021e-259 |
| **BCRL_vs_LSAT** | Monocytes | latent_time | 0.47981522473328864 | 0.34072623032405125 | 7.033131601180386e-200 | 4.812142674491842e-199 |
| **BCRL_vs_LSAT** | T/NK cells | velocity_length | 24.231457006369425 | 25.35486781811773 | 3.5133331488160405e-05 | 5.019047355451487e-05 |
| **BCRL_vs_LSAT** | T/NK cells | velocity_confidence | 0.8922677580166385 | 0.9422863972437676 | 4.931415686513836e-231 | 3.561577995815548e-230 |
| **BCRL_vs_LSAT** | T/NK cells | latent_time | 0.44181607221094144 | 0.4194824003200633 | 4.2883564065415154e-17 | 9.780461979831526e-17 |
| **BCRL_vs_LSAT** | NK cells | velocity_length | 25.712570150467666 | 28.091448754174156 | 8.40100932632433e-74 | 3.765969698007458e-73 |
| **BCRL_vs_LSAT** | NK cells | velocity_confidence | 0.8862450710825549 | 0.9534648123893398 | 0.0 | 0.0 |
| **BCRL_vs_LSAT** | NK cells | latent_time | 0.49842168239585355 | 0.47492939994653227 | 4.730020008008613e-38 | 1.6846646603866292e-37 |
| **BCRL_vs_LSAT** | Specialized T cells | velocity_length | 19.947533552042948 | 22.242536728697353 | 3.7165818645707146e-49 | 1.46410800725513e-48 |
| **BCRL_vs_LSAT** | Specialized T cells | velocity_confidence | 0.8532298971741756 | 0.9088264293860085 | 3.015805675267185e-166 | 1.8669273227844478e-165 |
| **BCRL_vs_LSAT** | Specialized T cells | latent_time | 0.469202289721225 | 0.3691843263129098 | 5.089161289899733e-302 | 4.725649769192609e-301 |
| **BCRL_vs_LSAT** | CD4+ T cells | velocity_length | 20.63328237967362 | 22.749027613624154 | 1.279391428840481e-175 | 8.316044287463126e-175 |
| **BCRL_vs_LSAT** | CD4+ T cells | velocity_confidence | 0.8445366564251812 | 0.9203705972273974 | 0.0 | 0.0 |
| **BCRL_vs_LSAT** | CD4+ T cells | latent_time | 0.4456005459616989 | 0.3992346683497997 | 7.208191516078383e-104 | 3.8247546820007747e-103 |
| **BCRL_vs_LSAT** | CD8+T cells | velocity_length | 22.22800627943485 | 22.93898590840463 | 0.0031814799823140128 | 0.004308254142716892 |
| **BCRL_vs_LSAT** | CD8+T cells | velocity_confidence | 0.851646079321915 | 0.9120328666326092 | 1.12945929471676e-309 | 1.1294592947167597e-308 |
| **BCRL_vs_LSAT** | CD8+T cells | latent_time | 0.39665756822566145 | 0.36658996858281917 | 8.548679826260347e-09 | 1.5990336365666837e-08 |
| **OSAT_vs_LSAT** | B cells | velocity_length | 21.686246056782334 | 21.857470588235294 | 0.6961105060867098 | 0.7125540613486006 |
| **OSAT_vs_LSAT** | B cells | velocity_confidence | 0.9120251824957027 | 0.9052971608582683 | 0.037077154778444824 | 0.04612469015500313 |
| **OSAT_vs_LSAT** | B cells | latent_time | 0.5602093115876321 | 0.566185720761974 | 0.003057413136062344 | 0.004161923640713138 |
| **OSAT_vs_LSAT** | Dendritic cells | velocity_length | 19.67513595166163 | 19.600227272727274 | 0.7407748759008009 | 0.7523494833367509 |
| **OSAT_vs_LSAT** | Dendritic cells | velocity_confidence | 0.8770827390661639 | 0.8794900146274047 | 0.4646090741361967 | 0.5033264969808797 |
| **OSAT_vs_LSAT** | Dendritic cells | latent_time | 0.5053148731130207 | 0.5011628624076312 | 0.6534874467121018 | 0.6769192675105437 |
| **OSAT_vs_LSAT** | Fibroblasts | velocity_length | 19.927500000000002 | 20.776756756756757 | 0.23097156505346994 | 0.26223845813931085 |
| **OSAT_vs_LSAT** | Fibroblasts | velocity_confidence | 0.8115423067923269 | 0.8902159844323749 | 2.059751072079893e-05 | 2.9587584460816146e-05 |
| **OSAT_vs_LSAT** | Fibroblasts | latent_time | 0.6882373069047728 | 0.7816918342867992 | 0.008202058886357937 | 0.010662676552265316 |
| **OSAT_vs_LSAT** | Innate lymphoid cells | velocity_length | 21.801802325581395 | 21.81745098039216 | 0.5841111890790899 | 0.6198730986145443 |
| **OSAT_vs_LSAT** | Innate lymphoid cells | velocity_confidence | 0.8805996378501848 | 0.8703947446192369 | 0.6098315986870807 | 0.6367719504363092 |
| **OSAT_vs_LSAT** | Innate lymphoid cells | latent_time | 0.36974755245404684 | 0.3653762755358181 | 0.3198921460175339 | 0.3539232253811013 |
| **OSAT_vs_LSAT** | Monocytes | velocity_length | 21.002942697113316 | 20.6109833857207 | 1.49080211280256e-07 | 2.5500562455833263e-07 |
| **OSAT_vs_LSAT** | Monocytes | velocity_confidence | 0.8972989801172201 | 0.8909853031542655 | 2.3329214876120145e-12 | 4.9313787543018194e-12 |
| **OSAT_vs_LSAT** | Monocytes | latent_time | 0.3383226368582533 | 0.34072623032405125 | 0.05919351280743042 | 0.07191735200902762 |
| **OSAT_vs_LSAT** | T/NK cells | velocity_length | 24.91435915010281 | 25.35486781811773 | 4.026043228407616e-10 | 7.870460446511129e-10 |
| **OSAT_vs_LSAT** | T/NK cells | velocity_confidence | 0.938826412768391 | 0.9422863972437676 | 1.4993178414443238e-05 | 2.1900148245815966e-05 |
| **OSAT_vs_LSAT** | T/NK cells | latent_time | 0.3927840629941915 | 0.4194824003200633 | 1.2866657801206985e-29 | 4.288885933735662e-29 |
| **OSAT_vs_LSAT** | NK cells | velocity_length | 25.765384295803905 | 28.091448754174156 | 5.111277545636119e-124 | 2.953182581923091e-123 |
| **OSAT_vs_LSAT** | NK cells | velocity_confidence | 0.9455734432967594 | 0.9534648123893398 | 5.852034545035364e-24 | 1.6360526685045104e-23 |
| **OSAT_vs_LSAT** | NK cells | latent_time | 0.4585916544302366 | 0.47492939994653227 | 3.727946584725004e-09 | 7.074935124295628e-09 |
| **OSAT_vs_LSAT** | Specialized T cells | velocity_length | 22.15116909522196 | 22.242536728697353 | 0.5262671985726083 | 0.5677571436882911 |
| **OSAT_vs_LSAT** | Specialized T cells | velocity_confidence | 0.9152988245614146 | 0.9088264293860085 | 1.6180968765207552e-07 | 2.7497071104274273e-07 |
| **OSAT_vs_LSAT** | Specialized T cells | latent_time | 0.37894041833032727 | 0.3691843263129098 | 0.1462830353403965 | 0.17209768863576058 |
| **OSAT_vs_LSAT** | CD4+ T cells | velocity_length | 22.496183669049167 | 22.749027613624154 | 1.8167863527212466e-25 | 5.557228843617931e-25 |
| **OSAT_vs_LSAT** | CD4+ T cells | velocity_confidence | 0.9172101567296246 | 0.9203705972273974 | 1.6800738086267405e-16 | 3.765682674508211e-16 |
| **OSAT_vs_LSAT** | CD4+ T cells | latent_time | 0.37259185648625853 | 0.3992346683497997 | 0.0 | 0.0 |
| **OSAT_vs_LSAT** | CD8+T cells | velocity_length | 22.80806888970319 | 22.93898590840463 | 0.0029385152546504965 | 0.004021126137942785 |
| **OSAT_vs_LSAT** | CD8+T cells | velocity_confidence | 0.9128471392844985 | 0.9120328666326092 | 0.1501122627785779 | 0.1758071546055417 |
| **OSAT_vs_LSAT** | CD8+T cells | latent_time | 0.3536768743895096 | 0.36658996858281917 | 3.2426038784754567e-56 | 1.3598016264574495e-55 |

Mean_Target and Mean_Ref indicate the average metric values for the target and reference conditions, respectively.

P-values were adjusted for multiple testing using the Benjamini-Hochberg method.

#### Supplementary figure 1

##### A. Heatmap: contribution of each sample to the overall cell types

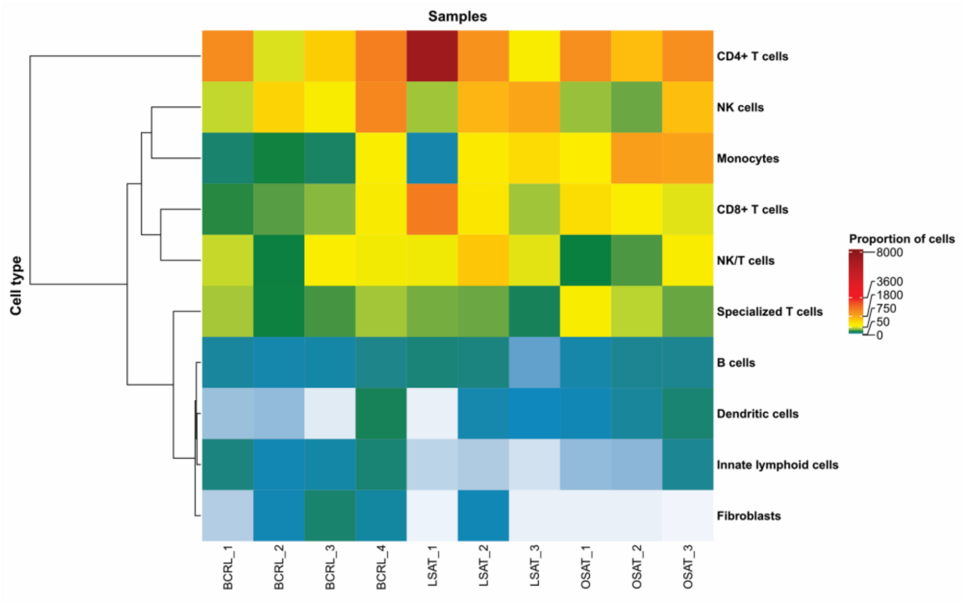


##### B. Distribution of cells from BCRL, LSAT and OSAT within the integrated UMAP embedding

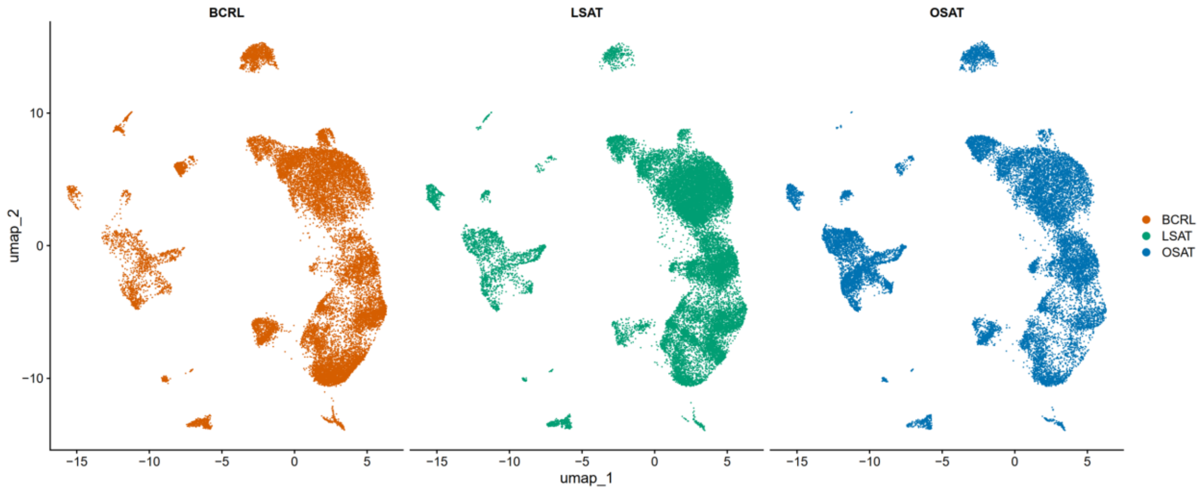

### Supplementary figure 2

A. Differential gene expression profile of Treg cells in BCRL

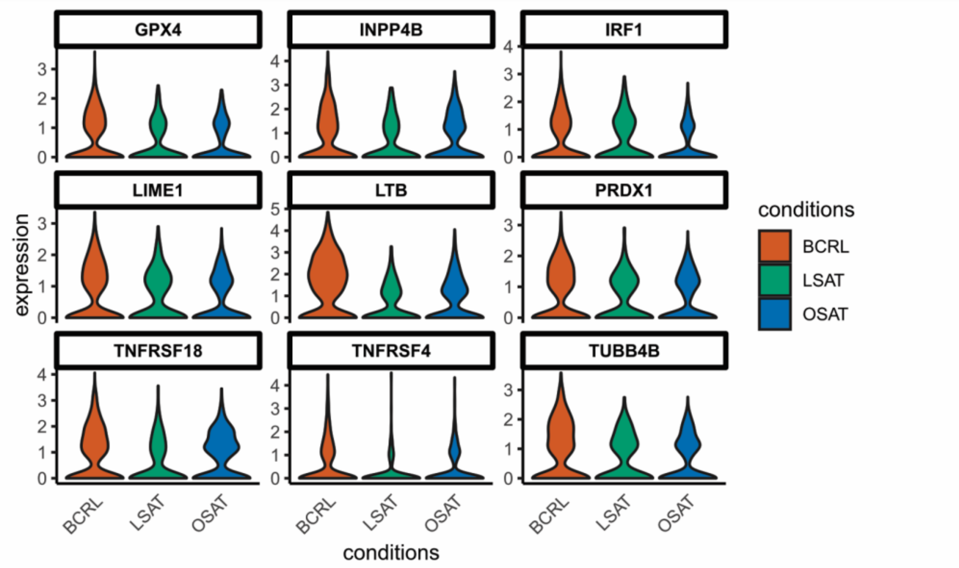


B. Differential gene expression profile of all Dendritic cells in BCRL

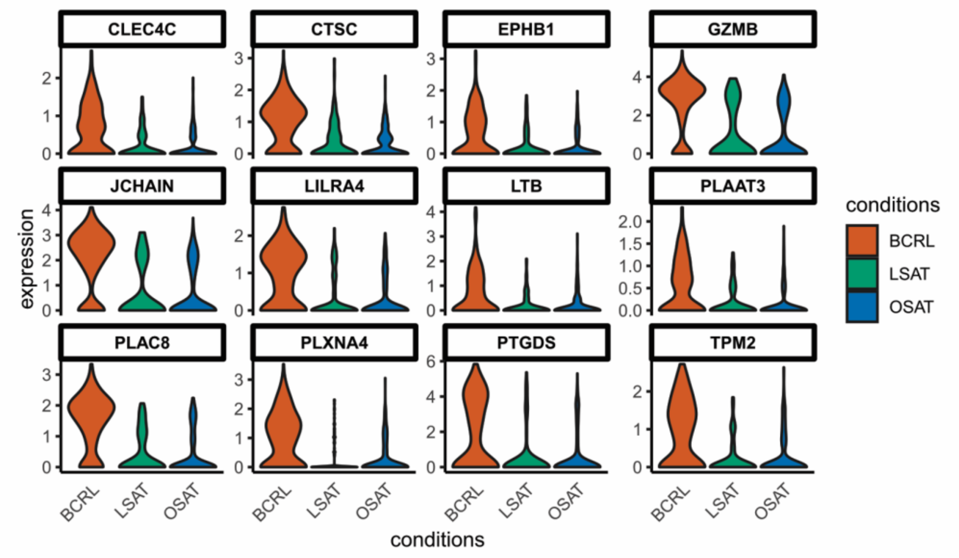

#### Supplementary figure 3

**A. UMAP plots from LSAT, BCRL and OSAT colored by latent-time inferred by scVelo**

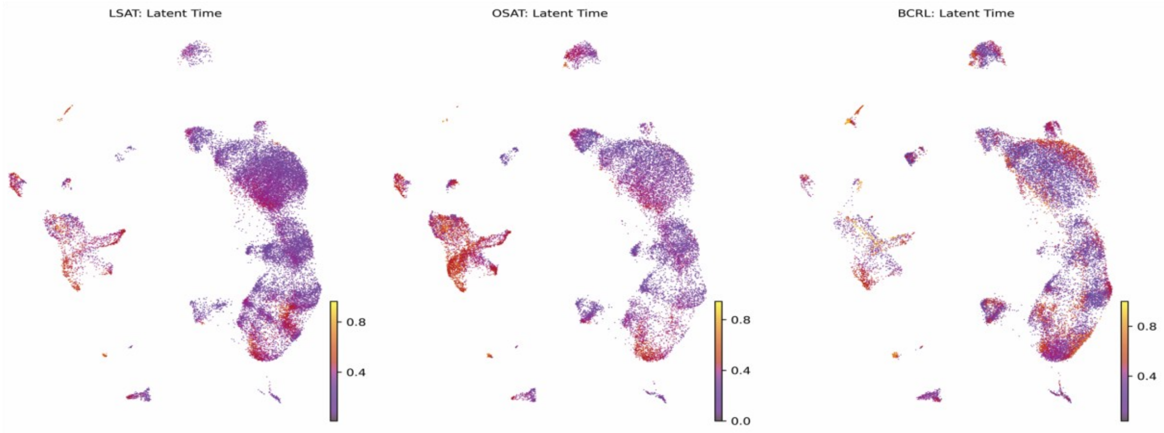


**B. Latent-time distributions in LSAT, BCRL and OSAT tissues per cell group**

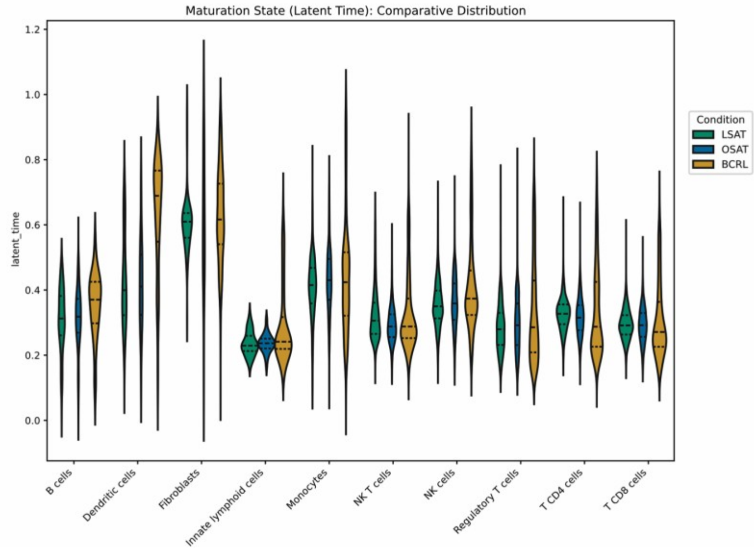

**Supplementary figure 4**

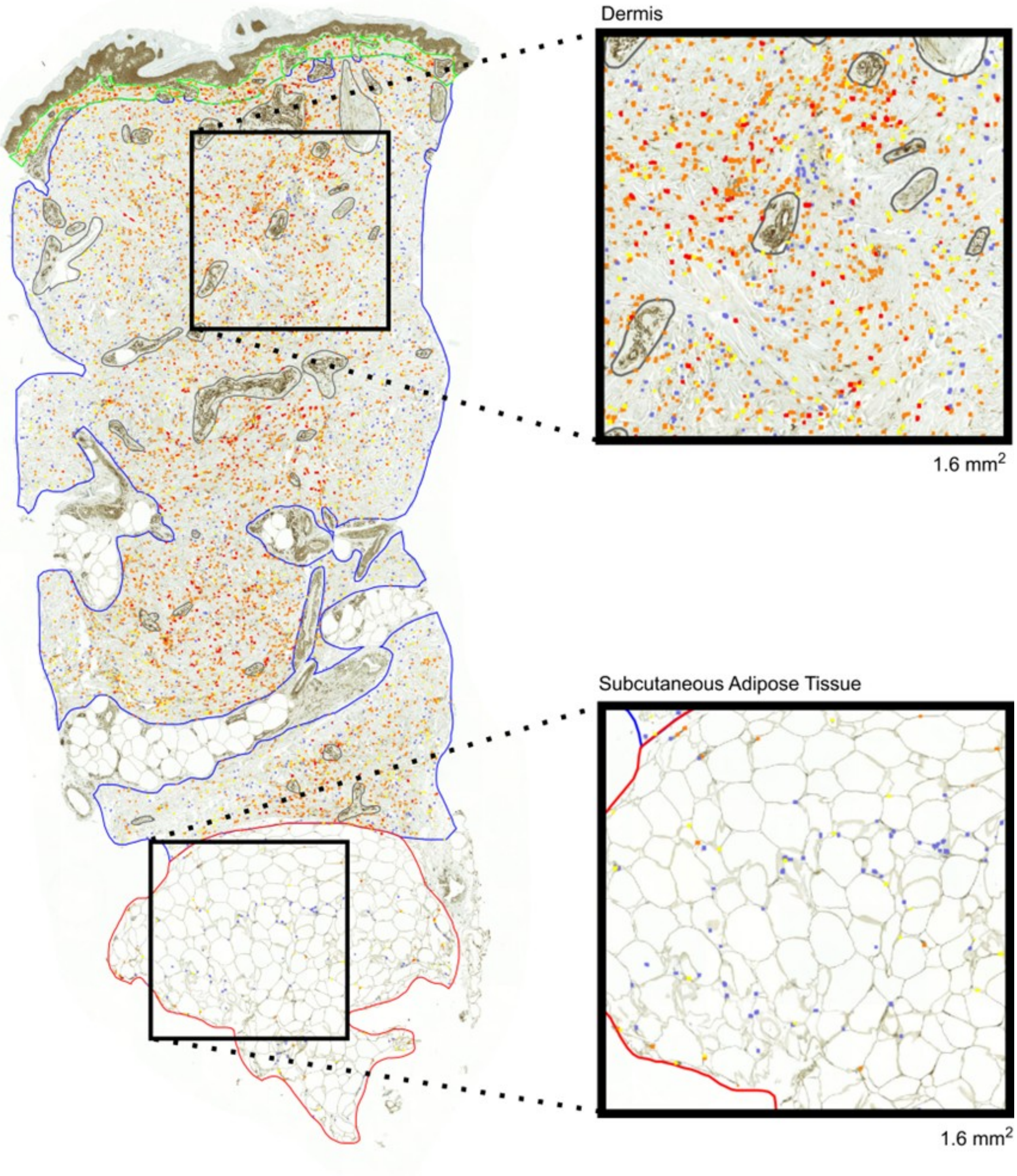
